# Tumor cell total mRNA expression shapes the molecular and clinical phenotype of cancer

**DOI:** 10.1101/2020.09.30.306795

**Authors:** Shaolong Cao, Jennifer R. Wang, Shuangxi Ji, Peng Yang, Matthew D. Montierth, Shuai Guo, John Paul Shen, Xiao Zhao, Jingxiao Chen, Jaewon James Lee, Paola A Guerrero, Nicholas Spetsieris, Nikolai Engedal, Sinja Taavitsainen, Kaixian Yu, Julie Livingstone, Vinayak Bhandari, Shawna M Hubert, Najat C. Daw, P. Andrew Futreal, Eleni Efstathiou, Bora Lim, Andrea Viale, Jianjun Zhang, Matti Nykter, Bogdan A Czerniak, Pavlos Msaouel, Anirban Maitra, Scott Kopetz, Peter Campbell, Terence P. Speed, Paul C. Boutros, Hongtu Zhu, Alfonso Urbanucci, Jonas Demeulemeester, Peter Van Loo, Wenyi Wang

## Abstract

Cancers can vary greatly in their transcriptomes. In contrast to alterations in specific genes or pathways, differences in tumor cell total mRNA content have not been comprehensively assessed. Technical and analytical challenges have impeded examination of total mRNA expression at scale across cancers. To address this, we developed a model for quantifying tumor-specific total mRNA expression (TmS) from bulk sequencing data, which performs transcriptomic deconvolution while adjusting for mixed genomes. We used single-cell RNA sequencing data to demonstrate total mRNA expression as a feature of tumor phenotype. We estimated and validated TmS in 5,015 patients across 15 cancer types identifying significant inter-individual variability. At a pan-cancer level, high TmS is associated with increased risk of disease progression and death. Cancer type-specific patterns of genetic alterations, intra-tumor genetic heterogeneity, as well as pan-cancer trends in metabolic dysregulation and hypoxia contribute to TmS. Taken together, our results suggest that measuring cell-type specific total mRNA expression offers a broader perspective of tracking cancer transcriptomes, which has important biological and clinical implications.

## Introduction

Reprogramming of the transcriptional landscape is a critical hallmark of cancer, which accompanies cancer progression, metastasis and resistance to treatment^1,2^. Recent single-cell studies revealed expansion of cell state heterogeneity in cancer cells that arise largely independently of genetic variation^3–5^, bringing new insights into long-standing topics of cancer cell plasticity^6^ and cancer stem cells^7^. Assessing these clinically relevant topics^8,9^ in large patient cohorts, however, has been difficult due to the high cost and sample quality requirements associated with most single-cell technologies. As bulk tumor RNA and DNA sequencing data are already obtained from large patient series with clinical outcomes, *in silico* approaches to analyze human tissues may expedite our understanding of tumor heterogeneity. We thus aim to develop such an approach.

Previous approaches to build cellular differentiation hierarchies are not suitable for *in vivo* human tissue studies and further require known cell type-specific genetic markers^10^. Variation in total mRNA amount, *i.e*., the sum of detectable mRNA transcripts across all genes per cell, has been linked indirectly to cancer progression and de-differentiation as a result of *MYC* activation^11,12^ or aneuploidy^13,14^. Single-cell studies recently demonstrated that the total number of expressed genes per cell is more predictive of cellular phenotype than alterations in any specific genes or pathways^15,16^. With current limitations in our knowledge of marker genes across cancers, total mRNA expression per tumor cell may represent a robust and measurable pan-cancer feature that warrants a systematic evaluation in patient cohorts.

Total tumor-cell mRNA expression information is masked during standard bulk data analysis, thus requiring deconvolution. Variation in total mRNA transcript levels is removed by routine normalization, together with technical biases such as read depth and library preparation^17–20^. Data generated from cancer studies contain reads from both tumor and admixed normal cells. Copy number aberrations such as gain or loss of chromosomal copies (*i.e*., ploidy) in tumor cells affect gene expression through dosage effect^14^.

Here, building upon prior work in bulk transcriptome deconvolution^21–23^ and in modelling tumor ploidy^24,25^, we created a measure of tumor-specific total mRNA expression (TmS), which captures the ratio of total mRNA expression per haploid genome in tumor cells versus surrounding non-tumor cells. We first scrutinized total mRNA expression using single-cell data across four cancer types^26–28^, and then calculated TmS in matching bulk RNA and DNA data from 5,015 patients across 15 cancer types from TCGA, ICGC^29^, and at MD Anderson Cancer Center^30^ (NCT01946165). Our analyses revealed that variation in total mRNA expression is a feature that captures cancer cell plasticity at a patient-sample level and predicts prognosis across cancers.

## Results

### Diversity in total mRNA expression as a hallmark of cancer cells

To motivate a model-based quantification of total mRNA expression in bulk tissue, we first studied singlecell RNA sequencing (scRNAseq) data generated from 46,468 cells in human colorectal, liver^27^, lung^26^ and pancreatic tumors^28^ (**Fig. 1a, Supplementary Fig. 1a; Methods, Supplementary Note 1.1**).

**Figure. 1:**
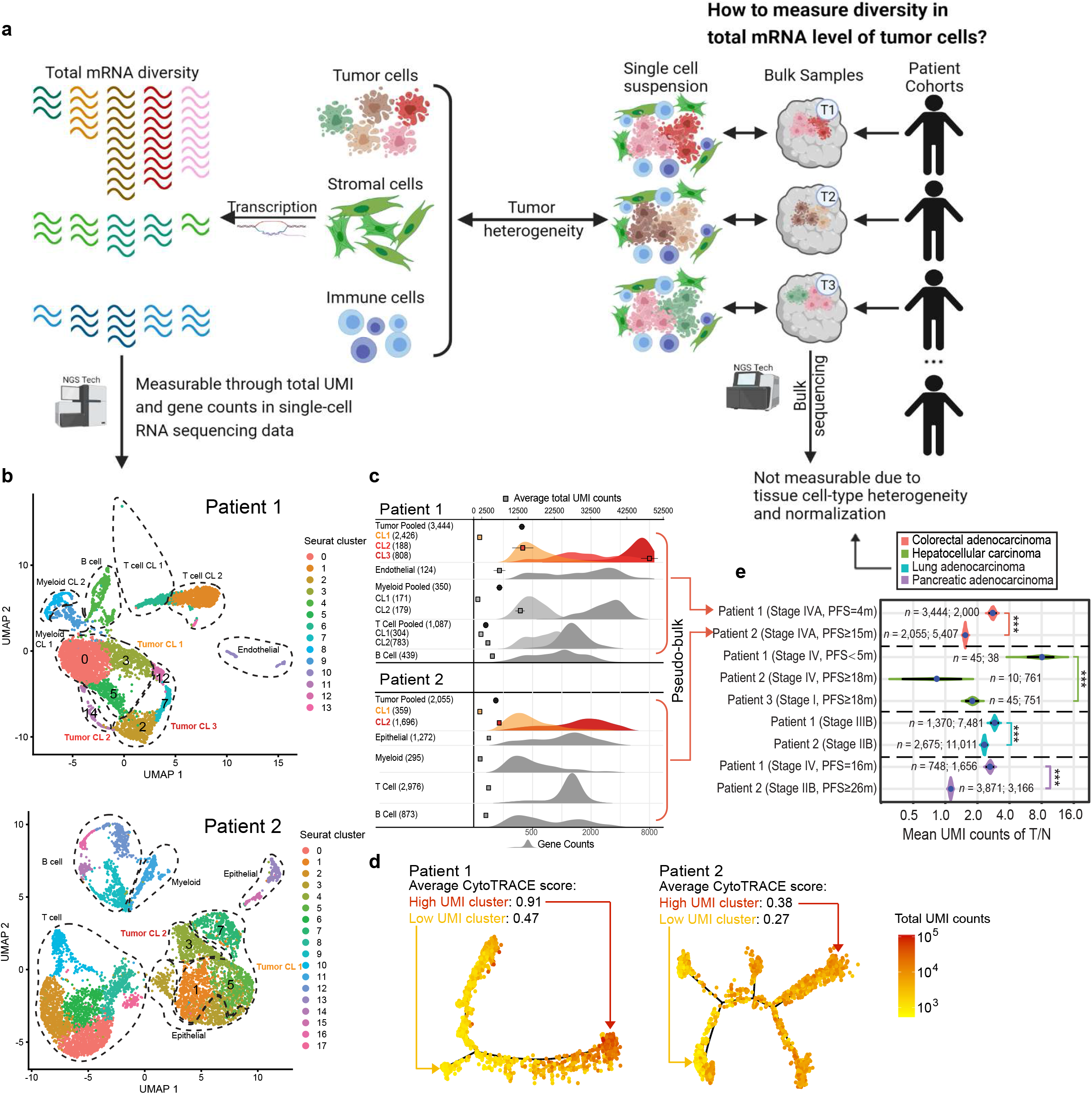
High diversity of total mRNA expression in cancer cells. **a**, Illustration of diversity in total mRNA levels in tumor cells versus other cell types. **b,** UMAP plots of scRNAseq data from two patients with colorectal cancer (CRC). Tumor cell clusters are highlighted in both samples. Dashed lines indicate clusters that are similar in total UMI and gene counts, which are merged for simplicity. **c,** Distributions of gene counts and total UMI counts by cell type in scRNAseq data from the two patients in **b**. The top x-axis annotates total UMI counts (means and 95% Confidence Intervals, CIs). The bottom x-axis annotates gene count distribution (density). Density curves are colored for tumor cells and shown in grayscale for non-tumor cells. Clusters with higher gene counts are shown in darker shades. Numbers in the parentheses indicate the number of cells analyzed. Tumor cell clusters are highlighted by the same colors as those in **b**. **d,** Monocle-inferred trajectories for tumor cells from the two patients. Cells on the trees are colored by total UMI counts. Average differentiation scores by CytoTRACE for high and low UMI clusters are provided. **e,** Ratios of mean total UMI counts of tumor cells (T) to non-tumor cells (N) and 95% confidence intervals in pooled scRNAseq data (pseudo-bulk) from nine patients with colorectal (patients 1, 2 in **b**-**d**), hepatocellular, lung or pancreatic cancer. The number of cells used in the calculation is denoted by *n*: number of tumor cells; number of non-tumor cells. Patients are ordered by pathological stage or survival outcome. The BH adjusted *P* values for Wilcoxon rank-sum tests comparing the ratios between patient samples are indicated by asterisks (* *P* < 0.05, ** < 0.01, *** < 0.001).

Total unique molecular identifier (UMI) counts can be modelled as cell size (total RNA molecule counts) multiplied by cell capture efficiency^31^. We find total UMI counts are highly correlated with gene counts, the total number of expressed genes per cell, across all cells and cell types (median Spearman *r* = 0.96, median absolute deviation (MAD) = 0.03, **Supplementary Note 1.2**). This suggests a similar utility of total UMIs as gene counts for annotating cellular phenotype^15^, where technological effects such as capture efficiency^32^ are nuisance parameters largely constant across cells (**Supplementary Note 1.2**). Expected as a cancer hallmark, we observe larger variability in total UMI and gene counts in tumor cells compared to non-tumor cells (epithelial, stromal and immune cells) within individual experiments where mRNAs from all cells were generated using the same library (F-test for variance *p* values < 0.02, **Supplementary Fig. 1b**). Consistent with recent reports from both human and mouse tumors^26,33^, we find multiple Seurat^34^ clusters within tumor and non-tumor cells presenting distinct total UMI and gene counts (**Fig. 1b-c, Supplementary Fig. 2a; Methods, Supplementary Note 1.2**). Trajectory inference using Monocle^35–37^ shows distinct gene expression states among these clusters (**Fig. 1d**, **Supplementary Fig. 2b**). High-UMI tumor cells present a less differentiated state^15^ (adjusted *P* values < 0.001, **Fig. 1d**, **Supplementary Fig. 2b**; **Methods**), together with a lower or similar cell cycle state. Hence, UMI count is not a surrogate measure for proliferation (**Supplementary Fig. 2c**). In four patients with a shorter time-to-disease progression (colon, liver and pancreas cancers), or advanced-stage disease (lung cancer), the high-UMI tumor cell clusters present differentiation scores close to 1, indicating a stem-like cell state (**Fig. 1c-d, Supplementary Fig. 2b**). This is further supported by their enrichment of stemness and the epithelial-mesenchymal-transition (EMT) gene sets across cancers, out of 18,617 gene sets^38,39^ investigated (**Supplementary Table 2; Methods**). Our observations confirm the utility of total UMI counts to measure a feature of cancer cell plasticity.

When normalized and pooled across cells (pseudo-bulk, **Methods**), increased average total UMI counts for tumor *vs*. non-tumor cells are observed in these four patient samples, as compared to other samples within each cancer type (**Fig. 1e**). Given that the observed differences in average total UMI counts are large and variable across patients, we hypothesize that a quantification of average tumor-specific total mRNA expression in bulk sequencing data is useful to track tumor phenotype and clinical behavior.

### Estimating tumor-specific total mRNA expression from bulk sequencing data

In order to quantify tumor-specific total RNA expression across a large number of patient samples, we employ three steps in a sequential deconvolution of matched DNA/RNAseq data (**Fig. 2a; Methods, Supplementary Note 2.1**). 1) We estimate the ratio of total RNA expression between two cellular populations, tumor versus non-tumor cells, to cancel out technical effects. This ratio can be estimated as an odds of transcript proportions (*π*), based on an optimized set of robust intrinsic tumor signature genes. 2) We divide the total RNA expression by their relative cell fractions to calculate a per-cell total RNA content for tumor and non-tumor cells separately. This step requires matched DNAseq data from which the tumor cell proportion, *i.e*., purity (*ρ*), as well as ploidy (Ψ_*T*_) are estimated. 3) We divide the above metric by ploidy in each component. This step adjusts for the dosage effect of chromosomal copies on gene expression. We thus calculate our final quantitative metric, the per cell per haploid genome total RNA expression for tumor, *i.e*., TmS, as 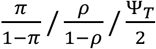. The parameters *ρ* and Ψ_*T*_ can be derived using DNAseq or SNP array data (*e.g*., using a consensus of ASCAT^24^ and ABSOLUTE^25^ or Sequenza^40^, **Supplementary Fig. 3**; **Methods**). The parameter *π* can be derived using RNAseq or microarray data (*e.g*., using DeMixT^23^). Using a profile likelihood of the DeMixT model to rank genes for each study cohort, we identify top-ranked genes as an intrinsic tumor signature gene set, where genes follow a unimodal distribution with low variance across the hidden tumor component and express differentially from the nontumor component (**Supplementary Fig. 4a-b**; **Methods, Supplementary Note 2.2)**. Simulation studies demonstrated improved *π* estimation when only the intrinsic tumor signature genes are used to perform transcriptome deconvolution (**Supplementary Note 2.2)**.

**Figure. 2:**
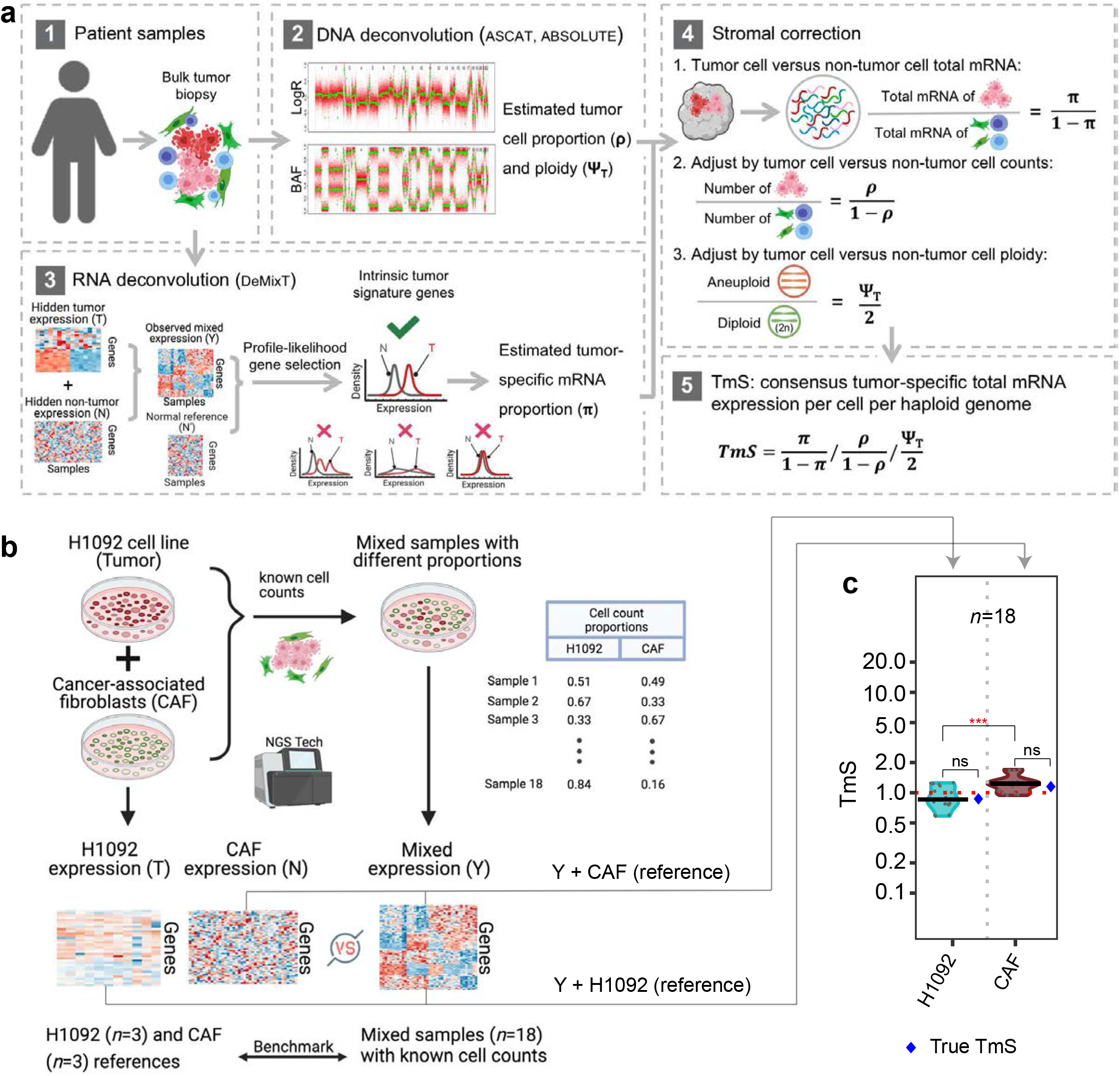
Analysis workflow to measure tumor-specific total mRNA expression. **a**, Calculation of TmS begins with deconvolution using matched DNAseq and RNAseq data. ASCAT and/or ABSOLTE are used to estimate tumor purity and ploidy from the DNAseq data while DeMixT estimates tumor-specific mRNA proportion from the RNAseq. **b**, Benchmarking using bulk RNAseq data from *in vitro* mixed cell lines (H1092 and CAF). **c**, Distribution of TmS in 18 mixed cell-line samples estimated under two scenarios using DeMixT: 1) 3 pure CAF samples as normal reference; 2) 3 pure H1092 samples as normal reference. The true TmS values for H1092 and CAF are provided in blue dots. They are measured as the ratio of total RNA amount (in ng/ul) in 1 million cells: 0.87 for H1092, and 1.2 for CAF. The median estimates of TmS are (0.86, 1.2), with MADs of (0.24, 0.18) for H1092 and CAF cells. The *P* values for Wilcoxon rank-sum tests comparing TmS between groups are indicated (ns: *P* > 0.05, ***: *P* < 0.001).

We benchmarked the performance of TmS estimation using total RNAseq data generated from mixed cell populations with known proportions^23^, resulting in accurate separation of H1092 lung cancer cells from cancer-associated fibroblasts (**Fig. 2b-c, Supplementary Table 3, Supplementary Fig. 3i; Methods**).

We calculated TmS across 15 TCGA cancer types and the early-onset prostate cancer (EOPC) ICGC-EOPC cohort, and found considerable variation across patients. All cancer types demonstrate a much wider TmS range than observed in the benchmarking data which were generated at single TmS values (**Fig. 3, Supplementary Table 4; Methods, Supplementary Note 2.3**). The intrinsic signature genes used in this calculation largely overlap for different cancers, with 2,042 genes selected by at least 1/3 of all cancer types (**Supplementary Fig. 4c**). Across cancers, they are consistently enriched in housekeeping and essential genes^41,42^, and in cancer hallmark^38^ and transcriptional regulation pathways, and are more accessible with open chromatins^43^ (**Supplementary Fig. 4c-e; Methods, Supplementary Note 2.3**). These pan-cancer characteristics support our profile-likelihood based approach to successfully select stably and differentially expressed genes in tumor cells.

**Figure. 3:**
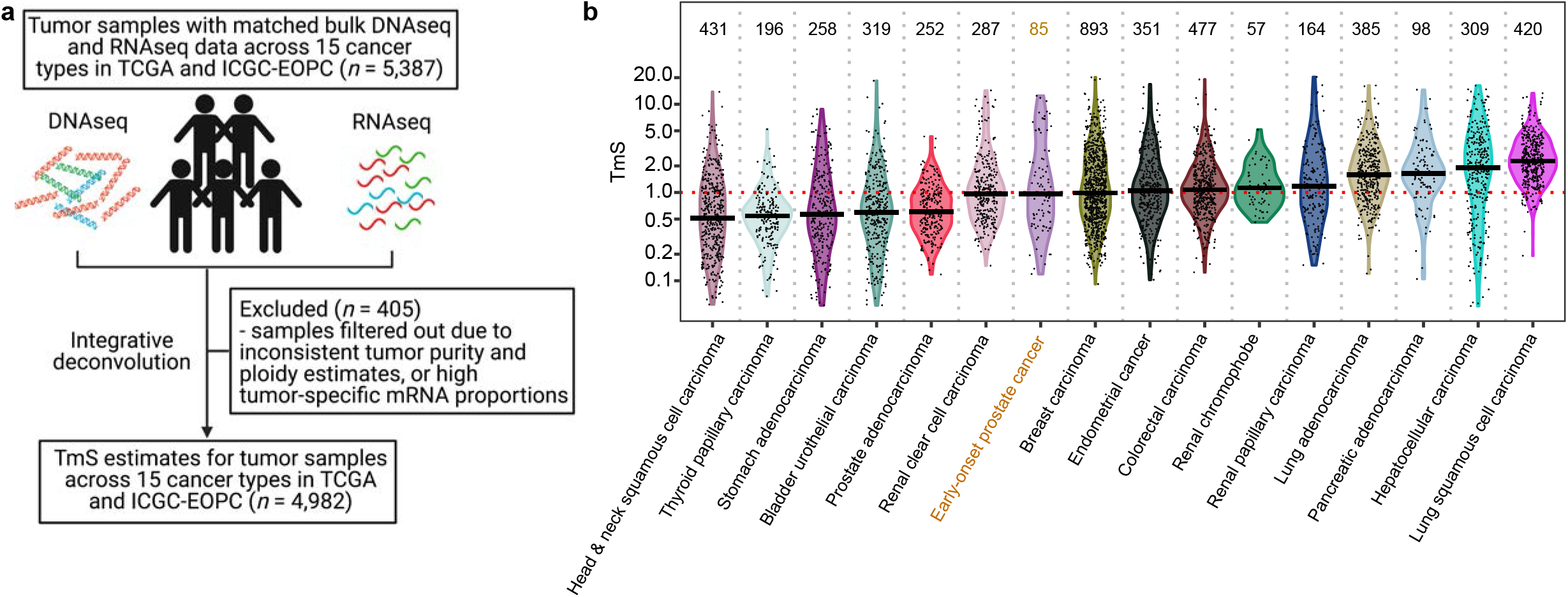
Estimation of tumor-specific total mRNA expression in bulk sequencing data. **a**, CONSORT diagram for the TmS calculation in TCGA and ICGC-EOPC datasets. The number of tumor samples is denoted by *n*. **b**, Distribution of TmS in 4,982 tumor samples across 15 cancer types in TCGA and ICGC-EOPC. The number of tumor samples for each cancer type is indicated above each violin plot.

### TmS is associated with prognostic clinicopathologic characteristics

TmS is dependent upon the background normal reference tissue. For tumors derived from the same tissue, comparisons can be made between known histopathologic and/or molecular subtypes. Consistent trends are observed between subtypes of head-and-neck squamous cell carcinoma, breast carcinoma, renal papillary carcinoma, and prostate adenocarcinoma, where prognostically favorable subtypes are enriched in tumors with lower TmS and *vice versa* (**Fig. 4a-e; Methods**). In head-and-neck squamous cell carcinoma, the prognostically favorable human papillomavirus (HPV)-positive subtype has lower median TmS than the HPV-negative subtype (*P* value = 0.004, **Fig. 4a**). Similarly, triple negative receptor status is associated with higher TmS in breast carcinoma (adjusted *P* value = 5 x 10^-36^, **Fig. 4b**), in keeping with this subtype’s known propensity for aggressive behavior. Subtypes of renal papillary carcinoma also show significant differences in TmS, where the more aggressive Type II tumors have higher TmS compared to Type I (*P* value = 7 x 10^-5^, **Fig. 4c**)^44^. In bladder cancer, increased TmS is associated with the basal molecular subtype, which is known to exhibit more aggressive behavior compared to luminal tumors (*P* value = 2 x 10^-4^, **Fig. 4d**)^45–47^. In prostate adenocarcinoma, TmS is associated with tumor grade as assessed by Gleason score, where high TmS tumors enrich for Gleason scores of 8 and above (*P* value = 0.004, **Fig. 4e**). To further support its unique biological identity, we find that TmS is not a surrogate for Tumor-Node-Metastasis stage, cellular proliferation or other patient characteristics such as age and sex, as is shown by weak to no correlations between TmS and the corresponding variables (**Supplementary Fig. 5a-c**).

**Figure. 4:**
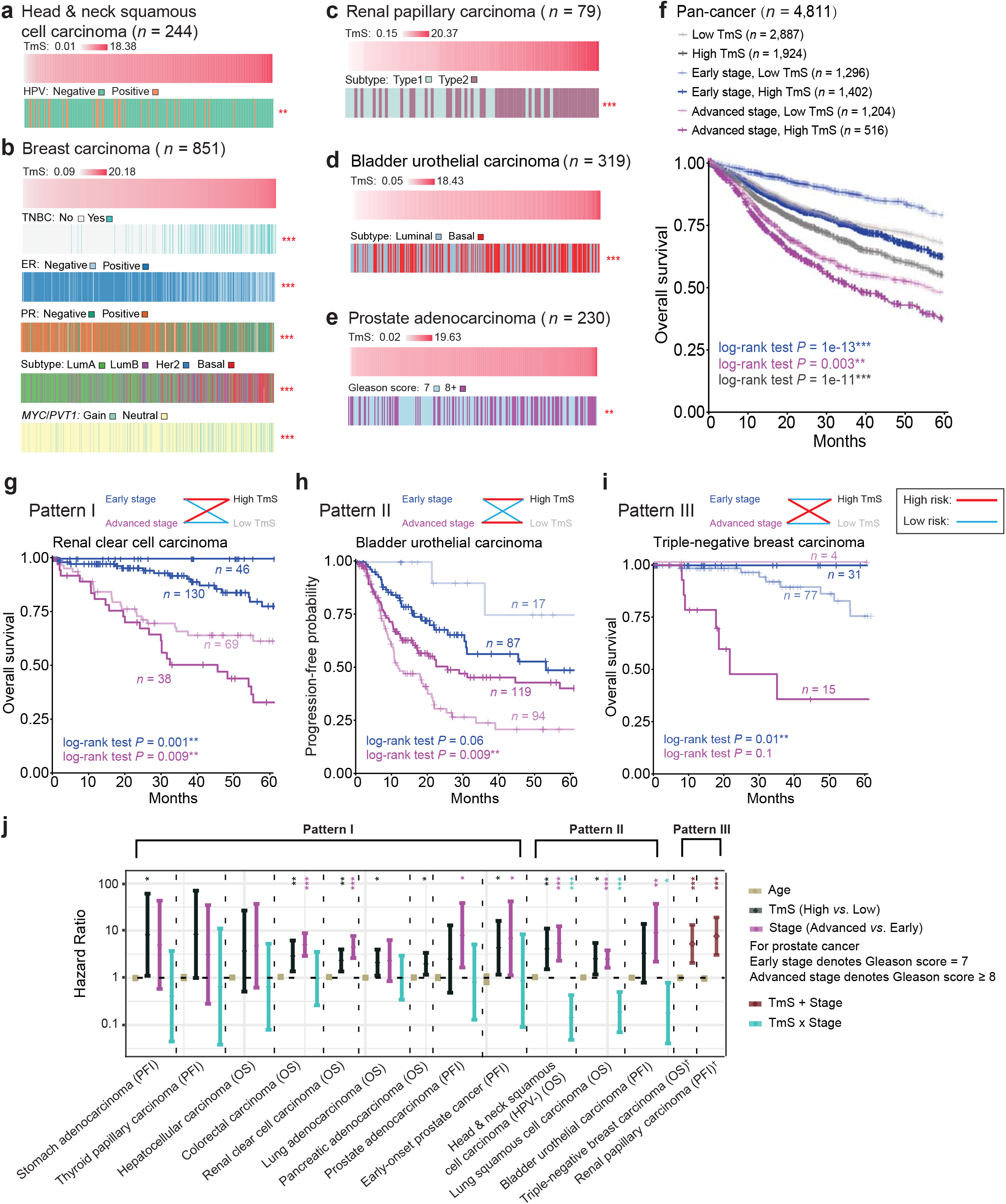
TmS is associated with known prognostic characteristics and refines prognostication in addition to stage classification. **a-e**, Clinicopathologic annotations are shown for (**a**) head & neck squamous cell carcinoma, (**b**) breast carcinoma, (**c**) renal papillary carcinoma, (**d**) bladder urothelial carcinoma, and (**e**) prostate adenocarcinoma samples. Tumor samples are ordered by TmS from low to high. The BH adjusted *P* values for Kruskal-Wallis tests comparing TmS between clinicopathologic subgroups are indicated by asterisks (* *P* < 0.05, ** < 0.01, *** < 0.001). For breast carcinoma, triplenegative breast cancers (TNBC) are shown on the second row. *MYC/PVT1* copy number alterations are shown on the sixth row, where “Gain” indicates that either *MYC* or *PVT1* was amplified and “Neutral” indicates that no copy number alternations were detected. **f**, Kaplan-Meier (KM) curves of overall survival (OS) for TCGA samples. Gray lines denote summary KM curves of patients with high *vs*. low TmS across all cancer types. KM curves are further grouped by TmS and pathological stage into four groups. *P* values of log-rank tests between high *vs*. low TmS groups are indicated by asterisks (* *P* < 0.05, ** < 0.01, *** < 0.001). Three patterns of associations were identified (summarized in **j**). **g-i**, KM curves for representative cancers in Pattern I (**g**), Pattern II (**h**) and Pattern III (**i**). **j**, Forest plot of hazard ratios and 95% of CIs of multivariate Cox proportional hazard models with Age, TmS (High *vs*. Low), Stage (Advanced *vs*. Early) and TmS x Stage as predictors, and with OS or progression-free interval (PFI) as response variable (See details in **Supplementary Table 5**).

### TmS refines prognostication across cancer types in TCGA and ICGC-EOPC

We examined the association of TmS with overall survival (OS) and progression-free interval (PFI) across TCGA cohorts (**Methods, Supplementary Note 3**). In pan-cancer analyses, high TmS is associated with reduced OS and PFI (**Fig. 4f, Supplementary Fig. 5d**). We next examined each cancer type in the context of overall TNM stage classification, which is used across cancers for predicting prognosis and treatment decision-making. Analysis stratified by early (I/II) *vs*. advanced (III/IV) stages reveal three different patterns for the effects of TmS by stage. The first group of tumors show consistent effects across stages (**Fig. 4g; Supplementary Fig. 5e**). This includes thyroid, lung adeno, colorectal, hepatocellular, stomach adeno, and renal clear cell carcinomas, where high TmS is associated with higher risks of death and/or disease progression within both early- and advanced-stage tumors. Prostate adenocarcinoma also belongs to this group, where stage is replaced by Gleason scores 7 *vs*. 8+ in line with clinical practice. In head-and-neck squamous cell, lung squamous cell, and bladder urothelial carcinomas, high TmS is associated with reduced survival in early-stage tumors only, while for advanced-stage tumors, high TmS is associated with improved survival (**Fig. 4h; Supplementary Fig. 5f**). An opposite pattern is observed in breast and renal papillary carcinomas, where high TmS is associated with poor prognosis in advanced-stage tumors, but improved survival in early-stage tumors (**Fig. 4i; Supplementary Fig. 5g)**. TmS remains significantly associated with survival outcomes in Cox regression models, after adjusting for key prognostic characteristics including subtype, stage and age, except in hepatocellular carcinomas and prostate adenocarcinoma (TCGA), where only a trend towards significance was observed (**Fig. 4j, Supplementary Table 5**). Early-onset prostate cancer from ICGC-EOPC shows a significant effect of TmS within Gleason score = 7 (**Fig. 4j, Supplementary Fig. 5e, Supplementary Table 5**). We also find that TmS, which is corrected for tumor ploidy and expressed per haploid genome copy, shows better risk stratification than an analogous measure without ploidy adjustment (**Supplementary Fig. 6**).

### Pan-cancer and cancer-specific contributions to tumor-specific total mRNA expression

We hypothesize that as a feature of tumor phenotype, tumor-specific total mRNA expression should be associated with interacting biological processes (**Fig. 5a**). Differential gene expression analyses comparing high to low TmS samples demonstrate enrichment for metabolic pathways across cancers (**Supplementary Fig. 7a; Methods**). Of the typically seven metabolic pathways of carbohydrates^48^, the pentose phosphate pathway is upregulated in high TmS samples, achieving statistical significance in 12 out of 15 cancer types (**Fig. 5b, Supplementary Fig. 7a-b**). This is a major pathway for nucleotide synthesis through the formation of ribose 5-phosphate. As such, tumors with high TmS may require upregulation of this pathway to provide nucleotides for mRNA synthesis. Moreover, we find a dichotomized hypoxia score^49^, based on 51 genes, is significantly associated with TmS across all 14 cancer types with available data (**Fig. 5c; Methods**). High TmS correlates with high hypoxia in all cancer types except head-and-neck cancers. While there are no overlapping genes between the hypoxia and pentose phosphate pathway gene sets, the hypoxia scores are moderately correlated with pentose phosphate score across cancers (**Supplementary Fig. 7c; Supplementary Note 4**). Network analysis shows dense connectivity between a subnetwork of 61 genes from these two pathways (density score = 0.32^50^, **Supplementary Fig. 7d; Methods**). The consistent and pan-cancer identification of both biological processes supports the molecular relevance of our proposed TmS measurement.

**Figure. 5:**
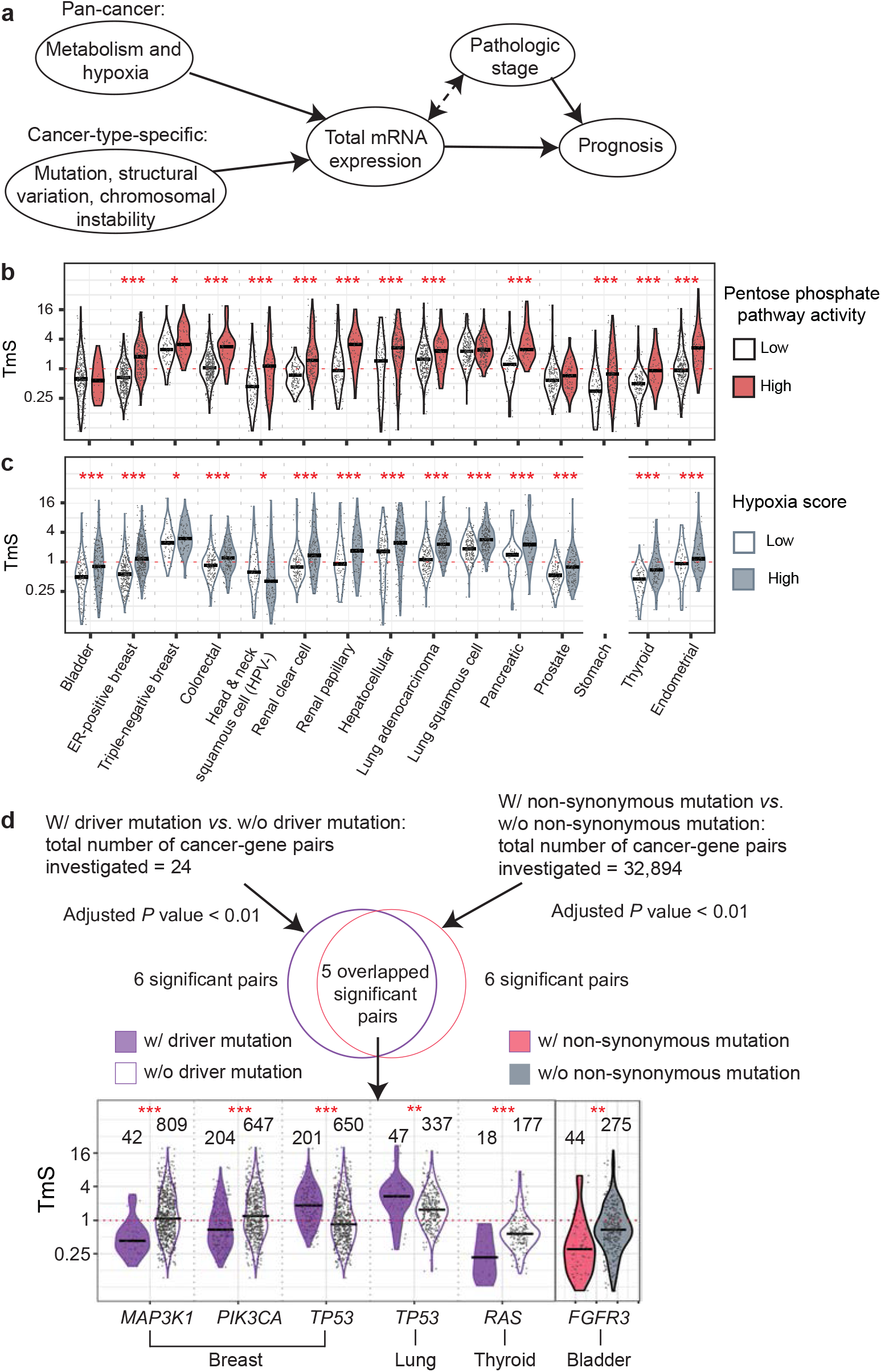
Pan-cancer and cancer-type-specific contributions to tumor-specific total mRNA expression. **a**, Total mRNA expression levels may capture the response of tumor cells to various environmental effects and determine disease trajectory, affecting prognosis. Some of these effects are cancer-type-specific while others may be consistent across cancers. **b-c**, Distributions of TmS for patient samples with (**b**) high or low for pentose phosphate pathway activity, where patient groups are defined by hierarchical clustering of expression levels from the 13 pathway genes (see example in **Supplementary Fig. 7b**); (**c**) high or low hypoxia score, where disease cohorts are divided in half at the median hypoxia scores. **d**, Distributions of TmS for TCGA samples with or without driver mutations in the five significant cancer-gene pairs and one significant cancer-gene pair with or without non-synonymous mutations. The number of samples is indicated on the top.

Interrogation of genetic alterations in TCGA revealed cancer-specific patterns associated with TmS. We performed two interrogations in parallel: an agnostic association analysis of TmS with all non-synonymous mutations (32,894 cancer-gene pairs, using logistic regression models to adjust for covariates such as tumor mutation burden), as well as a driver mutation-specific association analysis of TmS (24 cancer-gene pairs). We find 5 overlapping pairs out of 6 statistically significant pairs produced from each interrogation (**Fig. 5d,** adjusted *P* values < 0.01). The additional pair found through the agnostic search (*FGFR3* in bladder carcinoma) was missed in driver mutation analysis due to a lack of sample size. This suggests TmS can capture changes in tumor phenotypes induced by specific driver mutations^51^.

Furthermore, we examined broad-scale genetic alterations, including tumor mutation burden (TMB), chromosomal instability (CIN), and whole-genome duplication status (WGD). Significant associations are sparsely identified across cancers, suggesting that these features may contribute to tumor-specific total mRNA expression in specific cancer types but are not universal determinants (**Supplementary Fig. 8**).

### Case studies in cancer patients with varying tumor-cell total mRNA expression

Finally, we provide two independent cohort studies to validate that TmS can quantify variations in tumor cell total mRNA expression in response to different biological processes.

Renal medullary carcinoma (RMC) is a rare but highly aggressive disease characterized by very poor prognosis compared with clear cell renal cell carcinoma (KIRC), papillary renal cell carcinoma (KIRP) and chromophobe renal cell carcinoma (KICH), which are the three most common renal cell carcinoma subtypes^52^. All RMC tumors harbor inactivating *SMARCB1* alterations leading to high *MYC* expression and replicative stress^30^. As a result, morphological features consistent with high levels of RNA transcription, such as open phase nuclei with prominent nucleoli, are a hallmark of RMC (**Fig. 6a**). We analyzed matched RNAseq and exome sequencing data from untreated primary RMC tumor samples and adjacent kidney control tissues, derived from 9 patients with clinicopathologic features representative of this highly aggressive disease^30^ (**Methods**). Significantly higher TmS values are presented in RMC than those from KIRC, KIRP and KICH (with the same normal tissue background) in TCGA (**Fig. 6b**), demonstrating this *MYC*-driven renal cell carcinoma is distinguished from other renal cell carcinomas by increased tumor-cell total mRNA expression.

**Figure. 6:**
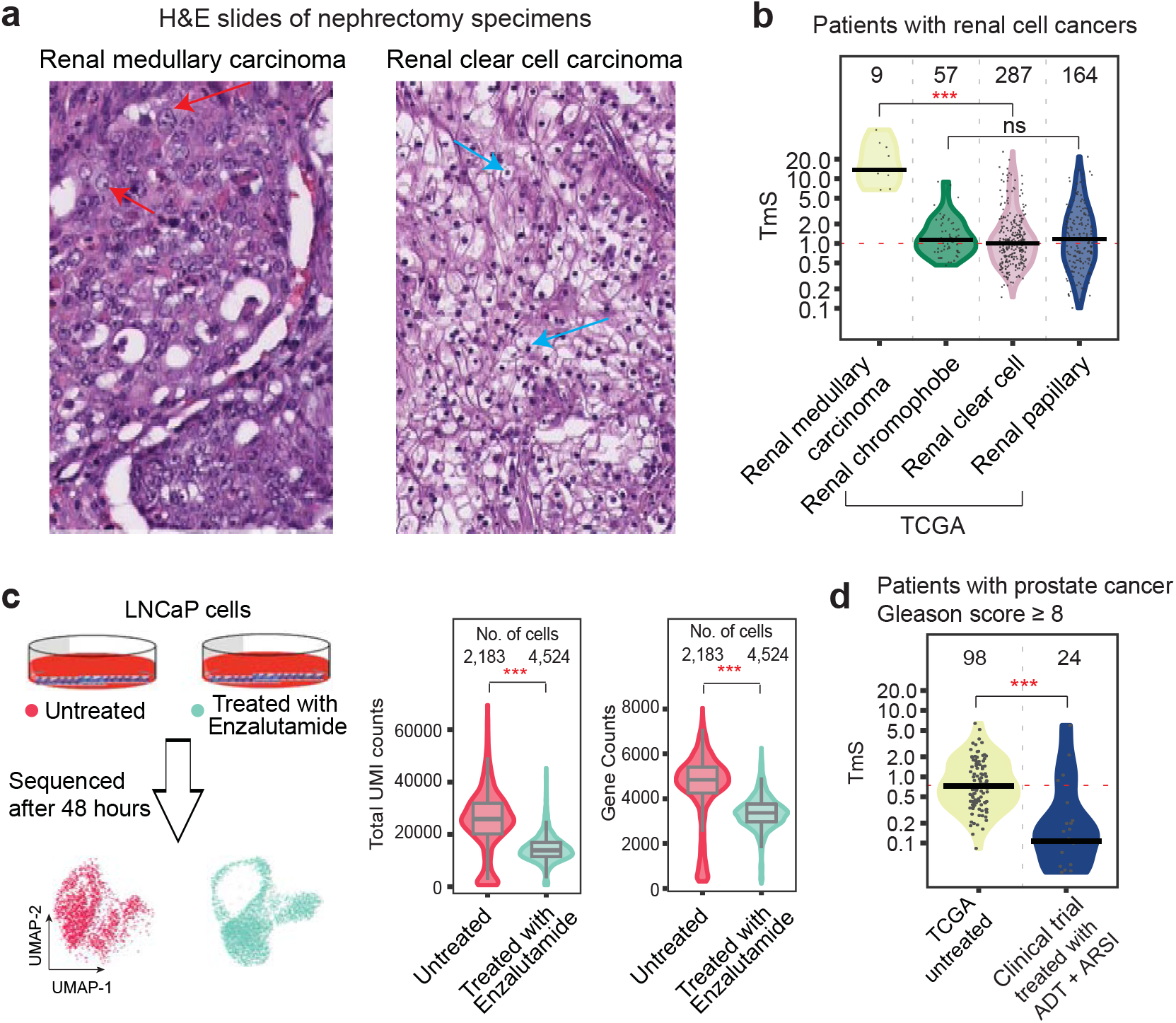
Case studies in cancer patients with varying tumor-cell total mRNA expression. **a**, Microphotographs of H&E slides of nephrectomy specimens from a patient with renal medullary carcinoma (left) and a patient with renal clear cell carcinoma (right) (20x magnification). Red arrows point to the prominent nucleoli in RMC as example. Blue arrows point to nuclei in renal clear cell carcinoma for comparison. **b**, Distributions of TmS for RMC samples and three types of renal cancer in TCGA. The *P* values of Wilcoxon rank-sum test comparing TmS between RMC and renal clear cell carcinoma in TCGA are indicated by asterisks (* *P* < 0.05, ** < 0.01, *** < 0.001); ns: not significant. **c**, Illustration of an *in vitro* experiment where the LNCaP cells were treated for 48 hours with DMSO or Enzalutamide and subsequently processed using single-cell RNAseq. These cells were then collected and sent for single-cell RNAseq experiments. The distributions of total UMI counts and gene counts are shown. **d**, Distribution of TmS for tumor samples from prostate cancer patients in TCGA who were untreated, and patients at MD Anderson who underwent treatment with ADT+ARSI. All samples from TCGA presented a Gleason score of 8+, matching the patient population at MDA.

Androgen deprivation therapy (ADT) and enhanced androgen receptor (AR) signaling inhibitors (ARSIs) such as Enzalutamide (Enza) and Abiraterone (Abi) are currently used to treat prostate cancer patients^53,54^. AR activity has been shown to induce chromatin changes which leads to increased transcriptional activity^55–58^. Therefore, we hypothesize that tumor cells that remain after ADT+ARSI treatments will have reduced tumor-cell total mRNA expression levels. First, scRNAseq data from the prostate cancer cell line LNCaP treated with and without Enza for 48 hours (**Methods, Supplementary Note 1**) showed Enza-treatment reduced TmS in these cells (**Fig. 6c**). ARSI agents are used in clinical trials studying their effects in the treatment of high-risk primary prostate cancer (e.g., NCT01946165 at MD Anderson). We find considerably lower TmS in radical prostatectomy tissue specimens from 24 patients with high-risk localized disease (with Gleason score ≥ 8) who were treated with ADT+ARSI for 6 months, as compared to untreated tumor samples from patients (with Gleason score ≥ 8) in TCGA (**Fig. 6d; Methods**), demonstrating that AR signaling inhibition in prostate cancer leads to decreased tumor-cell total mRNA expression.

## Discussion

Our study identifies TmS, a robust and measurable feature of cancer cell plasticity, from bulk tumor tissues that is clinically and molecularly relevant across cancers. While single-cell technology depicts tumor cell populations with distinct gene expression states (a microscopic view), questions remain on how these populations coexist and interchange to impact clinical outcome^6^. Average signals across all tumor cells intrinsically account for the magnitudes and fractions of each population and are demonstrated by this study to be informative for the clinical presentations of cancer (a macroscopic view).

Cancer cell plasticity, previously evaluated within a few tumors or in model systems only^9^, is measured in a pan-cancer panorama (5,015 patient samples from 15 cancer types) by TmS, an integrative RNA and DNA deconvolution metric for bulk tissues. Association of TmS with genomic features, metabolism and hypoxia, supports a consistent and biologically meaningful measurement of a bulk-level feature of tumor phenotype. The ability of TmS to refine prognostication is observed within each of the 13 cancer types where pathological stages are available, which was not achievable by signature genes for EMT (the most studied pathway for plasticity)^59^ alone in bulk tissues^60^.

While high tumor-cell total mRNA expression is generally associated with aggressive disease, clinical context remains important to evaluate its prognostic implications, as the direction of the prognostic effect was inverted by stage in five out of thirteen cancer types. Such complexity is expected and consistent with often contradictory findings of cancer stem cell models across cancer types^6,8^, as the biological underpinnings are complex and largely unknown. Given that early and advanced stage tumors and different tumor types are often treated using distinct modalities, the inverted effect may in part be underpinned by a differential response of tumors with low *vs*. high total mRNA expression to treatment conditions. The prognostication effect of TmS in bladder cancer as pattern II (**Fig. 4h**) is supported by evidence shown in the basal molecular subtype, which, although more aggressive than the luminal subtype and presenting mostly in an advanced stage, is more responsive to frontline cisplatin-based chemotherapy^45,46^. Additional studies incorporating data from clinical trials will be needed to elucidate how stage-specific and treatment-related factors interact with tumor-cell total mRNA expression to determine patient outcome.

Conceptually, analogous to DNA ploidy measuring the total DNA content calculated per haploid genome, the total mRNA content per haploid genome can be considered the “ploidy of the transcriptome”. Total mRNA content is a key parameter of tumor heterogeneity and phenotype plasticity, previously hidden in most RNA-based assays. While our current work focuses on interpretation of mRNA, TmS is validated for total RNA and the methodology developed here can readily be applied to the quantification of other RNA species (*e.g*. rRNA, miRNA, piRNA), further illuminating the cancer transcriptome. Enhanced attention to “transcriptome ploidy” will enable better phenotypic characterization and a deeper biological understanding of transcriptional dysregulation in cancer and other diseases.

## Methods

Additional details and results are described in **Supplementary Note**. Here, we summarize the key aspects of the analysis.

### Total mRNA expression in single-cell RNA sequencing data

#### Dataset

We collected single-cell RNA sequencing (scRNAseq) data from nine patients, comprising 2 with colorectal adenocarcinoma, 3 with hepatocellular carcinoma, 2 with lung adenocarcinoma, and 2 with pancreatic adenocarcinoma (**Supplementary Table 1**), as well as from the LNCaP prostate cancer cell line. A full description is provided in **Supplementary Note, Section 1.1**.

#### Quality control, clustering, cell type annotation, and normalized UMI

For each sample, we first filtered out cells based on number of genes expressed, total UMI (unique molecular identifier) counts, and proportion of total UMI counts derived from mitochondrial genes. We also removed cells that are detected as doublets. After the quality control, the nine patient samples across four cancer types had 46,468 cells remaining, and the two samples from the LNCaP cell line had 6,707 cells remaining, respectively. Within each patient sample, highly variable genes were detected and used for principal component analysis (PCA). Cells were then clustered with the Seurat package^34^. Cell type was annotated using known marker genes^26,27,61–63^. Tumor cells were identified based on the inferred presence of somatic copy number alterations by inferCNV^64^. We further merged Seurat^34^ identified clusters that were not significantly different in gene counts, which is the total number of expressed genes (Wilcoxon rank-sum test, α=0.001, **Fig. 1b**). A full description is provided in **Supplementary Note, Section 1.2.1**.

To enable comparison between different scRNAseq samples within the same study, we performed scale normalization for the total raw UMI counts across all cells, to ensure the total UMI count per cell are comparable for different samples from the same study. A full description is provided in **Supplementary Note, Section 1.2.2**.

#### Trajectory and differentiation score calculation in tumor cells

We applied Monocle 2 (version 2.14.0)^35–37^ to construct single-cell trajectories, and used the CytoTRACE score to measure the differentiation state of tumor cells^15^. For Monocle 2, we ran “differentialGeneTest” (cutoff: q-value < 0.01) to select differentially expressed (DE) genes across tumor cell clusters, then used the “DDR-Tree” method of “reduceDimension” with selected DE genes for dimensionality reduction and tree construction. In order to compare CytoTRACE scores between the tumor cell clusters from patient samples within the same cancer type, we integrated tumor cells from patient 1 and patient 2 from each of the colorectal, lung and pancreatic cancers using ComBat (version 3.20.0)^65^ embedded in CytoTRACE to correct for batch effects.

#### Gene set enrichment analysis in scRNAseq data

We performed gene set enrichment analyses for the high and low UMI tumor cell clusters in scRNAseq data. We first compiled a comprehensive set of signatures with 18,617 human gene sets (containing at least 4 genes) from the Molecular Signatures Database (MSigDB, v6.2)^38^ and CellMarker^39^. Among them, 341 gene sets were annotated as stemness signatures based on the key word ‘_stem_’ in their names. We quantified enrichment for high and low UMI tumor cells using the GeneOverlap R package (v1.24.0)^66^. GeneOverlap took the DE genes, a gene set and the background genome size (number of expressed genes in the scRNAseq data expressed in ≥ 10 cells) as input, and gave a *P* value for the enrichment significance and Jaccard Index, which was calculated by the number of common genes in the DE gene list and the signature gene set divided by the union of them. *P* values were adjusted for multiple comparisons using the Benjamini-Hochberg (BH) correction. The DE genes between high UMI and low tumor cells were obtained by the “FindMarkers” function from Seurat with Wilcoxon rank-sum test, based on criteria of adjusted *P* value < 0.1, genes expressed in ≥ 10 cells, and absolute log_2_(fold change) > 0.585 (1.5 fold change).

#### Pseudo-bulk analysis

We pooled scRNAseq data to form pseudo-bulk samples and estimated the ratio of the mean total UMI counts of tumor cells to that of the non-tumor cells for each sample. The 95% confidence intervals were constructed using bootstrapping of the same number of tumor and non-tumor cells with 1,000 repetitions.

### Tumor-specific total mRNA expression in bulk sequencing data

#### A mathematical model for tumor-specific total mRNA expression estimation

For any group of cells, we use *S* to denote the average global mRNA transcript level per cell per haploid genome, which follows 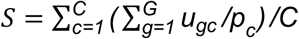. Here *U_gc_* denotes the number of mRNA transcripts of gene *g* in cell *c*, *G* denotes the total number of genes, *C* is the number of cells, *p_c_* is the ploidy, *i.e*., the number of copies of the haploid genome in cell *c*. However, the cell level ploidy *p_c_* is usually not measurable. Hence, in practice, we use average ploidy *Ψ* of the corresponding cell group to approximate it: 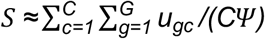. For non-tumor cells, which are commonly diploid, this assumption is assured.

In the analysis of bulk RNAseq data from mixed tumor samples, we are interested in comparing tumor with non-tumor cell groups. We let *T* denote tumor cells and *N* denote non-tumor cells. Therefore, we define a tumor-specific total mRNA expression score (TmS) to reflect the ratio of total mRNA transcript level per haploid genome of tumor cells to that of the surrounding non-tumor cells, *i.e*., *TmS_tumor_* = *S_T_* / *S_N_*, simplified as TmS from here forward. It is necessary to calculate this ratio in order to cancel out technical effects presented in sequencing data that confounds with both *S_T_* and *S_N_*. Let 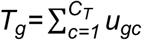 and 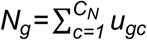 denote the total number of mRNA transcripts of gene *g* across all cells from tumor and nontumor cells, let *C_T_* and *C_N_* denote the total number of tumor and non-tumor cells, and let *ψ_T_* and *ψ_N_* represent the average ploidy of tumor and non-tumor cells, respectively. Under the assumption that the tumor cells have a similar ploidy, we can derive TmS without using single-cell-specific parameters as

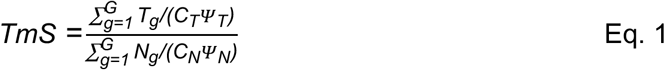

We further introduce the proportion of total bulk mRNA expression derived from tumor cells (hereafter ‘tumor-specific mRNA proportion’) 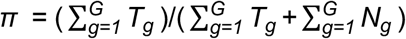 and the tumor cell proportion (hereafter ‘tumor purity’) *ρ* = *C_T_* /(*C_T_* + *C_N_*). We thus have

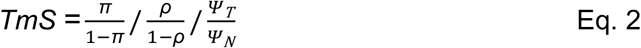

The tumor-specific mRNA proportion *π* derived from the tumor can be estimated using DeMixT^23^ as 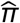; the tumor purity *ρ* and ploidy *Ψ_T_* can be estimated using ASCAT^24^, ABSOLUTE^25^ or Sequenza^40^ based on the matched DNA sequencing data as 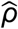 and 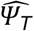, respectively; the ploidy of non-tumor cells *Ψ_N_* was assumed to be 2^24,25^. Hence, we have

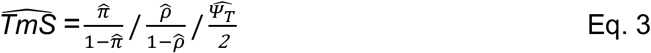

In what follows, we use *TmS* to represent 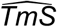 for simplicity. A full description is in **Supplementary Note, Section 2.1**.

#### Consensus of tumor purity and ploidy estimation

For DNA-based deconvolution methods such as ASCAT and ABSOLUTE, there could be multiple tumor purity *ρ* and ploidy *Ψ_T_* pairs that have similar likelihoods. Both ASCAT and ABSOLUTE can accurately estimate the product of purity and ploidy *ρΨ_T_*; however, they sometimes lack power to identify *ρ* and *Ψ_T_* separately. TmS is derived from estimates of tumor purity and ploidy via their product, automatically dealing with any ambiguity in tumor purity and ploidy estimation, ensuring the robustness of the TmS calculation. We illustrate this point by showing that among 20% of all TCGA samples, the agreement between TmS values calculated from ASCAT and ABSOLUTE are substantially improved, as compared to those for the ploidy or purity individually (**Supplementary Fig. 3f-g**). To calculate one final set of TmS values for a maximum number of samples, we take a consensus strategy. We first calculate TmS values with tumor purity and ploidy estimates derived from both ABSOLUTE and ASCAT, and then fit a linear regression model on the log_2_-transformed TmS values calculated with ASCAT by using the log_2_-transformed TmS values calculated with ABSOLUTE as a predictor variable. We remove samples with Cook’s distance ≥ 4/*n* (*n*=5,295, **Supplementary Fig. 3h**) and calculate the final 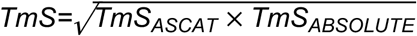.

### Improved estimation of tumor-specific mRNA proportion

The identifiability of model parameters is a major issue for high dimensional models. With the DeMixT model, there is hierarchy in model identifiability in which the cell-type specific mRNA proportions are the most identifiable parameters, requiring only a subset of genes with identifiable expression distributions. Therefore, our goal is to select an appropriate set of genes as input to DeMixT that optimizes the estimation of the tumor-specific mRNA proportions (*π*). In general, genes expressed at different numerical ranges can affect estimation of *π*. We found that including genes that are not differentially expressed between the tumor and non-tumor components, differentially expressed across tumor subtypes in different samples, or genes with large variance in expression within the non-tumor component, can introduce large biases into the estimated *π*. On the other hand, the tumor component is hidden in the mixed tumor samples, hence preventing a DE analysis between mixed and normal samples from finding the best genes. By applying a profile-likelihood based approach to detect the identifiability of model parameters^67^, we systematically evaluated the identifiability for all available genes based on the data, and selected the most identifiable genes for the estimation of *π*. As a general method, the profile-likelihood based gene selection strategy can be extended to any method that uses maximum likelihood estimation. We also employed a virtual spike-in strategy to balance proportion distributions which further improved the deconvolution performance. A full description is provided in **Supplementary Note, Section 2.2**.

#### Profile-likelihood based gene selection

Briefly, in the DeMixT model, for sample *i* ∈ (1,2,…, *M*) and gene *g* ∈ (1,2,…, *G*), we have

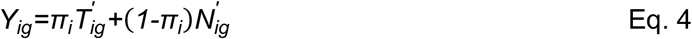

where *Y_ig_* represents the scale normalized expression matrix observed from mixed tumor samples, *T’_ig_* and *N’_ig_* represent the normalized relative expression of gene *g* within tumor and surrounding non-tumor cells, respectively. The estimated tumor-specific mRNA proportion 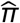 is the desirable quantity for Eq.3. We assume each hidden component follows the log_2_-normal distribution, *i.e*., 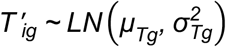 and 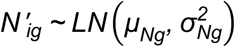. We will use notation *T* and *N* and drop the ’ sign from now on. The identifiability of a gene *k* in the DeMixT model is measured by the confidence interval 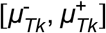 around the mean expression *μ_Tk_*. The definition of the profile likelihood function of *μ_Tk_* is

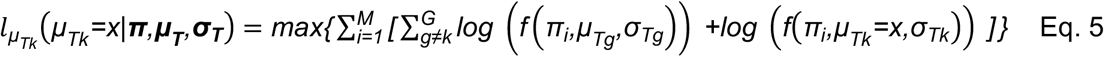

where 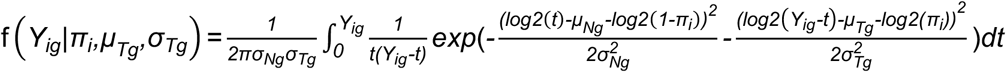 is the likelihood function of the DeMixT model.

The confidence interval of a profile likelihood function can be constructed through inverting a likelihoodratio test^68^. However, calculating the actual profile likelihood function of all genes (~20,000) is generally infeasible due to computational limits. We adopted an asymptotic approximation to quickly evaluate the profile likelihood function^67^, using the observed Fisher information of the log-likelihood, denoted as 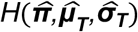. Then the asymptotic *α* level confidence interval of *μ_Tk_* can be written as^67^

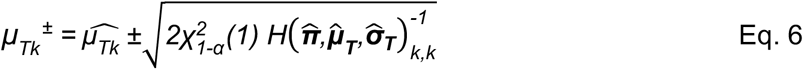

We hereby introduce a gene selection score to represent the length of an asymptotic profile-likelihood based 95% confidence interval of *μ_Tk_* for gene *k*,

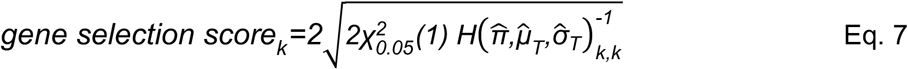

Genes with a lower score have a smaller confidence interval, hence higher identifiability for their corresponding parameters in the DeMixT. Genes are ranked based on the gene selection scores from the smallest to the largest. A subset of genes that are ranked on top will be used for parameter estimation. In the DeMixT R package (available from Bioconductor), our proposed profile-likelihood based gene selection approach is included as function “DeMixT_GS”. A full description is provided in **Supplementary Note, Section 2.2.2.**We performed a simulation study, mimicking the TCGA prostate adenocarcinoma dataset, to validate the proposed gene selection method. A full description is provided in **Supplementary Note, Section 2.2.3**. The implementation of virtual spike-ins and a simulation study is provided in **Supplementary Note, Section 2.2.4.**

### TmS validation using bulk RNAseq data from mixed cell lines

We validated TmS estimates using an experimental dataset from a previous mixed cell-line study (GEO: GSE121127)^23^, and selected a subset of 18 mixed samples with negligible immune component. Lung adenocarcinoma in humans (H1092) and cancer-associated fibroblast (CAF) cells were mixed at different cell count proportions (**Supplementary Table 3**) to generate each bulk sample, plus three additional samples of 100% H1092 or 100% CAF. The raw reads were generated from paired-end total RNA Illumina sequencing and mapped to the human reference genome build 37.2 from NCBI through TopHat^69^. SAMtools^70^ was applied to remove improperly mapped and duplicated reads. Picard tools were used to sort the cleaned SAM files according to their reference sequence names and create an index for the reads. The gene-level expression was quantified using the R packages GenomicFeatures and GenomicRanges.

For each cell line, we measured total RNA amount (in ng/ul) for 1 million cells in three repeats using the Qubit RNA Broad Range Assay Kit (Life Technologies). The true TmS values of H1092 or CAF were then derived as a ratio of the total RNA amount per cell between the two cell types. Specifically, 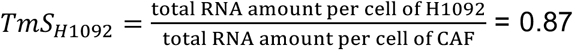, and 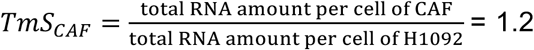. We estimated the RNA proportion of H1092 and CAFs using DeMixT (DeMixT_GS function with 4,000 genes selected), under two scenarios: (1) 3 pure CAFs samples were used as reference; (2) 3 pure H1092 samples were used as reference. To estimate TmS values, we used the known cell counts to calculate *ρ*’s.

### TmS estimation in TCGA

#### Dataset

Raw read counts of high throughput mRNA sequencing data, clinical data, and somatic mutations from 7,054 tumor samples across 15 TCGA cancer types (breast carcinoma, bladder urothelial carcinoma, colorectal cancer (colon adenocarcinoma + rectum adenocarcinoma), head-and-neck squamous cell carcinoma, kidney chromophobe, kidney renal clear cell carcinoma, kidney renal papillary cell carcinoma, liver hepatocellular carcinoma, lung adenocarcinoma, lung squamous cell carcinoma, pancreatic adenocarcinoma, prostate adenocarcinoma, stomach adenocarcinoma, thyroid carcinoma, uterine corpus endometrial carcinoma) were downloaded from the Genomic Data Commons Data Portal (https://portal.gdc.cancer.gov/). ATAC-seq data^43^, tumor purity and ploidy data^71,72^, and annotations of driver mutation and indels^51^ were downloaded for these samples. A full description of all datasets is provided in **Supplementary Note, Section 2.3.1**.

#### Estimation of tumor-specific mRNA proportions from RNAseq data

For each cancer type, we filtered out poor quality tumor and normal samples that were likely misclassified. We then selected available adjacent normal samples as reference for the tumor deconvolution using DeMixT. Based on simulation studies (**Supplementary Note 2.2.3**) and observed distributions of gene selection scores in real data, we chose the top 1,500 or 2,500 genes (varies across cancer types) to estimate tumor-specific mRNA proportions (*π*). For each cancer type, the selected 1,500 or 2,500 genes are defined as intrinsic tumor signature genes. We added varying number of virtual spike-in samples depending on cancer types. We additionally removed samples with extreme estimates of *π*, > 85% or ranked at the top 2.5 percentile of all samples within each cancer type, to mitigate the remaining underestimation when *π* is close to 1. A full description is provided in **Supplementary Note, Section 2.3.2.1**.

#### Consensus TmS estimation

We calculated a consensus TmS as 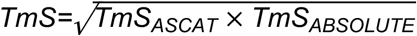 and removed 264 TCGA samples that deviated from our consensus model as described previously. A full description on sample exclusions is provided in **Supplementary Note, Section 2.3.2.2**.

#### Intrinsic tumor signature genes

For each cancer type, we conducted gene set enrichment analyses on Hallmark pathways and KEGG pathways^38^ for all genes with their gene selection scores using GSEA^73^ and g:Profiler^74^. The genes were ranked according to their gene selection scores from the smallest to the largest. We further evaluated the chromatin accessibility of intrinsic tumor signature genes using ATAC-seq data from TCGA samples^43^. For each sample, we calculated the mean of the peak scores of all intrinsic tumor signature genes and compared it with the corresponding permuted null distribution for each cancer type. A full description is provided in **Supplementary Note, Section 2.3.2.3**.

### TmS estimation in ICGC-EOPC (Early-Onset Prostate Cancer)

In this cohort, matched mRNA sequencing data and whole genome sequencing data, as well as clinical data including biochemical recurrence, Gleason score and pathologic stage, from 121 tumor samples and 9 adjacent normal samples from 96 patients (age at treatment < 55), were downloaded from Gerhauser *et al*.^29^ We used the 9 available adjacent normal samples as the normal reference to run DeMixT. The mRNAseq data came from 3 batches - batch 1 (17 patients, 25 samples), batch 2 (42 patients, 52 samples), and batch 3 (37 patients, 44 samples). To evaluate and adjust for potential batch effects, we applied the DeMixT deconvolution pipeline in three scenarios: (1) ran all samples together; (2) ran each batch separately; (3) applied Combat^65^ to adjust for batch effect then ran all samples together. The Spearman correlation coefficients between tumor-specific mRNA proportions obtained in pairwise comparisons were high: 0.9, 0.8, and 0.8. Given the observed consistency and the robustness of DeMixT to select intrinsic tumor signature genes, we report final results from scenario (1). We removed samples with *π* > 85%, without DNA-based tumor purity and ploidy or missing follow-up information of biochemical recurrence intervals (see **Supplementary Note 3** for further details).

### TmS estimation for patients with Renal Medullary Carcinoma

This dataset contains mRNAseq data of 11 tumor samples, each from a patient affected with renal medullary carcinoma (RMC)^30^. We used 7 available adjacent normal samples from this cohort as the normal reference to run DeMixT. The final tumor-specific mRNA proportions were estimated using the DeMixT_GS mode with the top 1,500 genes. For each patient sample, tumor purity and ploidy were estimated based on the matched whole exome sequencing (WES) data by Sequenza. We removed 2 patients from TmS calculation, as one is less RMC-like^30^, and the other one has unstable estimates of tumor purity.

### TmS estimation for patients with high-risk prostate cancer and treated with ADT+ARSI

This dataset contains 27 RNAseq data of tumor samples. We removed 1 sample for its low Gleason score of 6. We ran DeMixT on RNAseq raw count data from these 26 samples together with TCGA-PRAD tumor samples with a Gleason score of 8+. This dataset does not contain RNAseq data from adjacent normal samples, so we used RNAseq data from 47 normal prostate samples in TCGA as reference. To mitigate the batch effects between the 26 samples and the TCGA PRAD samples, scale normalization together with ComBat^65^ were applied before deconvolution. The final tumor-specific mRNA proportions were estimated using the DeMixT_GS mode with the top 1,500 genes. For each sample, tumor purity and ploidy were estimated using the WES data by Sequenza. We removed 2 tumor samples which presented an unstable estimation of tumor purity, *i.e*., SD = 0.46 and 0.24, across three top purity estimates provided by Sequenza.

### Statistical analysis

#### Association with clinical variables

Kruskal-Wallis tests were used to compare the distribution of TmS between subgroups defined by each clinical variable (**Fig. 4a-e**). The *P* values from the Kruskal-Wallis tests were adjusted using BH correction across all available clinical variables within the corresponding cancer type.

#### Association with survival outcomes

We evaluated the association between TmS and survival outcome (overall survival and progression-free interval) across 15 cancer types in TCGA and ICGC-EOPC. To ensure sufficient sample size in each group, we summarized pathologic stages into two categories: early (I/II) and advanced (III/IV). For prostate cancers, we used Gleason score (Gleason Score = 7 versus 8+) instead of early and advanced stages. We used a recursive partitioning survival tree model, rpart^75^, to find the optimal TmS cutoff separating different survival outcomes within each of the two stages defined above in each cancer type. Splits were assessed using the Gini index, and the maximum tree depth was set to 2. Log-rank tests between high and low TmS groups within early or advanced pathological stages were performed. We then fitted multivariate Cox Proportional Hazard models with age, TmS, stage, and an interaction term of TmS and Stage (TmS x stage) as predictors of overall survival and progression free interval for each TCGA cancer type. We also applied the same procedure to two other metrics for comparison: ploidy and TmS x ploidy. We provide further information in **Supplementary Note, Section 3**.

#### Association with metabolism, hypoxia, and genomic dysregulations

For each cancer type from TCGA, we conducted gene set enrichment analyses^73^ on the metabolism of carbohydrates pathways (the Reactome database^48^), including the pentose phosphate pathway, glucose metabolism, galactose catabolism, fructose metabolism, formation of xylulose-5-phosphate, glycosaminoglycan metabolism, glycogen metabolism, and blood group systems biosynthesis. The genes were ranked by the Spearman correlation coefficient between their expression levels and TmS across samples; they were then put through GSEA in the “pre-ranked” mode. For GSEA, we adopted permutation tests (1,000 times) to generate a normalized enrichment score (NES) for each candidate pathway. A hierarchical clustering on the expression levels of the Reactome pentose phosphate pathway (15 genes total, of which 2 genes were removed due to high-frequency zero counts across samples) for the tumor samples was performed using Euclidean distance and Ward linkage. The samples were then separated into two groups using the “cutree” function. For each cancer type, a Wilcoxon rank-sum test was used to compare the distributions of TmS estimates between the two tumor sample groups. *P* values were adjusted for multiple testing using the BH correction across all cancer types. For each tumor sample, we further defined a pentose-phosphate-13 score, *i.e*., a score based on the 13-gene-set, as the enrichment score calculated by gene set variation analysis (GSVA)^76^.

Hypoxia scores were generated as described previously^49,77^ using the Buffa Signature^78^, which has been previously shown to be well-correlated with direct and transcriptional measures of hypoxia^79^. High values of this score suggest more hypoxia (lower levels of oxygen), while low values suggest less hypoxia (higher levels of oxygen). See further information in **Supplementary Note, Section 4**.

We conducted a network analysis of the pentose phosphate pathway (15 genes) and hypoxia signature gene list (51 genes) using a genome-wide parsimonious composite network (PCNet)^80^ with visualization and analysis performed with Cytoscape version 3.8.2^81^.

Tumor mutation burden (TMB) was calculated by counting the total number of somatic mutations based on the consensus mutations calls (MC3)^82^. Chromosomal Instability (CIN) scores were calculated as the ploidy-adjusted percent of genome with an aberrant copy number state. ASCAT was used to calculate allele-specific copy numbers^24^. For samples present in both TCGA and PCAWG, the consensus copy number was derived from published results^83^. Tumor samples that had undergone WGD were identified based on homologous copy-number information^25^.

For each cancer type within TCGA, we considered genes which had driver mutations (including nonsense, missense and splice-site SNVs and indels)^51^ in at least 10 samples. For each of these genes, samples were labelled as “with driver mutation” if they carried at least one driver mutation in that gene or “without driver mutation” otherwise. We investigated 24 cancer-gene pairs for driver mutation analysis. We applied a Wilcoxon rank-sum test to each candidate gene to compare the distributions of TmS of the samples with driver mutations versus without driver mutations. The *P* values of each gene were adjusted for multiple testing using BH correction across all candidate genes within the corresponding cancer type.

We also implemented an agnostic search among non-synonymous mutations (including SNVs and indels) over all genes for the 15 cancer types to identify those that were significantly associated with TmS. For each cancer type, we considered a gene as a candidate gene if there were at least 10 samples containing non-synonymous mutations in that gene. We investigated 32,894 cancer-gene pairs for non-synonymous mutation analysis. We applied two statistical tests to evaluate the difference between the “with non-synonymous mutation” and “without non-synonymous mutation” samples. We first applied a Wilcoxon rank-sum test for each candidate gene to evaluate the difference between the distribution of TmS of the two group of samples. We then fitted a linear regression model using log_2_-transformed TmS as the dependent variable and mutation status as a predictor: *log_2_(TmS)=b_0_+b_1_ log_2_*(*TMB*) +*b_2_MUT*, where TMB represents tumor mutation burden. *MUT*=1 if the sample has at least one non-synonymous mutation in the candidate gene, and *MUT*=0 otherwise. The *P* values were calculated by a t-test of the regression coefficient *b_2_*. The *P* values of each gene based on Wilcoxon rank-sum test and t-test were adjusted by BH correction based on the number of candidate genes within the corresponding cancer type.

## Data availability

The UMI counts of the hepatocellular carcinoma single cell RNA sequencing data were downloaded from the Gene Expression Omnibus (GEO) with the accession code GSE125449. The UMI counts and cell type annotations of the lung adenocarcinoma single cell RNA sequencing data were downloaded from the ArrayExpress under accessions E-MTAB-6149. The UMI counts of the colorectal adenocarcinoma single cell RNA sequencing data are available at http://crcmoonshot.org/?page_id=189. FASTQ files of single-cell RNA sequencing data from pancreatic cancer will be publicly available on the GEO with the accession code GSE156405. UMI counts of the single cell RNA sequencing data from LNCaP cell lines will be public available on the GEO with the accession code GSE168668.

Raw read counts of RNA sequencing data, clinical data, and somatic mutations from 7,054 tumor samples across 15 TCGA cancer types are available for download from the Genomic Data Commons Data Portal (https://portal.gdc.cancer.gov/). ATACseq data for TCGA samples were downloaded from https://science.sciencemag.org/content/362/6413/eaav1898/tab-figures-data.

Clinical information of ICGC-EOPC was downloaded from https://www.sciencedirect.com/science/article/pii/S1535610818304823?via%3Dihub#gs1.

RNAseq data from patients with Renal Medullary Carcinoma is available at the NCBI Sequence Read Archive (SRA) hosted by the NIH (SRA accession: PRJNA605003): https://www.ncbi.nlm.nih.gov/sra/?term=PRJNA605003.

All other relevant data are available from the corresponding author upon reasonable request.

## Code availability

All code used for analyses was written in R version 3.6.1 and will be made available. The core computational pipelines developed for estimating tumor-specific mRNA expression proportion are available in R package DeMixT1.8.0, which can be downloaded from https://www.bioconductor.org/packages/release/bioc/html/DeMixT.html.

**Supplementary Figure. 1: High diversity of total mRNA expression in tumor cells. a,** Flowchart of scRNAseq data preprocessing. **b,** Illustration of expressed genes in tumor cells (left panels) compared to non-tumor cells: epithelial and stromal cells (middle panels) and immune cells (right panels). The data shown are based on cells randomly selected from each of the four “patient 1” samples with colorectal, hepatocellular, lung and pancreatic cancers. In each heatmap, expressed genes (UMI count > 0) are shown in black, and non-expressed genes (UMI count = 0) are shown in gray. Cells in the rows and genes in the columns are ordered from high to low by the total numbers of expressed genes and the number of cells with detected expression of each gene, respectively. Barplots provide the corresponding distributions of gene counts and total UMI counts.

**Supplementary Figure. 2: Using gene counts and total UMI counts to measure the global gene expression heterogeneity. a,** Distributions of gene counts and total UMI counts by cell type in scRNAseq data from seven patients with hepatocellular, lung or pancreatic cancers. The top x-axis annotates total UMI counts (means and 95% Confidence Intervals, CIs). The bottom x-axis annotates gene count distribution (density). Density curves are colored for tumor cells and shown in grayscale for non-tumor cells. Clusters with higher gene counts are shown in darker shades. Numbers in the parentheses indicate the number of cells analyzed. **b,** Monocle-inferred trajectories for tumor cells from four patients with lung and pancreatic cancers. Cells on the trees are colored by total UMI counts. Average differentiation scores by CytoTRACE for high- and low-UMI count tumor cell clusters are labelled. **c,** Distribution of cell cycle scores in tumor cell clusters from seven scRNAseq patient samples. *P* values of Wilcoxon rank-sum tests comparing the cell cycle scores across clusters are indicated by asterisks (* *P* < 0.05, ** < 0.01, *** < 0.001).

**Supplementary Figure. 3: Consensus estimation of TmS from matched RNAseq and DNAseq data in TCGA. a**, Quantitative relationship between cells, chromosomal copies and mRNA contents. Three example types of cells with ploidy = 2, 3, or 4 are given. Under the scenario of linear dosage effects, as shown in the boxes with a yellow background, suppose their corresponding total mRNA amounts are 2, 3, and 4, then the ploidy-adjusted, or per haploid genome, total mRNA amount would be 1, 1, and 1. Under the scenario of dosage compensation, *i.e*., more chromosomal copies but maintaining the same total dose, the second cell has a total mRNA amount of 2 and a per haploid genome value of 0.67. Under the scenario of dosage transgression, *i.e*., more chromosomal copies with more dose per copy, the third cell has a total mRNA amount of 6 and a per haploid genome value of 1.5. **b**, Definition of TmS and its analysis pipeline. **c**, Distribution of tumor-specific mRNA proportions estimated by DeMixT across cancer types. **d-e,** Distributions of tumor cell proportions estimated by (**d**) ASCAT or (**e**) ABSOLUTE across cancer types. **f**, Smoothed scatter plot of tumor ploidy estimates from ABSOLUTE *vs*. ASCAT across all samples. Gray points correspond to 968 samples that presented inconsistent tumor ploidy (and purity) estimates between the two methods. **g**, TmS estimates using either ABSOLUTE or ASCAT-derived purity and ploidy estimates with or without ploidy adjustment for the 968 discordant samples from (**f**). Blue and gray points correspond to TmS prior to and after ploidy adjustment, respectively. Ploidy adjustment improved consistency between the ABSOLUTE and ASCAT results. **h**, Scatter plot of TmS calculated using the two methods. A linear regression model was fitted using log_2_(TmS estimated by ABSOLUTE) as the predicted variable and log_2_(TmS estimated by ASCAT) as the predictor variable. Red points are outliers with a Cook’s distance ≥ 4/*n*, where *n* = 5,295 for the total number of TCGA samples. Cyan points are the remaining samples (95%) that showed a good fit for the model and hence their TmS estimates are deemed consistent and robust across two DNAseq deconvolution methods. **i**, Total mRNA proportion estimation for H1092 and CAF using DeMixT in the benchmarking study. The concordance correlation coefficient (CCC) for two variables x (true tumor-specific RNA proportions) and y (estimated tumorspecific RNA proportions) is expressed as 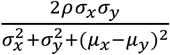, where *μ* and *σ*^2^ represent the mean and variance, and *ρ* is the Pearson correlation coefficient.

**Supplementary Figure. 4: Intrinsic tumor signature genes across cancer types. a**, Illustration of the RNAseq deconvolution workflow with intrinsic tumor signature genes selected using a profile-likelihood based gene selection score. **b**, Distributions of gene selection scores across four types of genes in a simulation study (**Supplementary Note 2.2**). For the profile-likelihood based gene selection, genes are ranked from the smallest to the largest score (left). For the DE based gene selection, genes are ranked from the largest to the smallest absolute t-statistics (middle). *P* values of Wilcoxon rank-sum tests are indicated by asterisks (* *P* < 0.05, ** < 0.01, *** < 0.001). The types of genes among the top 1,500 selected gene are shown (right) for the two rankings. Ideally only genes consistently differentially expressed between *T* and *N* should be selected. This is achieved by the profile-likelihood method but not the DE method. **c**, Histogram of the number of overlapping genes across cancer types and their annotation categories. The y axis represents the total number of genes and the x axis represents the number of cancer types for which a gene was selected. **d**, Heatmap of normalized enrichment scores of top cancer hallmark pathways and KEGG pathways. Only pathways with adjusted *P* value < 0.05 are colored. **e**, M-A plot comparing ATAC-seq peak scores of intrinsic tumor signature genes (signature) *vs*. other genes (non-signature) from matched tumor samples in each cancer type. Samples above the dashed line have higher ATAC-seq peaks score in intrinsic tumor signature genes compared to those in non-signature genes. Samples with adjusted *P* values < 0.05 from permutation tests are shown as circles.

**Supplementary Figure. 5: TmS refines prognostication on pathological stages. a**, Distribution of TmS for early (stage I and II) *vs*. advanced (stage III and IV) pathological stages across 15 cancer types. Adjusted *P* values of Wilcoxon rank-sum tests are indicated by asterisks (* *P* < 0.05, ** < 0.01, *** < 0.001). **b**, Spearman correlation coefficients between TmS and the expression levels of *MKI67* and *PCNA* across 15 cancer types. **c**, Distribution of TmS for female and male patient samples in TCGA across 15 cancer types. None of the adjusted *P* values of Wilcoxon rank-sum tests comparing TmS between the two groups reached significance at a confidence level of 0.05. Brown circles (read out on the right y-axis) represent Spearman correlation coefficients between TmS and age within the same sex and cancer type. The red dotted horizontal line represents TmS equal to 1 (left y axis) and correlation equal to 0 (right y axis). None of the adjusted *P* values for correlation tests reached significance at a confidence level of 0.05. **d**, KM curves of PFI for TCGA samples. Gray lines denote summary KM curves of patients with high *vs*. low TmS across all cancer types. KM curves are further grouped by TmS and pathological stage into four groups. *P* values of log-rank tests between high *vs*. low TmS groups are indicated by asterisks (* *P* < 0.05, ** < 0.01, *** < 0.001). **e**, KM survival curves for individual cancer types with pattern I. **f**, KM survival curves for individual cancer types with pattern II. **g**, KM survival curves for renal papillary carcinoma with pattern III.

**Supplementary Figure. 6: Prognostication using ploidy or ploidy-unadjusted TmS on pathological stages. a**, Scatter plots of TmS (y axis) *vs*. tumor ploidy (x axis) for samples from TCGA patient cohorts with head-and-neck squamous cell carcinoma (HPV negative), lung squamous cell carcinoma, renal clear cell carcinoma, and colorectal carcinoma. The samples were grouped into high *vs*. low TmS within early or advanced pathological stages, with different groups shown in distinct colors. **b**, KM survival curves of overall survival in four cancer types over patient groups defined by ploidy and stage. We grouped patients into high *vs*. low ploidy based on a cutoff of 2.5 within early or advanced pathological stage. **c**, KM survival curves of overall survival in four cancer types over patient groups defined by ploidy-unadjusted TmS and stage. **d**, KM survival curves of overall survival in four cancer types over patient groups defined by TmS and stage. *P* values of log-rank tests pairs of patient groups are shown with matching colors and are indicated by asterisk (* *P* < 0.05, ** < 0.01, *** < 0.001).

**Supplementary Figure. 7: Pan-cancer contributions of metabolic and hypoxia levels to TmS. a**, Heatmap of normalized enrichment scores (NES) of Reactome metabolism of carbohydrates pathways across 15 cancer types in TCGA. Pathways are ordered by the mean NES across 15 cancer types, from high to low. **b**, Hierarchical clustering on the expression levels of Reactome pentose phosphate pathway genes for the tumor samples from renal clear cell carcinoma. Samples were separated into two groups. **c**, Smoothed scatter plots showing correlations between pentose-phosphate-13 score and hypoxia score in each cancer type. Spearman correlation coefficients are labelled on top. **d**, Interaction network of genes from the pentose phosphate pathway and hypoxia signature. Each node represents a gene and each edge represents a known gene-gene interaction in the PCNet database.

**Supplementary Figure. 8: TmS is associated with tumor genomic features across cancer types. a-c**, Distribution of TmS for patient samples with (**a**) high or low tumor mutation burden (TMB); (**b**) high or low chromosomal instability score; (**c**) with or without a whole genome duplication event. Patient groups are divided in half at the median TMB and chromosomal instability scores in (**a**) and (**b**) respectively. Adjusted *P* values of Wilcoxon rank-sum tests are indicated by asterisks (* *P* < 0.05, ** < 0.01, *** < 0.001).

## Acknowledgments

S.C. is supported by the Norman Jaffe Professorship in Pediatrics Endowment Fund, MD Anderson Colorectal Cancer Moon Shot Program, and NIH R01CA183793. J.R.W. is supported by American Thyroid Association/ThyCa grant, Mark Foundation for Cancer Research ASPIRE award. S.J. is supported by Human Cell Atlas Seed Network - Breast, Chan Zuckerberg Institute, MD Anderson Colorectal Cancer Moon Shot Program, NIH R01CA183793. P.Y. is supported by NIH R01CA239342. J.C. is supported by NIH R01CA158113. P.V.L. and J.D. are supported in part by the Francis Crick Institute, which receives its core funding from Cancer Research UK (FC001202), the UK Medical Research Council (FC001202), and the Wellcome Trust (FC001202). For the purpose of Open Access, the authors have applied a CC BY public copyright lisence to any Author Accepted Manuscript version arising from this submission. P.V.L. and J.D. are also supported in part by the Medical Research Council (grant number MR/L016311/1). J.D. is supported in part by the European Union’s Horizon 2020 research and innovation program (Marie Skłodowska-Curie Grant Agreement No. 703594-DECODE) and the Research Foundation – Flanders (FWO, Grant No. 12J6916N). J.P.S. is supported by the Cancer Prevention and Research Institute of Texas as a CPRIT Scholar in Cancer Research and by National Institutes of Health (K22CA234406). J.J.L. is supported by the NIH (T32CA009599). S.T. and M.N. are supported by The Academy of Finland (2011–present); Cancer Society of Finland (2013–present); Sigrid Juselius Foundation (2016–present); Finnish Cultural Foundation (2020-present). N.C.D. is supported by the Norman Jaffe Professorship in Pediatrics Endowment Fund. P.A.F. is supported in part by the Welch Foundation, MEI Pharma, Inc., Cancer Research United Kingdom (CRUK), Kadoorie Charitable Foundation, NIH/NCI (U01 CA224044-01A1, 1R01CA231465-01A1). B.L. is supported by the Single cell transcriptome of IBC cells and surrounding microenvironment, SWOG HOPE foundation, Human Cell Atlas Seed Network - Breast, Chan Zuckerberg Institute. P.M. is supported by a Career Development Award by the American Society of Clinical Oncology, a Research Award by KCCure, the MD Anderson Khalifa Scholar Award, and the MD Anderson Physician-Scientist Award. P.C.B. is supported by the NIH/NCI under awards number P30CA016042, 1U01CA214194-01 and 1U24CA248265-01. A.U. and N.E. are supported by the Norwegian Cancer Society (grant number 198016-2018). J.Z. is supported by the MD Anderson Physician Scientist Program, the MD Anderson Lung Cancer Moon Shot Program and the Cancer Prevention and Research Institute of Texas Multi-Investigator Research Award grant (RP160668). A.M. is supported by the MD Anderson Pancreatic Cancer Moon Shot Program, the Khalifa Bin Zayed Al-Nahyan Foundation, and the National Institutes of Health (NIH U01CA196403, U01CA200468, U24CA224020, P50CA221707). PVL is a Winton Group Leader in recognition of the Winton Charitable Foundation’s support towards the establishment of The Francis Crick Institute. W.W. is supported by Human Cell Atlas Seed Network - Retina, Chan Zuckerberg Institute, NIH R01CA183793, NIH R01CA239342, NIH R01CA158113, P30CA016672.

## Declaration of Interests

A.M. receives royalties for a pancreatic cancer biomarker test from Cosmos Wisdom Biotechnology, and this financial relationship is managed and monitored by the UTMDACC Conflict of Interest Committee. A.M. is also listed as an inventor on a patent that has been licensed by Johns Hopkins University to Thrive Earlier Detection. J.Z. reports research funding from Merck, Johnson and Johnson, and consultant fees from BMS, Johnson and Johnson, AstraZeneca, Geneplus, OrigMed, Innovent outside the submitted work. P.M. has received honoraria for service on a Scientific Advisory Board for Mirati Therapeutics and BMS, non-branded educational programs supported by Exelixis and Pfizer, and research funding for clinical trials from Takeda, BMS, Mirati Therapeutics, Gateway for Cancer Research, and UT MD Anderson Cancer Center, all outside the submitted work.

