## Supplementary Figures and Tables for "Tumor cell total mRNA expression shapes the molecular and clinical phenotype of cancer"

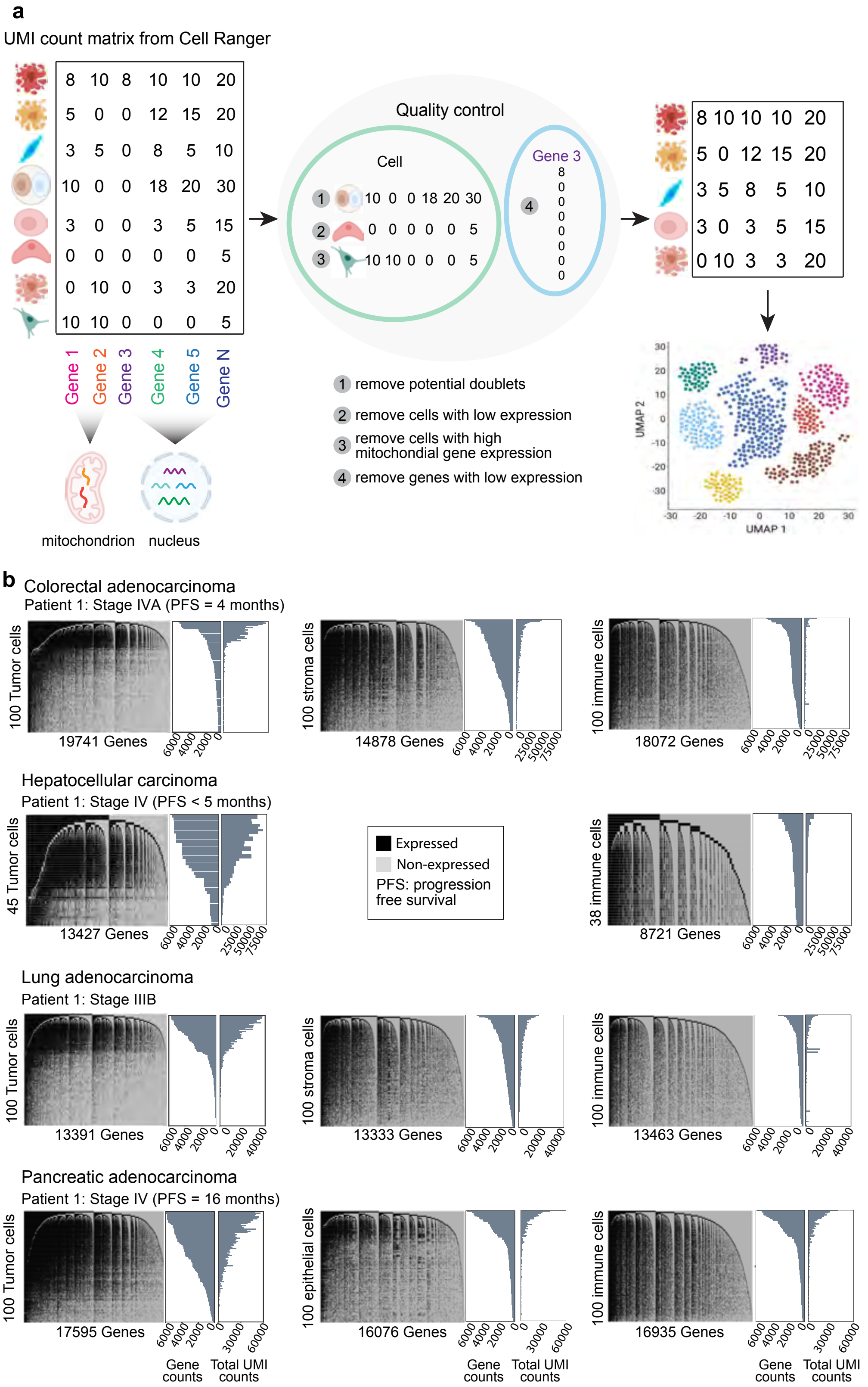


**Supplementary Figure. 1: High diversity of total mRNA expression in tumor cells. a,** Flowchart of scRNAseq data preprocessing. **b,** Illustration of expressed genes in tumor cells (left panels) compared to non-tumor cells: epithelial and stromal cells (middle panels) and immune cells (right panels). The data shown are based on cells randomly selected from each of the four "patient 1" samples with colorectal, hepatocellular, lung and pancreatic cancers. In each heatmap, expressed genes (UMI count > 0) are shown in black, and non-expressed genes (UMI count = 0) are shown in gray. Cells in the rows and genes in the columns are ordered from high to low by the total numbers of expressed genes and the number of cells with detected expression of each gene, respectively. Barplots provide the corresponding distributions of gene counts and total UMI counts.

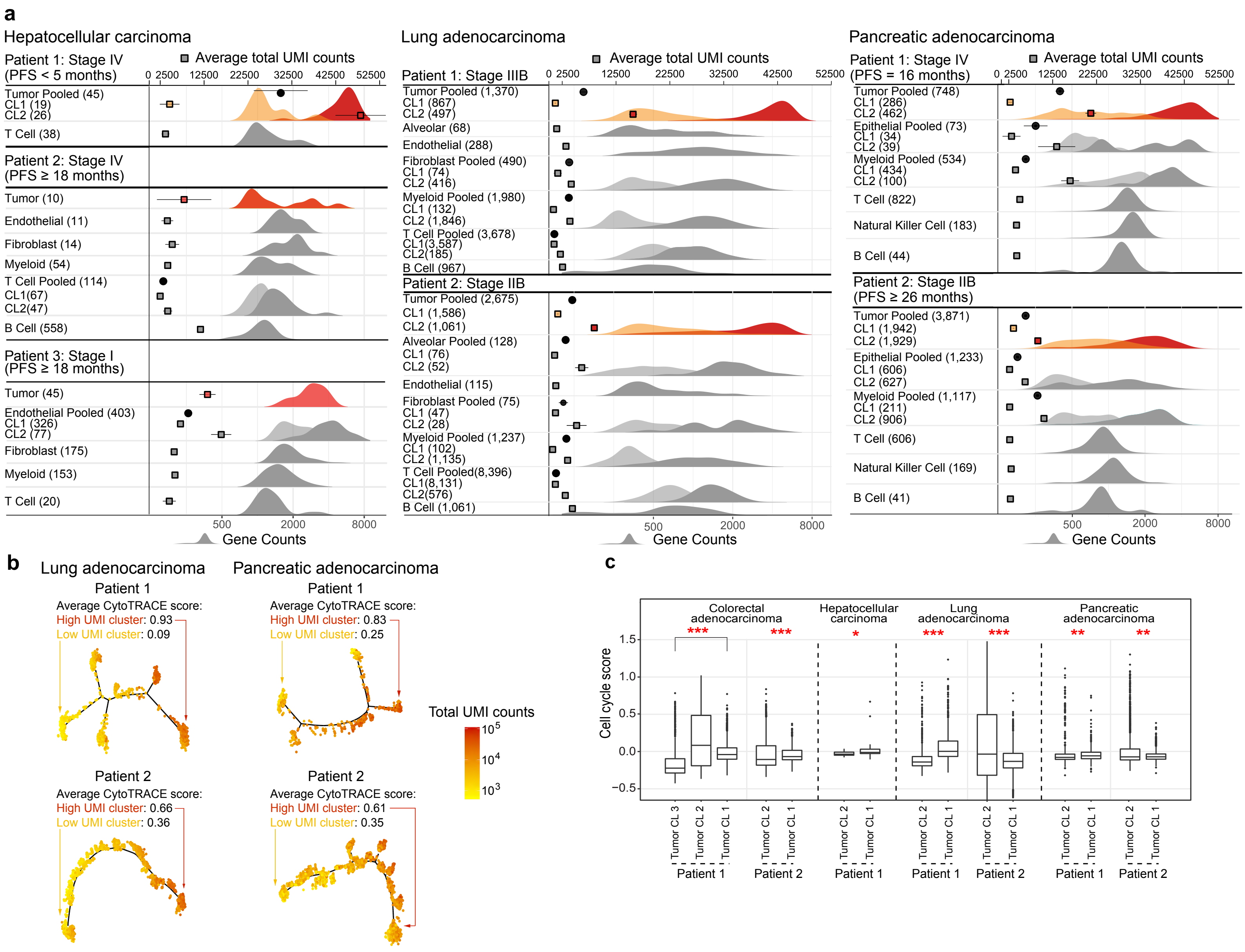


**Supplementary Figure. 2: Using gene counts and total UMI counts to measure the global gene expression heterogeneity. a,** Distributions of gene counts and total UMI counts by cell type in scRNAseq data from seven patients with hepatocellular, lung or pancreatic cancers. The top x-axis annotates total UMI counts (means and 95% Confidence Intervals, CIs). The bottom x-axis annotates gene count distribution (density). Density curves are colored for tumor cells and shown in grayscale for non-tumor cells. Clusters with higher gene counts are shown in darker shades. Numbers in the parentheses indicate the number of cells analyzed. **b,** Monocle-inferred trajectories for tumor cells from four patients with lung and pancreatic cancers. Cells on the trees are colored by total UMI counts. Average differentiation scores by CytoTRACE for high- and low-UMI count tumor cell clusters are labelled. **c,** Distribution of cell cycle scores in tumor cell clusters from seven scRNAseq patient samples. *P* values of Wilcoxon rank-sum tests comparing the cell cycle scores across clusters are indicated by asterisks (* *P* < 0.05, ** < 0.01, *** < 0.001).

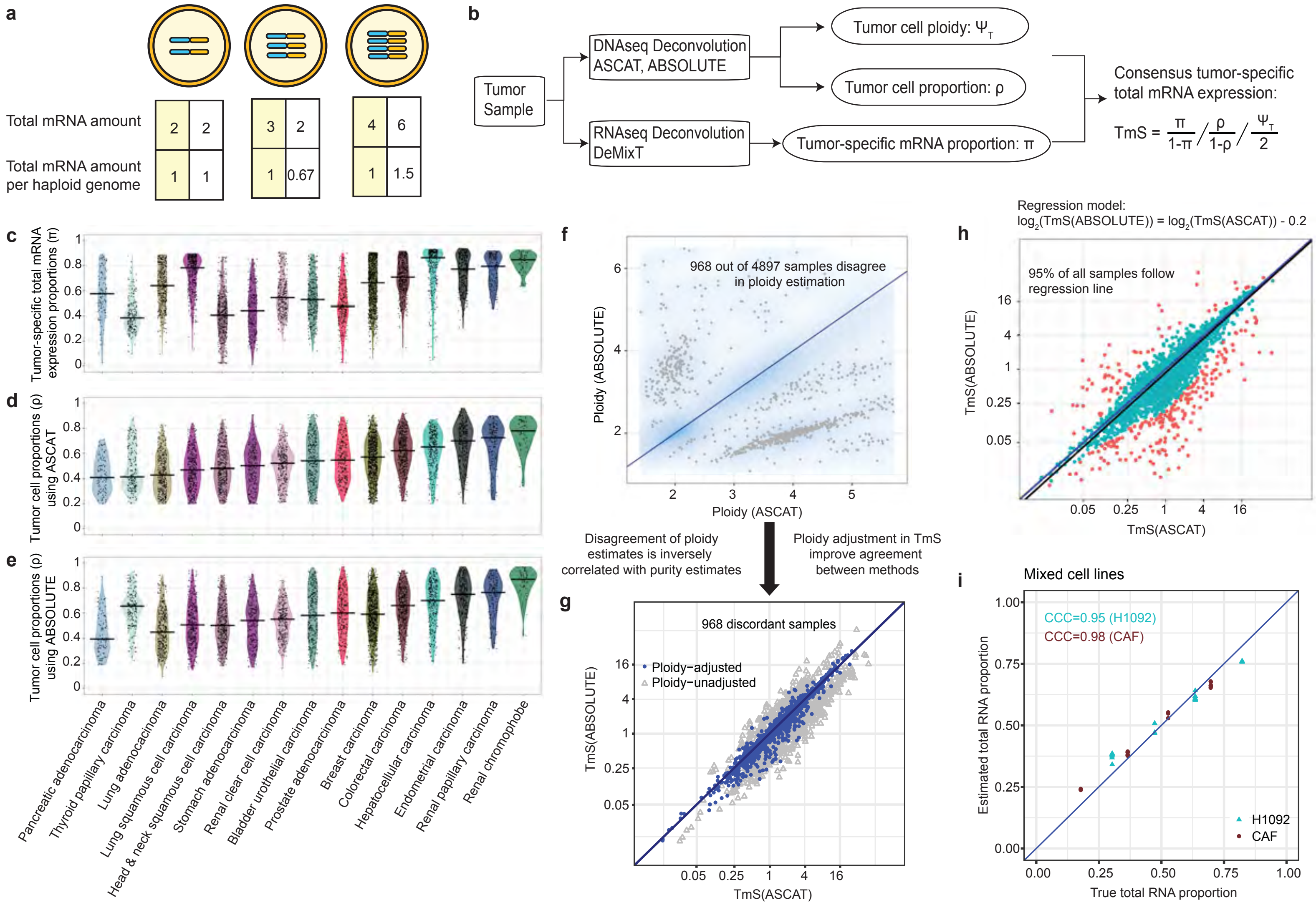


**Supplementary Figure. 3: Consensus estimation of TmS from matched RNAseq and DNAseq data in TCGA.**

**Supplementary Figure. 3 (cont'd): a**, Quantitative relationship between cells, chromosomal copies and mRNA contents. Three example types of cells with ploidy = 2, 3, or 4 are given. Under the scenario of linear dosage effects, as shown in the boxes with a yellow background, suppose their corresponding total mRNA amounts are 2, 3, and 4, then the ploidy-adjusted, or per haploid genome, total mRNA amount would be 1, 1, and 1. Under the scenario of dosage compensation, *i.e.*, more chromosomal copies but maintaining the same total dose, the second cell has a total mRNA amount of 2 and a per haploid genome value of 0.67. Under the scenario of dosage transgression, *i.e.*, more chromosomal copies with more dose per copy, the third cell has a total mRNA amount of 6 and a per haploid genome value of 1.5. **b**, Definition of TmS and its analysis pipeline. **c**, Distribution of tumor-specific mRNA proportions estimated by DeMixT across cancer types. **d-e,** Distributions of tumor cell proportions estimated by (**d**) ASCAT or (**e**) ABSOLUTE across cancer types. **f**, Smoothed scatter plot of tumor ploidy estimates from ABSOLUTE *vs*. ASCAT across all samples. Gray points correspond to 968 samples that presented inconsistent tumor ploidy (and purity) estimates between the two methods. **g**, TmS estimates using either ABSOLUTE or ASCAT-derived purity and ploidy estimates with or without ploidy adjustment for the 968 discordant samples from (**f**). Blue and gray points correspond to TmS prior to and after ploidy adjustment, respectively. Ploidy adjustment improved consistency between the ABSOLUTE and ASCAT results. **h**, Scatter plot of TmS calculated using the two methods. A linear regression model was fitted using $\log_2$(TmS estimated by ABSOLUTE) as the predicted variable and $\log_2$(TmS estimated by ASCAT) as the predictor variable. Red points are outliers with a Cook's distance ≥ 4/*n*, where *n* = 5,295 for the total number of TCGA samples. Cyan points are the remaining samples (95%) that showed a good fit for the model and hence their TmS estimates are deemed consistent and robust across two DNAseq deconvolution methods. **i**, Total mRNA proportion estimation for H1092 and CAF using DeMixT in the benchmarking study. The concordance correlation coefficient (CCC) for two variables x (true tumor-specific RNA proportions) and y (estimated tumor-specific RNA proportions) is expressed as $\frac{2\rho\sigma_x\sigma_y}{\sigma_x^2+\sigma_y^2+(\mu_x-\mu_y)^2}$, where $\mu$ and $\sigma^2$ represent the mean and variance, and $\rho$ is the Pearson correlation coefficient.

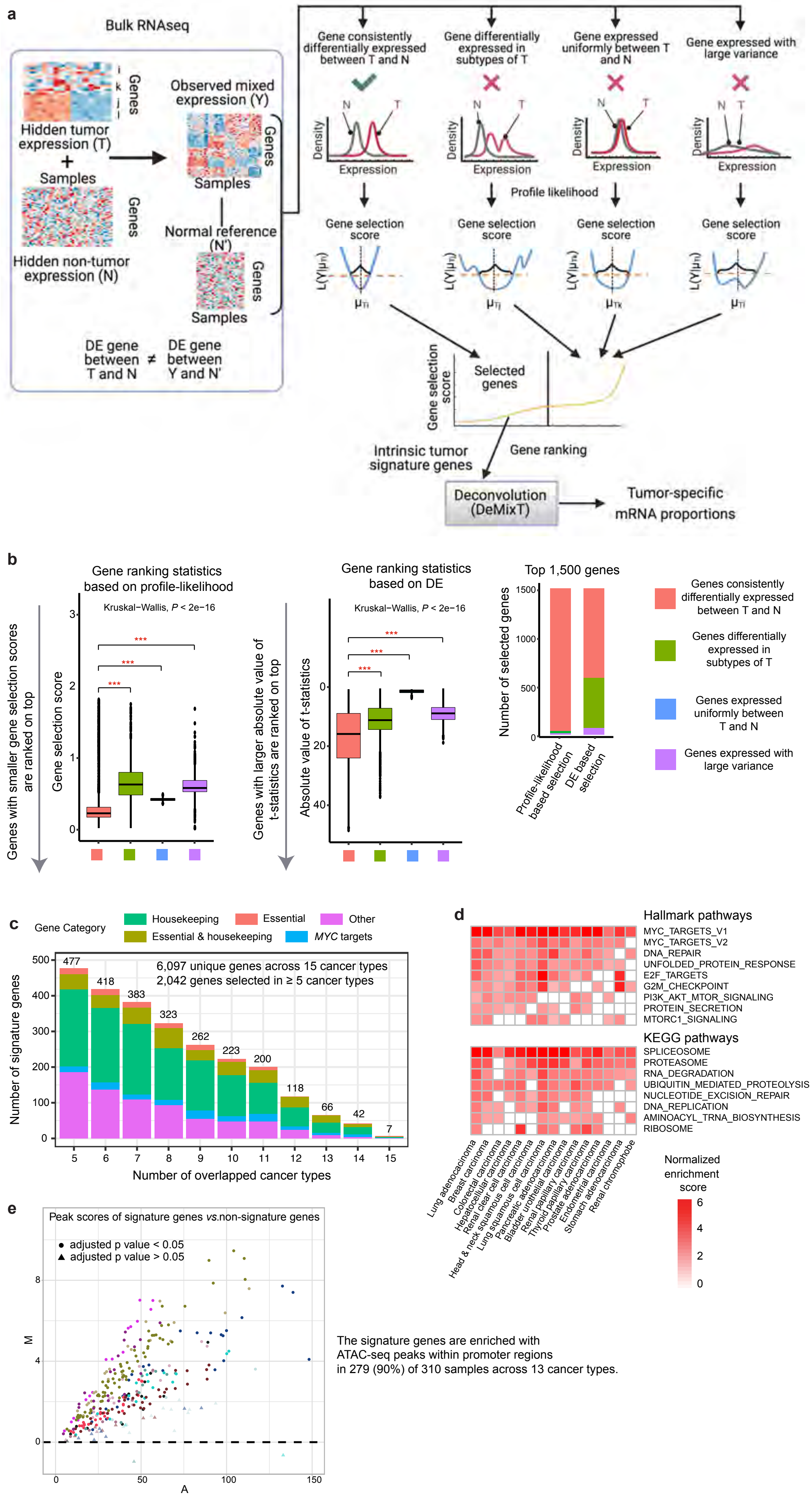


**Supplementary Figure. 4: Intrinsic tumor signature genes across cancer types.**

**Supplementary Figure. 4 (cont'd): a**, Illustration of the RNAseq deconvolution workflow with intrinsic tumor signature genes selected using a profile-likelihood based gene selection score. **b**, Distributions of gene selection scores across four types of genes in a simulation study (**Supplementary Note 2.2**). For the profile-likelihood based gene selection, genes are ranked from the smallest to the largest score (left). For the DE based gene selection, genes are ranked from the largest to the smallest absolute t-statistics (middle). *P* values of Wilcoxon rank-sum tests are indicated by asterisks (* $P < 0.05$, ** < 0.01, *** < 0.001). The types of genes among the top 1,500 selected gene are shown (right) for the two rankings. Ideally only genes consistently differentially expressed between *T* and *N* should be selected. This is achieved by the profile-likelihood method but not the DE method. **c**, Histogram of the number of overlapping genes across cancer types and their annotation categories. The y axis represents the total number of genes and the x axis represents the number of cancer types for which a gene was selected. **d**, Heatmap of normalized enrichment scores of top cancer hallmark pathways and KEGG pathways. Only pathways with adjusted *P* value < 0.05 are colored. **e**, M-A plot comparing ATAC-seq peak scores of intrinsic tumor signature genes (signature) *vs*. other genes (non-signature) from matched tumor samples in each cancer type. Samples above the dashed line have higher ATAC-seq peaks score in intrinsic tumor signature genes compared to those in non-signature genes. Samples with adjusted *P* values < 0.05 from permutation tests are shown as circles.

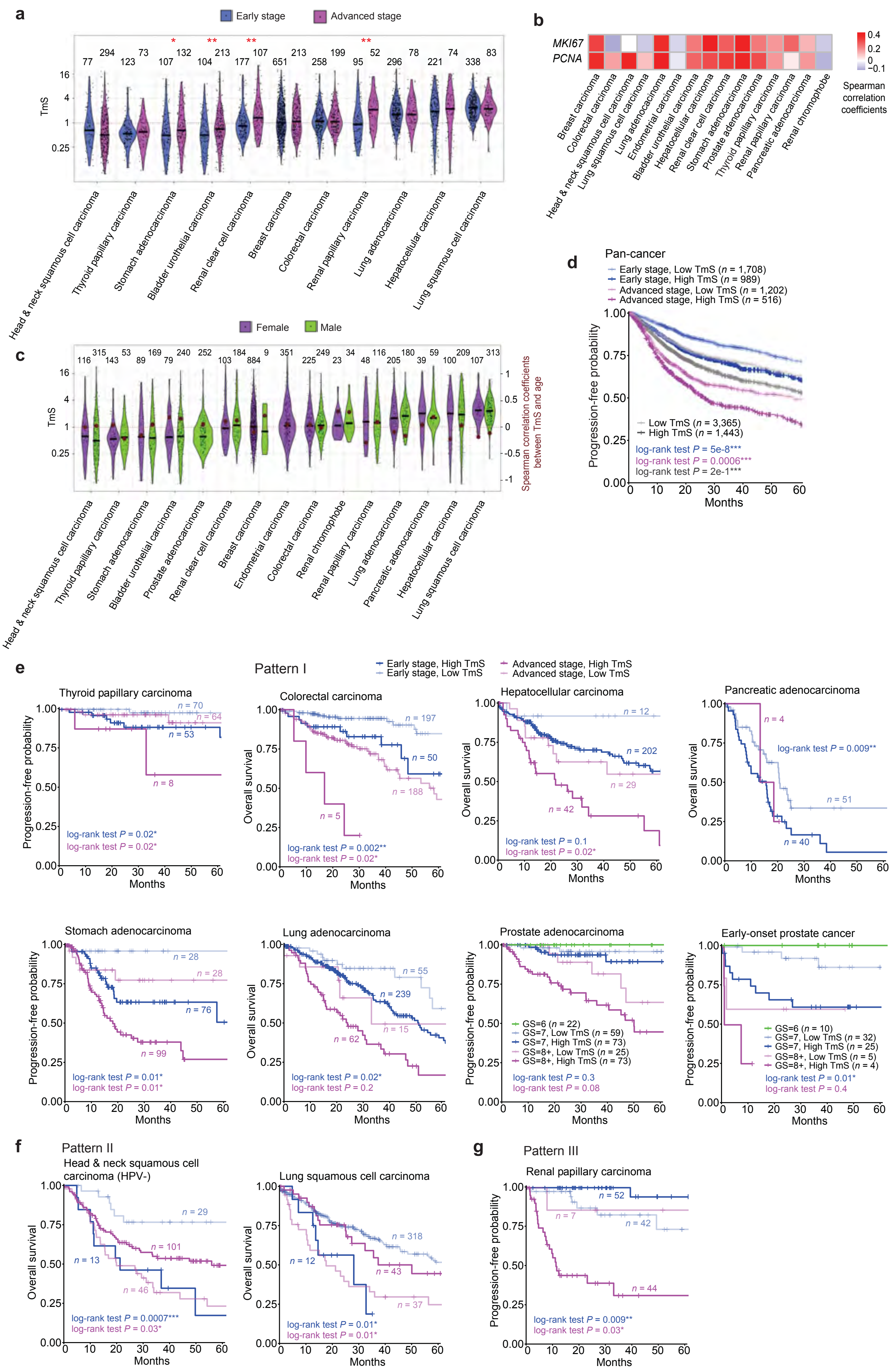

**Supplementary Figure. 5: TmS refines prognostication on pathological stages.**

**Supplementary Figure. 5 (cont'd): a**, Distribution of TmS for early (stage I and II) *vs*. advanced (stage III and IV) pathological stages across 15 cancer types. Adjusted *P* values of Wilcoxon rank-sum tests are indicated by asterisks (* $P < 0.05$, ** < 0.01, *** < 0.001). **b**, Spearman correlation coefficients between TmS and the expression levels of *MKI67* and *PCNA* across 15 cancer types. **c**, Distribution of TmS for female and male patient samples in TCGA across 15 cancer types. None of the adjusted *P* values of Wilcoxon rank-sum tests comparing TmS between the two groups reached significance at a confidence level of 0.05. Brown circles (read out on the right y-axis) represent Spearman correlation coefficients between TmS and age within the same sex and cancer type. The red dotted horizontal line represents TmS equal to 1 (left y axis) and correlation equal to 0 (right y axis). None of the adjusted *P* values for correlation tests reached significance at a confidence level of 0.05. **d**, KM curves of PFI for TCGA samples. Gray lines denote summary KM curves of patients with high *vs*. low TmS across all cancer types. KM curves are further grouped by TmS and pathological stage into four groups. *P* values of log-rank tests between high *vs*. low TmS groups are indicated by asterisks (* $P < 0.05$, ** < 0.01, *** < 0.001). **e**, KM survival curves for individual cancer types with pattern I. **f**, KM survival curves for individual cancer types with pattern II. **g**, KM survival curves for renal papillary carcinoma with pattern III.

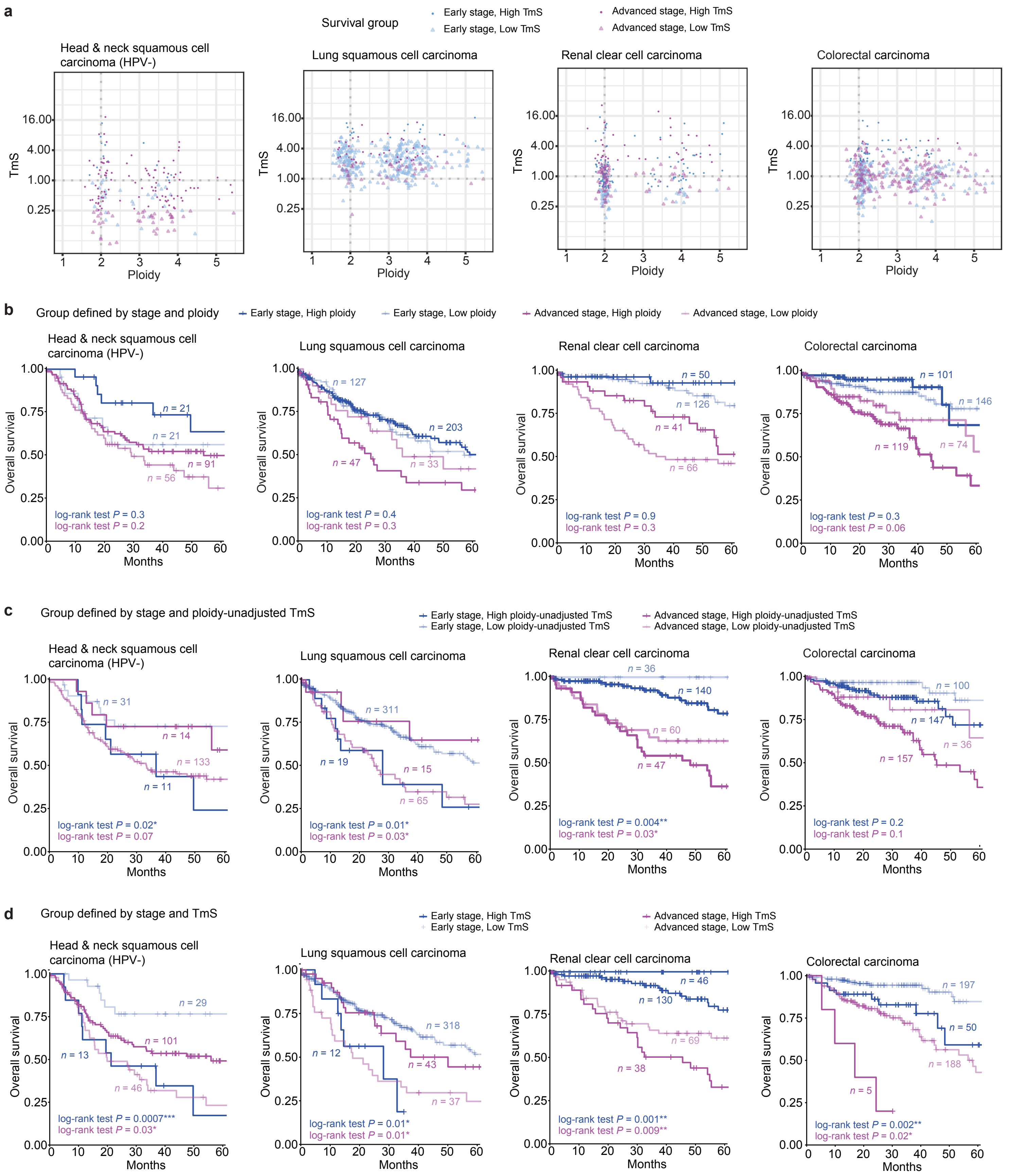


**Supplementary Figure. 6: Prognostication using ploidy or ploidy-unadjusted TmS on pathological stages.**

**Supplementary Figure. 6 (cont'd):** **a**, Scatter plots of TmS (y axis) *vs.* tumor ploidy (x axis) for samples from TCGA patient cohorts with head-and-neck squamous cell carcinoma (HPV negative), lung squamous cell carcinoma, renal clear cell carcinoma, and colorectal carcinoma. The samples were grouped into high *vs.* low TmS within early or advanced pathological stages, with different groups shown in distinct colors. **b**, KM survival curves of overall survival in four cancer types over patient groups defined by ploidy and stage. We grouped patients into high *vs.* low ploidy based on a cutoff of 2.5 within early or advanced pathological stage. **c**, KM survival curves of overall survival in four cancer types over patient groups defined by ploidy-unadjusted TmS and stage. **d**, KM survival curves of overall survival in four cancer types over patient groups defined by TmS and stage. *P* values of log-rank tests pairs of patient groups are shown with matching colors and are indicated by asterisk (* $P < 0.05$, ** < 0.01, *** < 0.001).

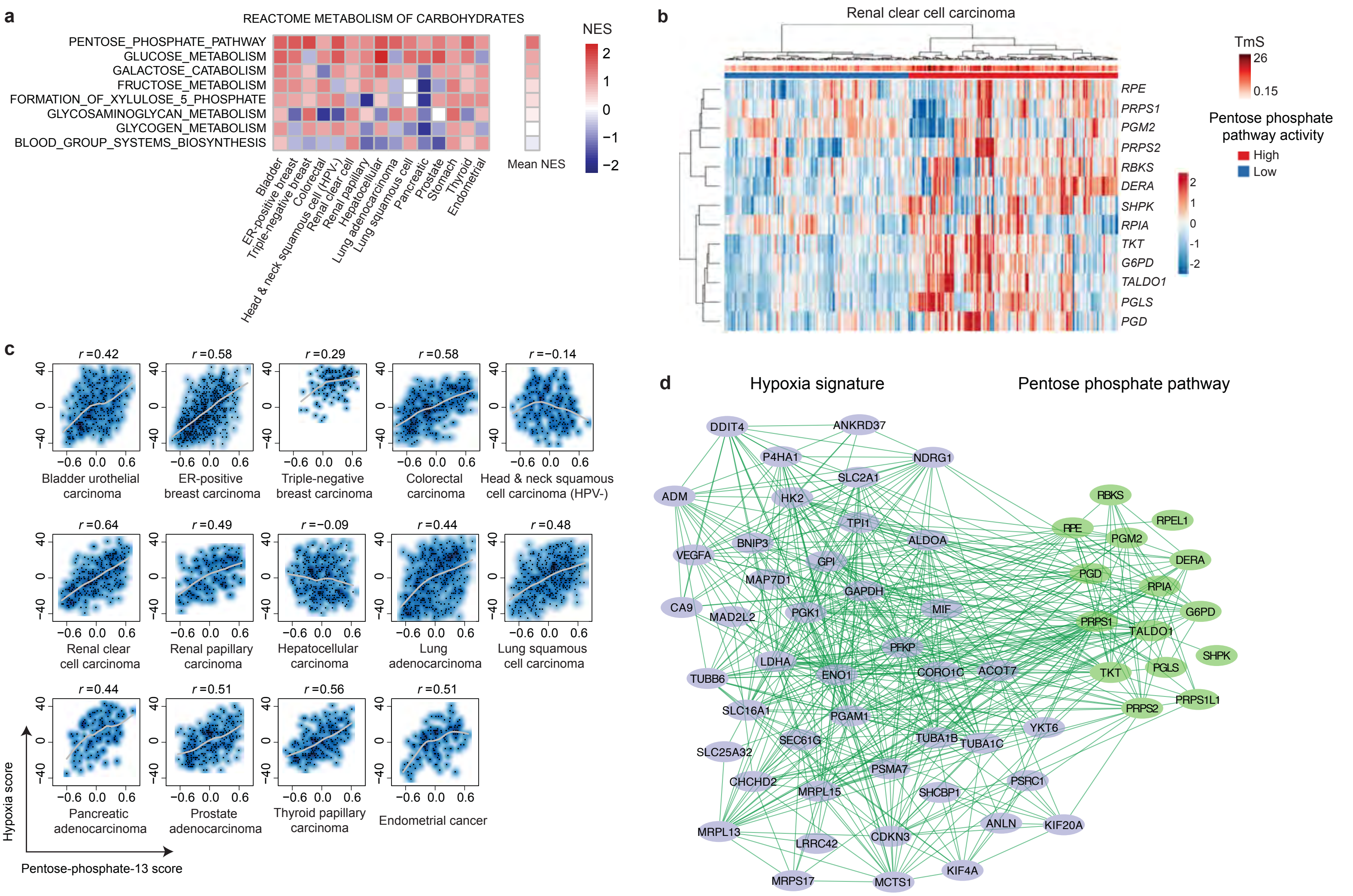


**Supplementary Figure. 7: Pan-cancer contributions of metabolic and hypoxia levels to TmS. a**, Heatmap of normalized enrichment scores (NES) of Reactome metabolism of carbohydrates pathways across 15 cancer types in TCGA. Pathways are ordered by the mean NES across 15 cancer types, from high to low. **b**, Hierarchical clustering on the expression levels of Reactome pentose phosphate pathway genes for the tumor samples from renal clear cell carcinoma. Samples were separated into two groups. **c**, Smoothed scatter plots showing correlations between pentose-phosphate-13 score and hypoxia score in each cancer type. Spearman correlation coefficients are labelled on top. **d**, Interaction network of genes from the pentose phosphate pathway and hypoxia signature. Each node represents a gene and each edge represents a known gene-gene interaction in the PCNet database.

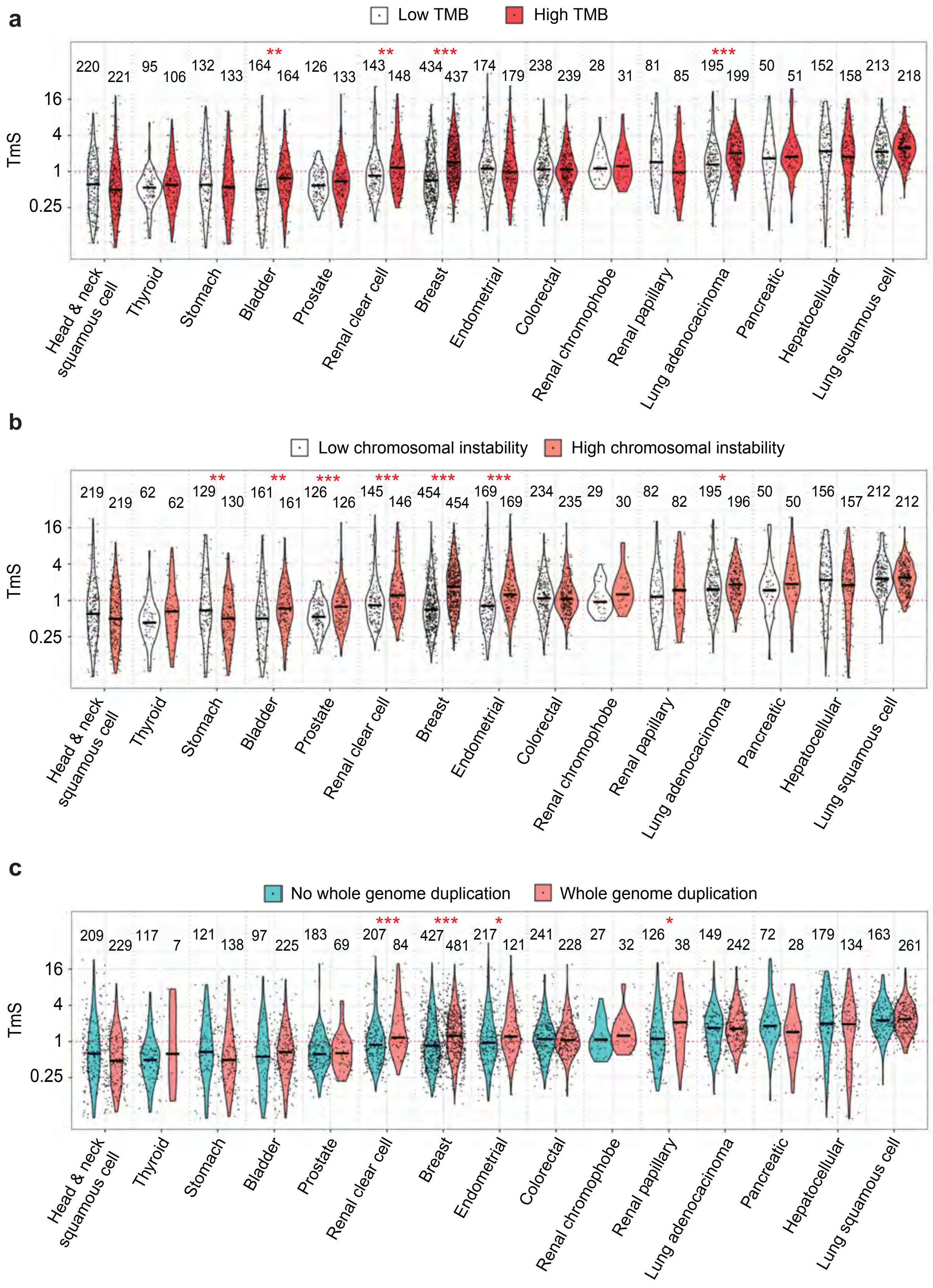


**Supplementary Figure. 8: TmS is associated with tumor genomic features across cancer types. a-c**, Distribution of TmS for patient samples with (**a**) high or low tumor mutation burden (TMB); (**b**) high or low chromosomal instability score; (**c**) with or without a whole genome duplication event. Patient groups are divided in half at the median TMB and chromosomal instability scores in (**a**) and (**b**) respectively. Adjusted *P* values of Wilcoxon rank-sum tests are indicated by asterisks (* $P < 0.05$, ** < 0.01, *** < 0.001).

**Supplementary Table 1 | Clinical information of the nine patients in the single cell RNA sequencing data analysis.**

| Cancer type | Patient id | Age | Sex | Treatment | Stage | Progression free survival (months) | Outcome | Tumor tissue collection | Details |
|---|---|---|---|---|---|---|---|---|---|
| Colorectal adenocarcinoma | Patient 1 | 45 | M | FOLFOX/ panitumumab | IVA | 4 | Relapsed with multiple tumors in liver | Surgical resection | Received 10 cycles of chemotherapy prior to surgery; regression grade 3 and 70% viable tumor; tumor is moderately differentiated; KRAS wild type |
| | Patient 2 | 63 | F | FOLFOX/bev | IVA | ≥ 15 | No tumor on last scan | Surgical resection | Received 4 cycles of chemotherapy prior to surgery; regression grade 3 and 90% viable tumor; tumor is moderately differentiated; KRAS mutation |
| Hepatocellular carcinoma | Patient 1 | 65 | M | Durvalumab/ tremelimumab | IV | < 5 | Alive but with decreased progression | Surgical resection | Etiology is Hepatitis C virus |
| | Patient 2 | 63 | M | Durvalumab/ tremelimumab | IV | ≥ 18 | Progression free | Surgical resection | Etiology is Hepatitis C virus |
| | Patient 3 | 74 | M | Treatment-naïve | I | ≥ 18 | Progression free | Core needle biopsy | NA |
| Lung adenocarcinoma | Patient 1 | 68 | M | Treatment-naïve | IIIB | NA | NA | Surgical resection | TNM stage: pT4N2M0 |
| | Patient 2 | 64 | F | Treatment-naïve | IIB | NA | NA | Surgical resection | TNM stage: pT2aN1M0 |
| Pancreatic adenocarcinoma | Patient 1 | 62 | F | Treatment-naïve | IV | 16 | Developed liver metastses | Fine needle aspiration | Alive; date of diagnosis is 12/18/2018; date of last follow up time is 5/6/2020; date of biopsy is 1/10/2019; developed liver metastases on 12/5/2019 |
| | Patient 2 | 70 | M | Treatment-naïve | IIB | ≥ 26 | Progression free | Fine needle aspiration | Alive; date of diagnosis is 1/7/2018; date of last follow up time is 3/23/2020; date of biopsy is 1/9/2019; date of surgery is 5/31/2019 |

**Supplementary Table 2 | Single-cell gene set enrichment analyses for high and low UMI tumor cells in each of the "patient 1" samples with colorectal, lung and pancreatic cancers.**

Within each data sheet, gene sets are sorted by adjusted *P* values from the low to high for both high UMI tumor cells (left) and low UMI tumor cells (right). Significantly enriched gene sets (adjusted *P* value < 0.1) are highlighted in yellow. The enriched gene sets that support EMT or deviation from the tissue origin are highlighted in brown. Adjusted *P* values are calculated with the BH correction.

| Sheet name | Patient sample | Gene sets investigated (size) | Number of significantly enriched gene sets |
|---|---|---|---|
| Stemness enrichment colorectal | Colorectal cancer patient 1 | Stemness signatures (314) | 40 for high UMI tumor cells<br>0 for low UMI tumor cells |
| Stemness enrichment lung | Lung cancer patient 1 | Stemness signatures (314) | 14 for high UMI tumor cells<br>2 for low UMI tumor cells |
| Stemness enrichment pancreatic | Pancreatic cancer patient 1 | Stemness signatures (314) | 16 for high UMI tumor cells<br>0 for low UMI tumor cells |
| All enrichment colorectal | Colorectal cancer patient 1 | All gene sets (18,617) | 2,124 for high UMI tumor cells<br>0 for low UMI tumor cells |
| All enrichment lung | Lung cancer patient 1 | All gene sets (18,617) | 395 for high UMI tumor cells<br>370 for low UMI tumor cells |
| All enrichment pancreatic | Pancreatic cancer patient 1 | All gene sets (18,617) | 961 for high UMI tumor cells<br>0 for low UMI tumor cells |

The full table can be accessed at: https://drive.google.com/drive/folders/1XSCy_3Lv8yJxu6s2dBfNfsI2k7xd4mjX

### Supplementary Table 3 | Benchmarking results using a mixed cell line experiment.

| Sample | H1092 (true TmS = 0.87) | | | | CAF (true TmS = 1.2) | | | |
|---|---|---|---|---|---|---|---|---|
| | Cell count | Cell count proportion | Estimated mRNA proportion | Estimated TmS | Cell count | Cell count proportion | Estimated mRNA proportion | Estimated TmS |
| 1 | 48,000 | 0.51 | 0.47 | 0.86 | 46,600 | 0.49 | 0.55 | 1.25 |
| 2 | 48,000 | 0.51 | 0.51 | 1.00 | 46,600 | 0.49 | 0.53 | 1.16 |
| 3 | 48,000 | 0.51 | 0.47 | 0.85 | 46,600 | 0.49 | 0.55 | 1.27 |
| 4 | 58,124 | 0.67 | 0.62 | 0.81 | 29,050 | 0.33 | 0.38 | 1.23 |
| 5 | 58,124 | 0.67 | 0.61 | 0.77 | 29,050 | 0.33 | 0.39 | 1.30 |
| 6 | 58,124 | 0.67 | 0.61 | 0.77 | 29,050 | 0.33 | 0.39 | 1.28 |
| 7 | 36,075 | 0.33 | 0.39 | 1.26 | 72,100 | 0.67 | 0.65 | 0.94 |
| 8 | 36,075 | 0.33 | 0.34 | 1.04 | 72,100 | 0.67 | 0.68 | 1.06 |
| 9 | 36,075 | 0.33 | 0.38 | 1.23 | 72,100 | 0.67 | 0.66 | 0.98 |
| 10 | 59,399 | 0.67 | 0.60 | 0.76 | 29,700 | 0.33 | 0.38 | 1.21 |
| 11 | 59,399 | 0.67 | 0.61 | 0.78 | 29,700 | 0.33 | 0.39 | 1.28 |
| 12 | 59,399 | 0.67 | 0.64 | 0.89 | 29,700 | 0.33 | 0.38 | 1.22 |
| 13 | 36,975 | 0.33 | 0.37 | 1.17 | 74,000 | 0.67 | 0.68 | 1.05 |
| 14 | 36,975 | 0.33 | 0.38 | 1.21 | 74,000 | 0.67 | 0.66 | 0.98 |
| 15 | 36,975 | 0.33 | 0.38 | 1.24 | 74,000 | 0.67 | 0.66 | 0.96 |
| 16 | 68,249 | 0.84 | 0.76 | 0.60 | 12,800 | 0.16 | 0.24 | 1.66 |
| 17 | 68,249 | 0.84 | 0.76 | 0.59 | 12,800 | 0.16 | 0.24 | 1.68 |
| 18 | 68,249 | 0.84 | 0.76 | 0.58 | 12,800 | 0.16 | 0.24 | 1.70 |
| Median of estimated TmS | 0.86 | | | | 1.2 | | | |
| Median absolute deviation (MAD) of estimated TmS | 0.24 | | | | 0.18 | | | |

**Supplementary Table 4 | Summary of the distributions of TmS across 15 cancer types in TCGA and ICGC-EOPC.**

| Cancer | Median TmS | MAD TmS | Sample size |
|---|---|---|---|
| Head-and-neck squamous cell carcinoma | 0.51 | 0.51 | 431 |
| Thyroid papillary carcinoma | 0.54 | 0.32 | 196 |
| Stomach adenocarcinoma | 0.56 | 0.52 | 258 |
| Bladder urothelial carcinoma | 0.60 | 0.52 | 319 |
| Prostate adenocarcinoma | 0.61 | 0.39 | 252 |
| Renal clear cell carcinoma | 1.0 | 0.67 | 287 |
| Early-onset prostate cancer | 1.0 | 0.84 | 85 |
| Breast carcinoma | 1.0 | 0.85 | 893 |
| Endometrial carcinoma | 1.0 | 0.80 | 351 |
| Colorectal carcinoma | 1.1 | 0.65 | 477 |
| Renal chromophobe | 1.1 | 0.59 | 57 |
| Renal papillary carcinoma | 1.2 | 1.0 | 164 |
| Lung adenocarcinoma | 1.6 | 1.1 | 385 |
| Pancreatic adenocarcinoma | 1.6 | 1.1 | 98 |
| Hepatocellular carcinoma | 1.9 | 2.1 | 309 |
| Lung squamous cell carcinoma | 2.3 | 1.3 | 420 |
| Overall | 1.0 | 0.91 | 4,982 |

MAD: Median absolute deviation

**Supplementary Table 5 | Multivariate Cox proportional hazard models with Age, TmS, Stage and TmS x Stage as predictors for overall survival and progression-free interval analysis across cancer types in TCGA and ICGC-EOPC.**

| Cancer | Sample size | Clinical Outcome Endpoint | Variable | Hazard ratio | 95% CI | P value (Wald test) | TmS prognostication type |
|---|---|---|---|---|---|---|---|
| Colorectal carcinoma | *n*=440 | OS | Age | 1.04 | (1.03, 1.06) | $5x10^{-6}$ | Pattern I |
| | | | TmS (High vs. Low) | 2.9 | (1.4, 6.2) | 0.01 | |
| | | | Stage (Advanced vs. Early) | 5.1 | (2.9, 8.8) | $1x10^{-8}$ | |
| | | | TmS x Stage | 0.96 | (0.26, 3.6) | 1 | |
| | | PFI | Age | 1.0 | (0.98, 1.02) | 1 | |
| | | | TmS (High vs. Low in Advanced Stage) | 2.4 | (0.97, 6.0) | 0.06 | |
| | | | Stage/(TmS x Stage) | - | - | - | |
| Hepatocellular carcinoma | *n*=285 | OS | Age | 1.0 | (1.00, 1.03) | 0.09 | |
| | | | TmS (High vs. Low) | 3.7 | (0.51, 26) | 0.2 | |
| | | | Stage (Advanced vs. Early) | 5.1 | (0.67, 39) | 0.1 | |
| | | | TmS x Stage | 0.62 | (0.078, 5.1) | 0.7 | |
| | | PFI | Age | 1.0 | (0.98, 1.0) | 0.7 | |
| | | | TmS (High vs. Low in Advanced Stage) | 1.8 | (0.83, 3.7) | 0.1 | |
| | | | Stage/(TmS x Stage) | - | - | - | |
| Lung adenocarcinoma | *n*=371 | OS | Age | 1.0 | (0.99, 1.03) | 0.2 | |
| | | | TmS (High vs. Low) | 2.1 | (1.2, 4.0) | 0.02 | |
| | | | Stage (Advanced vs. Early) | 2.3 | (0.86, 6.3) | 0.1 | |
| | | | TmS x Stage | 0.99 | (0.34, 2.9) | 1.0 | |
| | | PFI | Age | 1.0 | (0.98, 1.01) | 0.8 | |
| | | | TmS (High vs. Low) | 2.9 | (0.93, 9.2) | 0.1 | |
| | | | Stage (Advanced vs. Early) | 3.6 | (1.1, 11.7) | 0.04 | |
| | | | TmS x Stage | 1.2 | (0.23, 6.4) | 0.8 | |
| Panceatic adenocarcinoma | *n*=95 | OS | Age | 1.0 | (0.99, 1.0) | 0.3 | |
| | | | TmS (High vs. Low in Early Stage) | 2.0 | (1.2, 3.4) | 0.01 | |
| | | | Stage/(TmS x Stage) | - | - | - | |
| | | PFI | Age | 1.0 | (0.98, 1.0) | 0.8 | |
| | | | TmS (High vs. Low in Early Stage) | 6.0 | (1.7, 20) | 0.004 | |
| | | | Stage/(TmS x Stage) | - | - | - | |
| Renal clear cell carcinoma | *n*=283 | OS | Age | 1.0 | (1.02, 1.06) | 0.0002 | |
| | | | TmS (High vs. Low) | 2.3 | (1.4, 4.0) | 0.002 | |
| | | | Stage (Advanced vs. Early) | 4.5 | (2.6, 7.7) | $5x10^{-8}$ | |
| | | | TmS x Stage | - | - | - | |
| | | PFI | Age | 1.01 | (0.99, 1.0) | 0.3 | |
| | | | TmS (High vs. Low) | 2.6 | (1.5, 4.3) | 0.0004 | |
| | | | Stage (Advanced vs. Early) | 9.7 | (5.5, 17) | $1x10^{-15}$ | |
| | | | TmS x Stage | - | - | - | |
| Stomach adenocarcinoma | *n*=231 | OS | Age | 1.0 | (1.00, 1.05) | 0.03 | |
| | | | TmS (High vs. Low) | 2.0 | (0.85, 4.6) | 0.1 | |
| | | | Stage (Advanced vs. Early) | 1.1 | (0.23, 5.1) | 0.9 | |
| | | | TmS x Stage | 2.0 | (0.38, 10) | 0.4 | |
| | | PFI | Age | 0.99 | (0.97, 1.01) | 0.5 | |
| | | | TmS (High vs. Low) | 8.3 | (1.1, 62) | 0.04 | |
| | | | Stage (Advanced vs. Early) | 5.0 | (0.59, 43) | 0.1 | |
| | | | TmS x Stage | 0.41 | (0.045, 3.7) | 0.4 | |
| Thyroid papillary carcinoma | *n*=195 | PFI | Age | 1.0 | (0.96, 1.06) | 0.8 | |
| | | | TmS (High vs. Low) | 7.3 | (0.88, 61) | 0.07 | |
| | | | Stage (Advanced vs. Early) | 3.6 | (0.32, 40) | 0.3 | |
| | | | TmS x Stage | 1.2 | (0.083, 16) | 0.9 | |
| Prostate adenocarcinoma | *n*=230 | PFI | Age | 1.0 | (0.97, 1.06) | 0.6 | |
| | | | TmS (High vs. Low) | 2.5 | (0.48, 13) | 0.3 | |

| | | | | | | | |
|---|---|---|---|---|---|---|---|
| | | | Gleason score (7 vs. 8+) | 8.0 | (1.7, 39) | 0.01 | |
| | | | TmS x Gleason score | 0.81 | (0.13, 5.1) | 0.8 | |
| Early-onset prostate cancer | *n*=66 | PFI | Age | 0.93 | (0.79, 1.09) | 0.4 | |
| | | | TmS (High vs. Low) | 4.3 | (1.2, 16) | 0.03 | |
| | | | Gleason score (7 vs. 8+) | 6.8 | (1.1, 42) | 0.04 | |
| | | | TmS x Gleason score | 0.87 | (0.090, 8.3) | 0.9 | |
| Bladder urothelial carcinoma | *n*=317 | OS | Age | 1.0 | (1.0, 1.1) | 0.04 | Pattern II |
| | | | TmS (High vs. Low) | - | - | - | |
| | | | Stage/(TmS x Stage) | - | - | - | |
| | | PFI | Age | 1.0 | (0.99, 1.02) | 0.4 | |
| | | | TmS (High vs. Low) | 3.3 | (0.79, 14) | 0.1 | |
| | | | Stage (Advanced vs. Early) | 9.1 | (2.2, 37) | 0.002 | |
| | | | TmS x Stage | 0.18 | (0.04, 0.79) | 0.02 | |
| Head-and-neck squamous cell carcinoma (HPV-) | *n*=189 | OS | Age | 1.0 | (1.0, 1.1) | 0.001 | |
| | | | TmS (High vs. Low) | 4.4 | (1.6, 12) | 0.003 | |
| | | | Stage (Advanced vs. Early) | 5.4 | (2.3, 12) | $9x10^{-5}$ | |
| | | | TmS x Stage | 0.14 | (0.047, 0.40) | 0.0003 | |
| | | PFI | Age | 1.0 | (0.99, 1.0) | 0.4 | |
| | | | TmS (High vs. Low) | 0.55 | (0.34, 0.91) | 0.02 | |
| | | | Stage (Advanced vs. Early) | 1.4 | (0.75, 2.7) | 0.3 | |
| | | | TmS x Stage | - | - | - | |
| Lung squamous cell carcinoma | *n*=410 | OS | Age | 1.0 | (1.0, 1.03) | 0.06 | |
| | | | TmS (High vs. Low) | 2.8 | (1.4, 5.8) | 0.005 | |
| | | | Stage (Advanced vs. Early) | 2.6 | (1.7, 4.0) | $7x10^{-6}$ | |
| | | | TmS x Stage | 0.16 | (0.064, 0.42) | 0.0002 | |
| | | PFI | Age | 1.0 | (0.98, 1.0) | 0.7 | |
| | | | TmS (High vs. Low) | 2.2 | (1.0, 4.5) | 0.04 | |
| | | | Stage (Advanced vs. Early) | 2.2 | (1.5, 3.4) | 0.0001 | |
| | | | TmS x Stage | - | - | - | |
| Triple-negative breast carcinoma† | *n*=127 | OS | Age | 1.0 | (0.97, 1.0) | 0.9 | Pattern III |
| | | | TmS + Stage | 6.3 | (2.6, 15) | $6x10^{-5}$ | |
| | | PFI | Age | 1.0 | (0.96, 1.03) | 0.5 | |
| | | | TmS + Stage | 6.4 | (2.7, 15) | $2x10^{-5}$ | |
| Renal papillary carcinoma† | *n*=145 | PFI | Age | 0.97 | (0.94, 1.0) | 0.02 | |
| | | | TmS + Stage | 7.6 | (3.0, 19) | $1x10^{-5}$ | |

OS: Overall survival; PFI: Progression-free interval.

"-" stands for missing coefficients due to lack of events (death or progression) in the corresponding patient group.

For the cancer types with "†", hazard ratios of TmS and Stage cannot be estimated separately due to lack of events. Instead, a simplified model comparing "Early stage, Low TmS" and "Advanced stage, High TmS" was used and denoted as "TmS + Stage".
