## Supplementary Note for "Tumor cell total mRNA expression shapes the molecular and clinical phenotype of cancer"

### 1. TOTAL MRNA EXPRESSION IN SINGLE-CELL RNA SEQUENCING DATA

#### 1.1. Datasets

*Colorectal cancer single-cell RNA sequencing data*

Two fresh colorectal adenocarcinoma samples of primary tumor were collected from patients who were receiving chemotherapies by surgical resection at the University of Texas MD Anderson Cancer Center (**Supplementary Table 1**). Single-cell data was generated using the Chromium Single Cell 3' Library, Gel Bead & Multiplex Kit, and Chip Kit (v3, 10x Genomics). Libraries were sequenced on an Illumina NovaSeq6000. Alignment, tagging, and gene and transcript counting were conducted using the 10x Genomic Cell Ranger pipeline (version 3.0).

*Liver cancer single-cell RNA sequencing data*[1]

Three fresh hepatocellular carcinoma samples of primary tumor were collected at the NIH Clinical Center for immune checkpoint inhibition studies (NCT01313442) (**Supplementary Table 1**). Two of them (patient 1 and patient 2) were from patients who were receiving immunotherapies by needle biopsy, and the other was collected from an untreated patient by surgical resection. Single-cell data was generated using the Chromium Single Cell 3' Library, Gel Bead & Multiplex Kit, and Chip Kit (v2, 10x Genomics). Libraries were sequenced on an Illumina NextSeq500. Alignment, tagging, and gene and transcript counting were conducted using the 10x Genomic Cell Ranger pipeline (version 2.0.2).

*Lung cancer single-cell RNA sequencing data*[2]

Two fresh lung adenocarcinoma samples of primary, non-metastatic lung tumor were collected from untreated patients by surgical resection at University Hospital Leuven (**Supplementary Table 1**). Single-cell data was generated using the Chromium Single Cell 3' Library, Gel Bead & Multiplex Kit, and Chip Kit (v1, 10x Genomics). Libraries were sequenced on Illumina HiSeq4000. Alignment, tagging, and gene and transcript counting were conducted using the 10x Genomic Cell Ranger pipeline (version 2.0.0).

*Pancreatic cancer single-cell RNA sequencing data*[3]

Two untreated patients with primary pancreatic cancer were recruited at the University of Texas MD Anderson Cancer Center and informed written consents following institutional review board approval were obtained (Lab00-396 and PA15-0014). Fresh biopsies were collected from the tumors by fine needle aspiration (**Supplementary Table 1**). Single-cell data was generated using the Chromium Single Cell 3' Library, Gel Bead & Multiplex Kit, and Chip Kit (v1, 10x Genomics). Libraries were sequenced on an Illumina NextSeq500. Alignment, tagging, and gene and transcript counting were conducted by using the 10x Genomic Cell Ranger pipeline (version 3.1).

*LNCaP prostate cancer cell line single-cell RNA sequencing data*

LNCaP cells were provided by Prof. Frank Claessens, KU Leuven[4], and cultured in RPMI 1640 (Sigma R0883) supplemented with 10% FBS (Sigma F7524), 2 mM Alanyl-glutamine (Sigma G8541), 1 mM sodium pyruvate (Merck TMS-005-C), 2.5 g/L glucose (Sigma G8769), and 1x Antibiotic-Antimycotic (Gibco, 15240062) in a humidified 37°C incubator with 5% $CO_2$. For experimental treatments, ~1x$10^6$ cells were seeded into 5 cm culture plate dishes, and allowed to settle for 2 days before exposure to 10 µM Enzalutamide (MedChemExpress HY-70002) or DMSO vehicle control (0.1%) for 48 hours. Cells were then harvested with 0.05% Trypsin-EDTA (Sigma T3924). After neutralization with complete medium, centrifugation (300 x g for 5 min) and resuspension in PBS/0.5%BSA, the cells were filtered through a 35 µm Cell Strainer (Corning 352235), and a single-cell suspension of living cells was acquired through sorting on a FACS Aria II cell sorter. The cell concentration of the single-cell suspension was assessed with a Countess II FL Automated Cell Counter, and ~3 x $10^4$ cells were pelleted (300 x g for 5 min). Single-cell data was generated using the Chromium Single Cell 3' Library, Gel Bead & Multiplex Kit, and Chip Kit (v1, 10x Genomics). Libraries were sequenced on an Illumina NextSeq 500 (scRNAseq kit v3). Sequencing reads were processed into FASTQ format and single cell feature counts using Cell Ranger (version 3.0.2).

##### 1.2. Single-cell RNA sequencing data processing

In this section, we first introduce the preprocessing for the single-cell RNA sequencing (scRNAseq) datasets described above (including quality control, cell clustering, cell type annotation), followed by the introduction of a method to group cell clusters within a cell type based on gene counts, *i.e.*, the total number of expressed genes, to simplify the characterization of heterogeneity within the cell type. We then introduce a scale normalization method to correct for sequencing or experimental biases on total UMI counts, and finally the trajectory and cell cycle analyses for the scRNAseq data.

**Supplementary Note Table 1. Marker genes used to annotate cell types in scRNAseq patient samples from four cancer types.**

| | Colorectal adenocarcinoma[6] | Hepatocellular carcinoma[1] | Lung adenocarcinoma[2] | Pancreatic adenocarcinoma[3,8,9] |
|---|---|---|---|---|
| B cell | *CD79A*, *CD38* | *CD79A*, *SLAMF7*, *BLNK* | *CD79A*, *IGKC*, *IGLC3* | *CD79A*, *CD38* |
| T cell | *CD2*, *CD3E*, *CD3D* | *CD2*, *CD3E*, *CD3D* | *CD3D*, *TRBC1*, *TRBC2* | *CD2*, *CD3D* |
| NK cell | | | | *NKG7*, *KLRF1* |
| Myeloid | *CD14*, *CD68*, *ITGAX* | *CD14*, *CD163*, *CD68* | *LYZ*, *MARCO*, *CD68* | *CD14*, *CD68* |
| Fibroblast | *COLA1A*, *COL1A2*, *COL3A1* | *COL1A2*, *FAP*, *PDPN* | *COLA1A*, *DCN*, *COL1A2* | *COLA1A*, *COL1A2* |
| Endothelial | *PECAM1*, *VWF*, *ENG* | *PECAM1*, *VWF*, *ENG* | *CLDN5*, *FLT1*, *CDH5* | |
| Alveolar | | | *FOLR1*, *AQP4*, *PEBP4* | |
| Epithelial | *EPCAM*, *KRT18*, *KRT20* | | *CAPS*, *TEME190*, *PIFO*, *SNTN* | *EPCAM*, *KRT18*, *KRT20* |
| Tumor cell | | | *LCN2*, *CCL20*, *PTTG1* | |

##### 1.2.1. Quality control, clustering and cell type annotation

For each of the two colorectal adenocarcinoma scRNAseq samples generated at MD Anderson, genes expressed in less than three cells were removed. Cells with either fewer than 500 total UMIs, below 200 expressed genes, or more than 50% total UMI counts derived from mitochondrial genes were excluded. The total number of transcripts in each cell was normalized to 10,000, which was followed by a natural log transformation. Highly variable genes were detected and used for principal component analysis (PCA). Cells were then clustered with the Seurat package[5]. The cell type for each cell was annotated based on known marker genes[6] (**Supplementary Note Figure 1**, **Supplementary Note Table 1**). Initial somatic copy number variation (CNV) estimates were made using inferCNV[7], which was used to calculate CNV scores and CNV correlation scores[1]. The CNV score of a single cell was defined as the sum of the squared copy number variants across all gene positions. The CNV correlation score was calculated as the correlation between the copy number variations of a single cell and the average copy number variation of the top 2% cells ranked by CNV scores from the same sample. Tumor cells were identified as epithelial cells with an average CNV score greater than 0.0015. The two samples from patient 1 and patient 2 had 5,422 and 7,462 cells remaining, respectively, after data pre-processing.

The quality control of the three hepatocellular carcinoma scRNAseq patient samples was conducted following the method described by Ma, L. *et al*[1]. For each sample, genes expressed in less than 0.1% of the cells were removed. Cells with fewer than 700 total UMIs, fewer than 500 expressed genes, or more than 20% total UMI counts derived from mitochondrial genes were excluded. An additional quality control step of doublet removal was performed based on the number of cells loaded and recovered. The total number of transcripts in each cell was normalized to 10,000, followed by a natural log transformation. Highly variable genes were detected and used for PCA. Cells were then clustered with the Seurat package[5]. The cell type for each cell was annotated based on known marker genes[1] (**Supplementary Note Figure 1**, **Supplementary Note Table 1**). Tumor cells were identified as epithelial cells with CNV scores above the 80th percentile and CNV correlation scores above 0.4. The three samples of patient 1, patient 2 and patient 3 had 83, 761 and 796 cells remaining, respectively, after data pre-processing.

The quality control of the two lung adenocarcinoma scRNAseq patient samples was conducted following the method described by Lambrechts, D. *et al*[2]. For each sample, genes expressed in less than 0.5% of the cells were removed. Any cell with either fewer than 201 total UMI counts, below 101 or over 6,000 expressed genes, or more than 10% total UMI counts derived from mitochondrial genes were filtered out from downstream analysis. The total number of transcripts in each cell was normalized to 10,000, followed by a natural log transformation. Highly variable genes were detected and used for PCA. Cells were then clustered with the Seurat package[5]. Cell type (including tumor cell) of each cell was annotated based on known marker genes[2] (**Supplementary Note Figure 1**, **Supplementary Note Table 1**). The two samples (patient 1 and patient 2) had 8,845 and 13,658 cells remaining, respectively, after data pre-processing.

For each of the two pancreatic adenocarcinoma scRNAseq samples, genes expressed in less than three cells were removed. Cells with either fewer than 500 total UMIs, below 200 expressed genes, or more than 50% total UMI counts derived from mitochondrial genes were filtered out. The total number of transcripts in each cell was normalized to 10,000, followed by a natural log transformation. Highly variable genes were detected and used for PCA. Cells were then clustered with the Seurat package[5]. Cell type of each cell was annotated based on known marker genes[8,9] (**Supplementary Note Figure 1**, **Supplementary Note Table 1**). Tumor cells were identified as epithelial cells with CNV score above 0.015 and CNV correlation above 0.4. The two samples (patient 1 and patient 2) had 2,404 and 7,037 cells remaining after QC, respectively, after data pre-processing.

Within each cell type, we further merged clusters that did not significantly differ in gene counts (Wilcoxon rank-sum test, α=0.001, **Supplementary Note Figure 2**).

For the two LNCaP scRNAseq samples, cells were treated either with Enzalutamide for 48 hours (treated sample) or with DMSO control for 48 hours (untreated sample). Genes expressed in fewer than three cells were removed. Poor quality cells were filtered based on the number of detected genes, the total number of molecules detected in each cell, and the percentage of reads arising from the mitochondrial genome. Specifically, any cell with either fewer than 500 total UMI counts, below 200 expressed genes, or more than 15% reads derived from mitochondrial genes were filtered out. Doublets (145 and 36 for treated and untreated sample, respectively) were detected by DoubletFinder[10] with default settings and removed from downstream analysis.

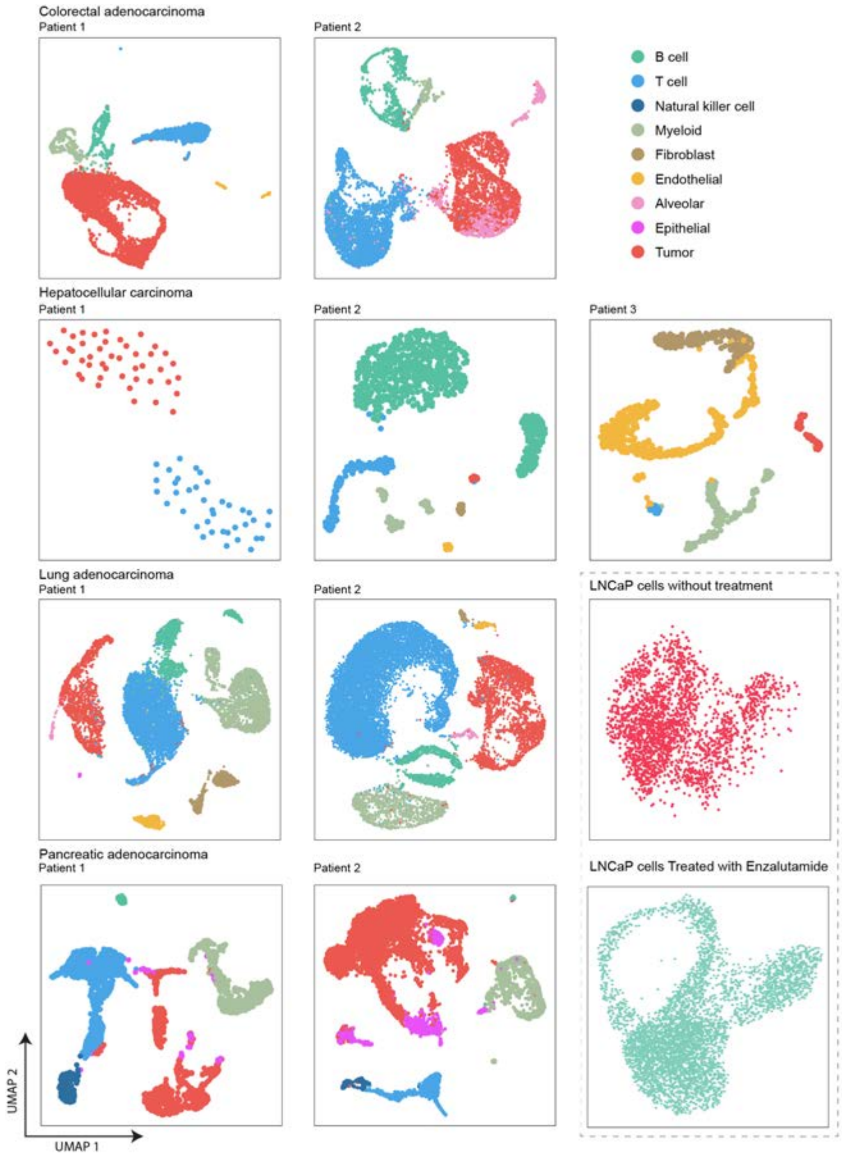

**Supplementary Note Figure 1. UMAPs of scRNAseq data from four cancer types and the LNCaP cell line data.**

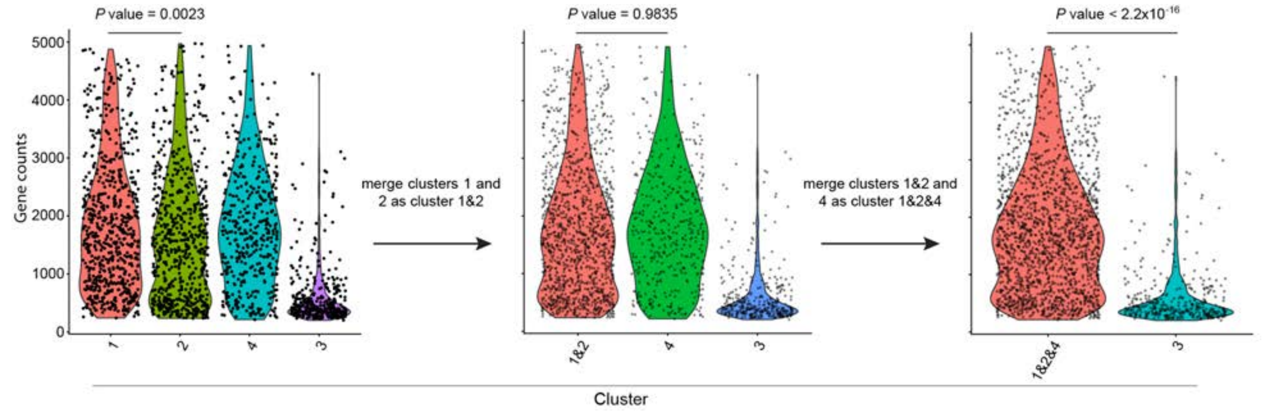

**Supplementary Note Figure 2**. **An example of merging cell clusters by gene counts.** Tumor cells in patient 2 of colorectal adenocarcinoma are used. The initial 4 clusters were determined by Seurat clustering (resolution=0.5). Wilcoxon rank-sum tests comparing gene counts were performed between clusters and those that did not pass the significance level of 0.001 were merged. The resulting two tumor cell clusters had 1,696 cells (low UMI cluster, e.g. 1, 2, and 4) and 359 cells (high UMI cluster, e.g. 3), respectively. We repeated this process based on the initial Seurat clustering with resolution=1.0. There were still two tumor cell clusters after merging. The differences of tumor cells in the high UMI cluster and in the low UMI cluster based on the two resolutions were only 12 cells and 13 cells, respectively.

##### 1.2.2. Normalized total UMI counts

We performed scale normalization on the raw count data to ensure the total UMI count per cell across all cells are comparable for different samples. Specifically, let $UMI_i = \{UMI_{igc}\}_{GxC_i}$ be a matrix of raw UMI counts for the scRNAseq data for sample *i* being investigated, with genes *g* on the rows and cells *c* on the columns. *G* denotes the total number of genes, $C_i$ is the number of cells in sample *i*. Then, the normalized UMI matrix $UMI_i$, denoted as $UMI_i^{norm}$, is calculated as $UMI_i^{norm}=UMI_i/r_i$, where, $r_i = \frac{UMI_i^{sum}/C_i}{baseline}$, $baseline = median\{UMI_1^{sum}/C_1, UMI_2^{sum}/C_2, \ldots, UMI_n^{sum}/C_n\}$, $UMI_i^{sum} = \sum_{c=1}^{C_i} \sum_{g=1}^{G} UMI_{igc}$.

Given a cell cluster, we let $u_{gc}$ denote the amount of mRNA of gene *g* in cell *c*. The average total mRNA amount per cell is $\sum_{c=1}^{C} (\sum_{g=1}^{G} u_{gc})/C$. For scRNAseq data, we assume the $UMI_{gc}$ from gene *g*, cell *c* is proportional to the total mRNA $u_{gc}$ of gene *g* in that cell, with a constant $k_g$ that represents technical effects: $UMI_{gc}=k_g * u_{gc}$. The constant $k_g$ is introduced because every single-cell sequencing platform presents a <100% capture efficiency for mRNA, and such efficiency varies across different platforms[11]. Under the assumption that the technical effect $k_g$ remains constant across cells and is often evaluated as an average effect across genes within the same platform, we can evaluate total mRNA expression in the scRNAseq data using the average total UMI counts, which is $\sum_{c=1}^{C} (\sum_{g=1}^{G} UMI_{gc})/C$. Notably, we observed strong correlations between gene counts and total UMI across cells in each cell cluster across all cell types and cancer types (**Supplementary Note Figure 3**). This observation supports our assumption of a stable technical effect $k_g$ within each study, and that the average total UMI counts serve as a reasonable surrogate to compare total mRNA expression across cells that are generated from the same experiment.

The average gene counts and average total UMI counts for both individual cell clusters and all the clusters pooled within a cell type are summarized in **Supplementary Note Table 2**.

The observed fold changes in total UMI counts between tumor cell clusters were significantly higher than those expected from expression dosage response from genome ploidy changes alone (at 2-3 folds[12,13]) among tumor cells (**Supplementary Note Table 3**). For the two tumor cell clusters in each patient across four cancer types, our null hypothesis is that there is no difference between the distribution of the total UMI counts from the tumor cell high-UMI cluster and the distribution of the total UMI counts from the tumor cell low-UMI cluster multiplied by three. For each patient, the *P* value was obtained with a t-test and adjusted by the Benjamini-Hochberg (BH) method[14]; the 95% confidence intervals were calculated using bootstrapping with 1,000 iterations.

**Supplementary Note Table 2. The average gene counts and average total UMI counts for both individual cell clusters and all the clusters pooled.** The 95% CI was estimated using bootstrapping with 1,000 iterations.

| Cancer type | Patient id | Cell cluster | Cell type | | | | | | | | | |
|---|---|---|---|---|---|---|---|---|---|---|---|---|
| | | | Tumor | | Epithelial | | Alveolar | | Endothelial | | Fibroblast | |
| | | | Average gene counts (95% CI) | Average total UMI counts (95% CI) | Average gene counts (95% CI) | Average total UMI counts (95% CI) | Average gene counts (95% CI) | Average total UMI counts (95% CI) | Average gene counts (95% CI) | Average total UMI counts (95% CI) | Average gene counts (95% CI) | Average total UMI counts (95% CI) |
| Colorectal adenocarcinoma | Patient 1 (Stage IVA, PFS = 4 months) | Cluster 3 | 5,649 (5,526, 5,779) | 48,706 (46,811, 50,691) | NA | NA | NA | NA | NA | NA | NA | NA |
| | | Cluster 2 | 2,438 (2,325, 2,561) | 13,787 (12,661, 15,063) | NA | NA | NA | NA | NA | NA | NA | NA |
| | | Cluster 1 | 646 (620, 675) | 1,929 (1,805, 2,058) | NA | NA | NA | NA | NA | NA | NA | NA |
| | | Pooled | 1,926 (1,785, 2,074) | 13,626 (11,990, 15,228) | NA | NA | NA | NA | 2,135 (2,041, 2,239) | 7,378 (6,845, 7,899) | NA | NA |
| | Patient 2 (Stage IVA, PFS ≥ 8 months) | Cluster 2 | 1,782 (1,710, 1,848) | 7,307 (6,910, 7,700) | NA | NA | NA | NA | NA | NA | NA | NA |
| | | Cluster 1 | 604 (573, 638) | 1,964 (1,800, 2,142) | NA | NA | NA | NA | NA | NA | NA | NA |
| | | Pooled | 1,576 (1,506, 1,642) | 6373 (5988, 6781) | 1,272 (1,225, 1,326) | 4,455 (4,199, 4,721) | NA | NA | NA | NA | NA | NA |
| Hepatocellular carcinoma | Patient 1 (Stage IV, PFS < 5 months) | Cluster 2 | 5,338 (5,252, 5,425) | 48,457 (47,098, 49,861) | NA | NA | NA | NA | NA | NA | NA | NA |
| | | Cluster 1 | 1,364 (1,325, 1,405) | 4,670 (4,402, 4,954) | NA | NA | NA | NA | NA | NA | NA | NA |
| | | Pooled | 3,660 (3,524, 3,800) | 29,969 (28,109, 31,625) | NA | NA | NA | NA | NA | NA | NA | NA |
| | Patient 2* (Stage IV, PFS ≥ 18 months) | Pooled | 1,871 (1,787, 1,949) | 7,961 (7,471, 8,460) | NA | NA | NA | NA | 1,760 (1,734, 1,787) | 4,150 (4,037, 4,259) | 1,947 (1,906, 1,986) | 5,255 (5,089, 5,415) |
| | Patient 3 (Stage I, PFS ≥ 18 months) | Cluster 2 | NA | NA | NA | NA | NA | NA | 4,149 (4,064, 4,241) | 16,410 (15,796, 17,043) | NA | NA |
| | | Cluster 1 | NA | NA | NA | NA | NA | NA | 2,368 (2,306, 2,429) | 7,131 (6,813, 7,462) | NA | NA |
| | | Pooled | 2,921 (2,876, 2,966) | 13,289 (12,897, 13,647) | NA | NA | NA | NA | 2,708 (2,634, 2,788) | 8,904 (8,447, 9,380) | 1,961 (1,918, 2,002) | 5,699 (5,491, 5,922) |
| Lung adenocarcinoma | Patient 1 {Stage IIIB} | Cluster 2 | 3,999 (3,921, 4,073) | 15,664 (15,180, 16,128) | NA | NA | NA | NA | NA | NA | 1,663 (1,612, 1,713) | 4,179 (3,995, 4,375) |
| | | Cluster 1 | 649 (616, 682) | 1,230 (1,132, 1,341) | NA | NA | NA | NA | NA | NA | 717 (675, 767) | 1,680 (1,498, 1,869) |
| | | Pooled | 1,869 (1,761, 1,979) | 6,489 (5,952, 7,050) | 3,233 (3,156, 3,314) | 9,846 (9,502, 10,176) | 724 (692, 754) | 1,479 (1,394, 1,569) | 1,371 (1,314, 1,432) | 3,200 (2,989, 3,417) | 1,520 (1,464, 1,574) | 3,801 (3,612, 3,993) |
| | Patient 2 {Stage IIB} | Cluster 2 | 2,778 (2,680, 2,871) | 8,458 (8,041, 8,868) | NA | NA | 2,097 (2,039, 2,160) | 6,123 (5,856, 6,411) | NA | NA | 1,898 (1,836, 1,968) | 5,179 (4,899, 5,486) |
| | | Cluster 1 | 831 (784, 880) | 1,703 (1,535, 1,897) | NA | NA | 586 (561, 612) | 1,091 (1,024, 1,168) | NA | NA | 649 (623, 676) | 1,255 (1,190, 1,321) |
| | | Pooled | 1612 (1515, 1703) | 4,411 (4,058, 4,763) | NA | NA | 1200 (1134, 1266) | 3,135 (2,917, 3,360) | 684 (652, 715) | 1,316 (1,232, 1,401) | 1,128 (1,071, 1,188) | 2,760 (2,549, 2,985) |
| Pancreatic adenocarcinoma | Patient 1 (Stage IV, PFS = 11 months) | Cluster 2 | 4,315 (4,205, 4,421) | 21,718 (20,860, 22,550) | 2,549 (2,437, 2,654) | 11,066 (10,341, 11,818) | NA | NA | NA | NA | NA | NA |
| | | Cluster 1 | 1,510 (1,439, 1,578) | 4,631 (4,314, 4,938) | 616 (588, 645) | 1,491 (1,393, 1,599) | NA | NA | NA | NA | NA | NA |
| | | Pooled | 3,323 (3,193, 3,458) | 15,675 (14,823, 16,549) | 1,423 (1,334, 1,527) | 5,489 (4,933, 6,075) | NA | NA | NA | NA | NA | NA |
| | Patient 2 (Stage IIB, PFS ≥ 26 months) | Cluster 2 | 2,235 (2,160, 2,306) | 8,896 (8,437, 9,317) | 1,382 (1,324, 1,442) | 6,017 (5,635, 6,386) | NA | NA | NA | NA | NA | NA |
| | | Cluster 1 | 997 (959, 1,039) | 3,381 (3,173, 3,581) | 779 (728, 831) | 2,496 (2,246, 2,765) | NA | NA | NA | NA | NA | NA |
| | | Pooled | 1,614 (1,542, 1,682) | 6,129 (5,715, 6,486) | 1,086 (1,028, 1,142) | 4,286 (3,965, 4,644) | NA | NA | NA | NA | NA | NA |

#### Supplementary Note Table 2. (Continued)

| Cancer type | Patient id | Cell cluster | Cell type | | | | | | | |
|---|---|---|---|---|---|---|---|---|---|---|
| | | | Myeloid | | T cell | | Natural killer cell | | B cell | |
| | | | Average gene counts (95% CI) | Average total UMI counts (95% CI) | Average gene counts (95% CI) | Average total UMI counts (95% CI) | Average gene counts (95% CI) | Average total UMI counts (95% CI) | Average gene counts (95% CI) | Average total UMI counts (95% CI) |
| Colorectal adenocarcinoma | Patient 1 (Stage IVA, PFS = 4 months) | Cluster 3 | NA | NA | NA | NA | NA | NA | NA | NA |
| | | Cluster 2 | 2,984 (2,920, 3,054) | 13,050 (12,518, 13,573) | NA | NA | NA | NA | NA | NA |
| | | Cluster 1 | 535 (523, 548) | 1,415 (1,372, 1,457) | NA | NA | NA | NA | NA | NA |
| | | Pooled | 1,787 (1,691, 1,877) | 7,365 (6,853, 7,920) | 1,211 (1,177, 1,243) | 3,600 (3,473, 3,743) | NA | NA | 1,102 (1,058, 1,149) | 5,422 (5,012, 5,873) |
| | Patient 2 (Stage IVA, PFS ≥ 8 months) | Cluster 2 | NA | NA | NA | NA | NA | NA | NA | NA |
| | | Cluster 1 | NA | NA | NA | NA | NA | NA | NA | NA |
| | | Pooled | 662 (627, 700) | 2,054 (1,881, 2,231) | 1,203 (1,177, 1,230) | 4,377 (4,250, 4,505) | NA | NA | 963 (911, 1,011) | 3,633 (3,330, 3,936) |
| Hepatocellular carcinoma | Patient 1 (Stage IV, PFS < 5 months) | Cluster 2 | NA | NA | NA | NA | NA | NA | NA | NA |
| | | Cluster 1 | NA | NA | NA | NA | NA | NA | NA | NA |
| | | Pooled | NA | NA | 1,318 (1,285, 1,350) | 3,720 (3,561, 3,879) | NA | NA | NA | NA |
| | Patient 2 (Stage IV, PFS ≥ 18 months) | Pooled | 1,406 (1,376, 1,434) | 4,228 (4,085, 4,368) | 1,303 (1,272, 1,340) | 3,221 (3,079, 3,366) | NA | NA | 1,105 (1,088, 1,122) | 11,686 (11,426, 11,965) |
| | Patient 3 (Stage I, PFS ≥ 18 months) | Cluster 2 | NA | NA | NA | NA | NA | NA | NA | NA |
| | | Cluster 1 | NA | NA | NA | NA | NA | NA | NA | NA |
| | | Pooled | 1,602 (1,571, 1,634) | 5,879 (5,711, 6,057) | 1,410 (1,373, 1,448) | 4,590 (4,412, 4,782) | NA | NA | NA | NA |
| Lung adenocarcinoma | Patient 1 {Stage IIIB} | Cluster 2 | 1,293 (1,253, 1,329) | 3,936 (3,776, 4,102) | NA | NA | NA | NA | NA | NA |
| | | Cluster 1 | 393 (381, 407) | 876 (838, 917) | NA | NA | NA | NA | NA | NA |
| | | Pooled | 1,233 (1,191, 1,271) | 3,732 (3,573, 3,889) | 584 (566, 602) | 1,050 (1,011, 1,090) | NA | NA | 544 (520, 568) | 2,569 (2,278, 2,858) |
| | Patient 2 {Stage IIB} | Cluster 2 | 1,207 (1,164, 1,251) | 3,504 (3,302, 3,721) | NA | NA | NA | NA | NA | NA |
| | | Cluster 1 | 361 (352, 370) | 728 (699, 754) | NA | NA | NA | NA | NA | NA |
| | | Pooled | 1,137 (1,089, 1,183) | 3,275 (3,080, 3,482) | 765 (745, 789) | 1362 (1310, 1417) | NA | NA | 762 (732, 796) | 4,396 (4,083, 4,746) |
| Pancreatic adenocarcinoma | Patient 1 (Stage IV, PFS = 11 months) | Cluster 2 | 3,213 (3,131, 3,292) | 16,221 (15,618, 16,851) | NA | NA | NA | NA | NA | NA |
| | | Cluster 1 | 1,460 (1,420, 1,504) | 3,840 (3,688, 3,980) | NA | NA | NA | NA | NA | NA |
| | | Pooled | 1,788 (1,726, 1,851) | 6,158 (5,746, 6,568) | 1,531 (1,508, 1,554) | 4,814 (4,716, 4,922) | 1,651 (1,630, 1,671) | 4,064 (4,001, 4,126) | 1,352 (1,327, 1,378) | 4,119 (4,022, 4,213) |
| | Patient 2 (Stage IIB, PFS ≥ 26 months) | Cluster 2 | 2,210 (2,155, 2,271) | 10,285 (9,860, 10,748) | NA | NA | NA | NA | NA | NA |
| | | Cluster 1 | 890 (857, 926) | 2,523 (2,344, 2,711) | NA | NA | NA | NA | NA | NA |
| | | Pooled | 1,960 (1,891, 2,030) | 8,818 (8,327, 9,245) | 936 (919, 953) | 2,526 (2,467, 2,586) | 1,169 (1,148, 1,191) | 2,855 (2,763, 2,950) | 927 (904, 950) | 2,684 (2,586, 2,775) |

PFS: progression free survival.
*: for a patient, if all cell types have one cluster each, only the results from the pooled cells of each cell type are shown.
NA: due to no cells or only one cell cluster in the corresponding cell type; for the latter, the results of gene counts and total UMI counts are shown in the "Pooled" position.

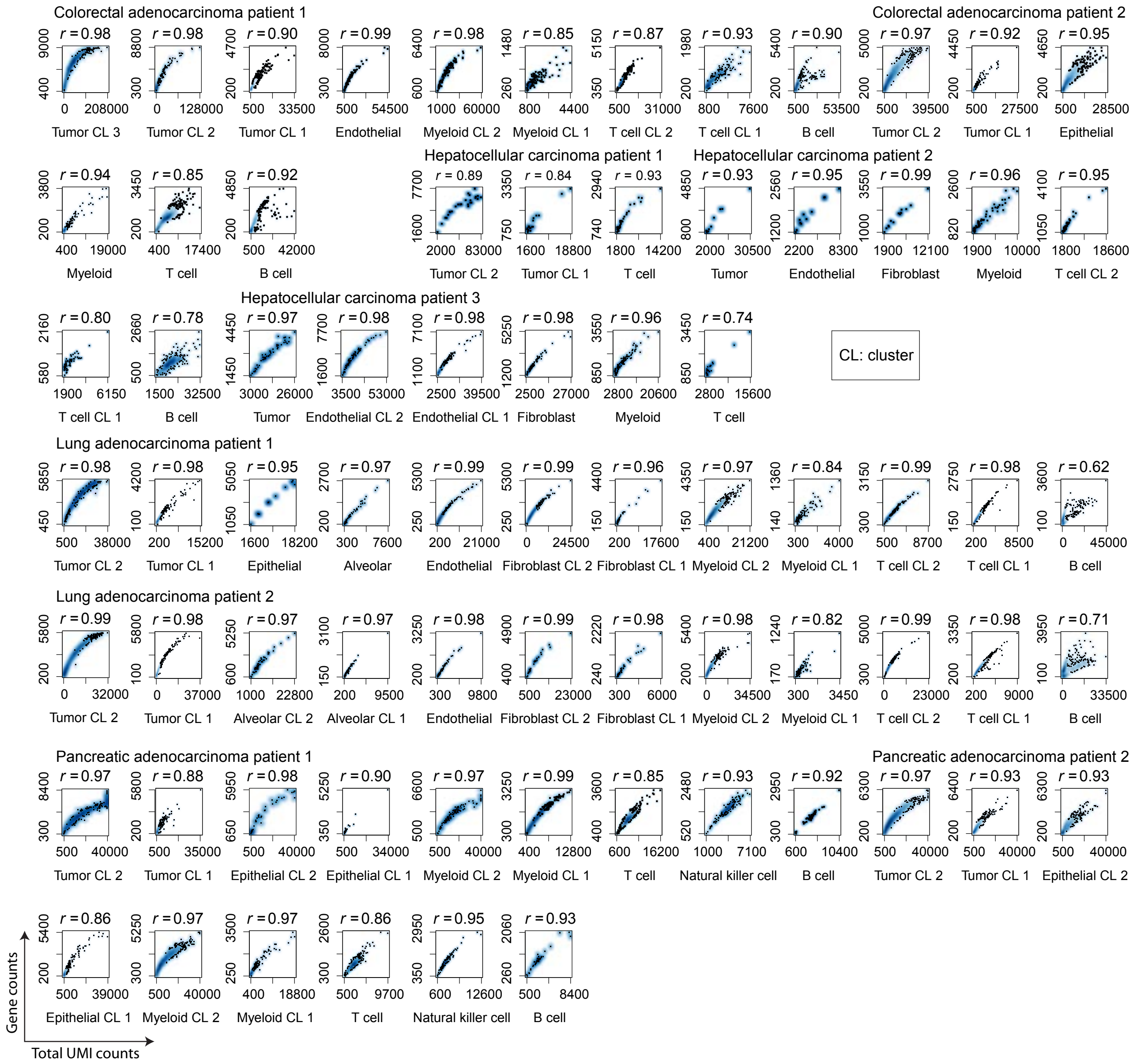

**Supplementary Note Figure 3. Correlations between gene counts and total UMI counts.** Smoothed scatter plots show the correlations between gene counts and total UMI counts in cell clusters from each patient sample. In each smoothed scatter plot, the Spearman correlation coefficient is labeled on the top (r).

**Supplementary Note Table 3. T-tests of total UMI counts between two tumor cell clusters within each patient across four cancer types.**

| Cancer type | Patient 1 | | Patient 2 | |
|---|---|---|---|---|
| | *P* value | $\mu_2/\mu_1$(95% CI)* | *P* value | $\mu_2/\mu_1$(95% CI) |
| Colorectal adenocarcinoma | $1 \times 10^{-11}$ | 25 (23, 27) | $6 \times 10^{-10}$ | 3.7 (3.4, 4.2) |
| Hepatocellular carcinoma | $7 \times 10^{-7}$ | 10 (10,11) | NA | NA |
| Lung adenocarcinoma | $< 2 \times 10^{-16}$ | 13 (12, 14) | $< 2 \times 10^{-16}$ | 5.0 (4.4, 5.6) |
| Pancreatic adenocarcinoma | $2 \times 10^{-15}$ | 4.7 (4.3, 5.0) | 0.009 | 2.6 (2.4, 2.8) |

* $\mu_2$ and $\mu_1$ are the means of the total UMI counts from the tumor cell high-UMI cluster and tumor cell low-UMI cluster, respectively.

#### 2. TUMOR-SPECIFIC TOTAL MRNA EXPRESSION IN BULK SEQUENCING DATA

##### 2.1. A mathematical model for tumor-specific total mRNA expression

For any group of cells, we use $S$ to denote the average global mRNA transcript level per cell per haploid genome, which follows $S = \sum_{c=1}^{C} (\sum_{g=1}^{G} u_{gc}/p_c)/C$. Here $p_c$ is the ploidy, *i.e.*, the number of copies of the haploid genome in cell *c*. However, the cell level ploidy $p_c$ is usually not measurable. Hence, in practice, we use average ploidy $\Psi$ of the corresponding cell group to approximate it: $S \approx \sum_{c=1}^{C} \sum_{g=1}^{G} u_{gc}/(C\Psi)$. For non-tumor cells, which are commonly diploid, this assumption is assured.

###### 2.1.1. Model

In the analysis of bulk RNAseq data from mixed tumor samples, we are interested in comparing tumor with non-tumor cell groups. We denote tumor cells by group *T*, and non-tumor cells by group *N*. Therefore, we define a tumor-specific total mRNA expression score (TmS) to reflect the ratio of total mRNA transcript level per haploid genome of tumor cells to that of the surrounding non-tumor cells, *i.e.*, $TmS_{tumor} = S_T / S_N$, simplified as TmS from here forward. It is necessary to calculate this ratio in order to cancel out technical effects presented in sequencing data that confound with both $S_T$ and $S_N$. Let $T_g = \sum_{c=1}^{C_T} u_{gc}$ and $N_g = \sum_{c=1}^{C_N} u_{gc}$ denote the total number of mRNA transcripts of gene g across all cells from tumor and non-tumor cells, let $C_T$ and $C_N$ denote the total number of tumor and non-tumor cells, and let $\Psi_T$ and $\Psi_N$ represent the average ploidy of tumor and non-tumor cells, respectively. Under the assumption that the tumor cells have a similar ploidy, we can derive TmS without using single-cell-specific parameters as

$$TmS = \frac{\sum_{g=1}^{G} T_g/(C_T\Psi_T)}{\sum_{g=1}^{G} N_g/(C_N\Psi_N)}. \quad \text{Eq.S1}$$

Here we further introduce a tumor-specific mRNA proportion $\pi = (\sum_{g=1}^{G} T_g) / (\sum_{g=1}^{G} T_g + \sum_{g=1}^{G} N_g)$ and a tumor cell proportion of tumor cells within a sample (termed "tumor purity") $\rho = C_T / (C_T + C_N)$. Note that deconvolution of just the gene expression data will not provide information on the total number of tumor cells and non-tumor cells, but only the sum of total mRNA expression across all cells of each cell type.

Using these parameters, we rewrite Eq.S1 as

$$TmS = \frac{\pi}{1-\pi} / \frac{\rho}{1-\rho} / \frac{\Psi_T}{\Psi_N}. \qquad \text{Eq.S2}$$

Additionally, we can define a ploidy-unadjusted *TmS* by removing the ploidy terms.

###### 2.1.2. Estimation

It is common practice to assume the ploidy of non-tumor cells $\Psi_N$ equals to 2[15,16]. Hence, we have

$$\widehat{TmS} = \frac{\hat{\pi}}{1-\hat{\pi}} / \frac{\hat{\rho}}{1-\hat{\rho}} / \frac{\widehat{\Psi_T}}{2}. \qquad \text{Eq.S3}$$

In what follows, we use *TmS* to represent $\widehat{TmS}$, for the sake of simplicity.

Estimation of tumor-specific mRNA proportion $\pi$ using high-throughput RNA sequencing has not been possible due to several technical and analytical factors including: 1) the need to account for technical artifacts introduced by varied library size, which currently involves normalization procedures across samples; 2) total mRNA transcripts per cell are confounded with technical artefacts so that normalization procedures adjust for both effects at once, consequently losing the ability to evaluate the downstream global transcriptome feature[17]; and 3) a limited focus on estimating cell proportions by popular methods[18–20].

Using deconvolution to partition tumor and non-tumor cells within the same sample under the same experimental conditions provides a mathematical means to cancel out the effect of technical artefacts while maintaining the effect of cell-type-specific total mRNA counts. We use the DeMixT model[21] to estimate $\pi$. For sample *i* and across any gene *g*, we have

$$Y_{ig} = \pi_i T'_{ig} + (1-\pi_i) N'_{ig} \qquad \text{Eq.S4}$$

where $Y_{ig}$ represents the scale normalized expression matrix from mixed tumor samples, $T'_{ig}$ and $N'_{ig}$ represent the normalized relative expression of gene *g* within tumor and surrounding non-tumor cells, respectively. The estimated $\hat{\pi}$ is the quantity desired for Eq.S3.

Computational deconvolution methods, e.g., ASCAT[16] and ABSOLUTE[15], have been developed to perform allele-specific copy number analysis and to estimate tumor purity $\rho$ and ploidy $\Psi_T$ from tumor DNA sequencing data. Such statistical methods jointly model the distribution of *logR* and *B* allele (or variant allele) frequency (BAF) across germline SNPs, with tumor purity and allele-specific copy number as parameters of interest. Then the tumor purity and ploidy (the average tumor copy number) can be estimated through minimizing the loss function or maximizing the likelihood. Below, we provide a detailed description for these methods using the ASCAT model as an example.

Sequence read counts at known SNP loci were computed from tumor DNA sequencing data. The $logR_i$ can be computed from the total read counts in the tumor versus normal for the $i$th SNP, which provides information on the ratio of total copy number between the tumor and the normal. Specifically, $logR_i$ can be expressed as[16]

$$logR_i = \gamma log_2\left(\frac{2(1\text{-}\rho)+\rho(n_{A,i}+n_{B,i})}{2(1\text{-}\rho)+\psi_T}\right), \qquad \text{Eq.S5}$$

where $\rho$ is the tumor purity, $\psi_T$ is the tumor ploidy, $\gamma$ is a constant depends on which DNA sequencing technology is used. $n_{A,i}$ and $n_{B,i}$ stand for the allele-specific copy number of A allele and B allele for the $i$th SNP in tumor cells, respectively.

On the other hand, allelic imbalance can be inferred from the $BAF_i$ for $i$th SNP. The $BAF_i$ can be expressed as[16]

$$BAF_i = \frac{1\text{-}\rho+\rho n_{B,i}}{2(1\text{-}\rho)+\rho(n_{A,i}+n_{B,i})}. \qquad \text{Eq.S6}$$

Based on Eq.S5 and Eq.S6, the allele-specific copy number can be expressed as a function of the tumor purity and ploidy. Specifically, we have

$$\hat{n}_{A,i} = \frac{\rho\text{-}1+2^{\frac{logR_i}{\gamma}}(1-BAF_i)(2(1\text{-}\rho)+\Psi_T)}{\rho};$$

$$\hat{n}_{B,i} = \frac{\rho\text{-}1+2^{\frac{logR_i}{\gamma}}BAF_i(2(1\text{-}\rho)+\Psi_T)}{\rho}.$$

Allele-specific piecewise constant fitting (ASPCF)[16] was then applied to both $logR_i$ and $BAF_i$ simultaneously, which enforced the change points to occur at the same genomic locations. Consequently, a segmentation of the genome was obtained, each segment corresponding to a genomic region between two adjacent change points. Using the ASPCF smoothed $logR_i$ and $BAF_i$, the final values for $\hat{\rho}$ and $\hat{\Psi}_T$ were obtained through the optimization, such that the allele-specific copy number estimates $\hat{n}_{A,i}$ and $\hat{n}_{B,i}$ were as close to nonnegative integers as possible for germline heterozygous SNPs.

##### 2.2. Improved estimation using DeMixT

Many computational deconvolution methods have been developed to estimate cell type proportions through transcriptome data; however, most of them focus on the cellular proportion and not the global gene expression level of each cell type, due to lack of appropriate normalization approaches. The DeMixT[21] model is unique in aiming to estimate the global tumor-specific gene expression level relative to the normal reference in the context of admixed tumor samples. ISOpure[22] is the other model that presents similar objectives as the DeMixT model. The following issues and our proposed solution are generally applicable to both models.

The identifiability of model parameters is a major issue for high dimensional models. Due to technical limitations, given a certain amount and quality of experimental data, not all model parameters are guaranteed for unambiguous estimation. Frequently, only a subset of model parameters are identifiable based on the available data, with the rest of the parameters considered unidentifiable. Confidence intervals can be derived for identifiable parameters, which contain the true value of the parameter with a desired probability[23]. Fortunately, with the DeMixT model, there is hierarchy in model identifiability in which the cell-type specific global gene expression proportions ($\pi$) are the most identifiable parameters, requiring only a subset of genes with identifiable expression distributions. Therefore, our goal is to select an appropriate set of genes as input to DeMixT that optimizes the estimation of $\pi$. In general, genes are expressed at different levels, which, due to different numerical ranges, can affect estimation of $\pi$. We found that including genes that are not differentially expressed between the tumor and non-tumor components within the bulk sample, or genes with large variance in expression within the non-tumor component, can introduce large biases into the estimated $\pi$. On the other hand, the tumor component is hidden in the mixed tumor samples, hence preventing a DE analysis between mixed and normal samples from finding the best genes. By applying a profile-likelihood based approach to detect the identifiability of model parameters[24], we systematically evaluated the identifiability for all available genes based on the data, and selected the most identifiable genes for the estimation of $\pi$. As a result, the accuracy of the estimated proportions has been improved. As a general method, the profile-likelihood based gene selection strategy can be extended to any method that uses maximum likelihood estimation. We also employed an additional virtual spike-in strategy to balance proportion distributions which further improved model identifiability.

###### 2.2.1. Likelihood model for DeMixT

In the DeMixT model[21] (Eq.S4), we assumed that the observed expression level $Y_{ig}$ is a linear combination of two hidden components $T_{ig}$ (tumor, in place of $T'_{ig}$ from now on) and $N_{ig}$ (non-tumor, in place of $N'_{ig}$ from now on), where gene $g$ = 1,2,…,$G$, sample $i$ = 1,2,…,$M$ , and $\pi_i$ is the tumor-specific total mRNA expression proportions. We assume each hidden component follows the $log_2$-normal distribution, *i.e.*, $T_{ig} \sim LN\left(\mu_{Tg}, \sigma^2_{Tg}\right)$ and $N_{ig} \sim LN\left(\mu_{Ng}, \sigma^2_{Ng}\right)$.

Fitting the deconvolution model in Eq.S1 can be formally defined as an optimization problem that seeks to identify optimal estimates for sample-level, tumor-specific mRNA proportions $\pi_i$, and gene-level parameters. The full parameter set is denoted by $(\boldsymbol{\pi},\boldsymbol{\mu_T},\boldsymbol{\sigma_T})$, where $\boldsymbol{\pi} = (\pi_1,\pi_2,\ldots,\pi_M)$, $\boldsymbol{\mu_T} = (\mu_{T1}, \mu_{T2},\ldots, \mu_{TG})$, $\boldsymbol{\sigma_T} = (\sigma_{T1}, \sigma_{T2},\ldots,\sigma_{TG})$. The full log-likelihood of the DeMixT model can be written as

$$l(\boldsymbol{\pi},\boldsymbol{\mu_T},\boldsymbol{\sigma_T})=\sum_{i=1}^{M}\sum_{g=1}^{G} log(f(Y_{ig}|\pi_i,\mu_{Tg},\sigma_{Tg})),$$

where $f\left(Y_{ig}|\pi_i,\mu_{Tg},\sigma_{Tg}\right)=\frac{1}{2\pi\sigma_{Ng}\sigma_{Tg}}\int_0^{Y_{ig}}\frac{1}{t(Y_{ig}-t)}exp(-\frac{(log2(t)-\mu_{Ng}-log2(1-\pi_i))^2}{2\sigma^2_{Ng}}-\frac{(log2(Y_{ig}-t)-\mu_{Tg}-log2(\pi_i))^2}{2\sigma^2_{Tg}})dt.$

The DeMixT model applies an optimization method, iterated conditional modes (ICM)[25], to maximize the full log-likelihood function and estimate all distribution parameters $(\boldsymbol{\mu_T},\boldsymbol{\sigma_T})$ and proportions $\boldsymbol{\pi}$.

##### 2.2.2. Optimized model identifiability

Based on the most stringent definition, for a parametric model $l(\boldsymbol{Y} \mid \theta)$, $\theta$ is identifiable if, $l(\boldsymbol{Y} \mid \theta_1) = l(\boldsymbol{Y} \mid \theta_2) => \theta_1 = \theta_2$. However, this rigorous identifiability is difficult to validate for a general high-dimensional and non-convex model, which is the case for the DeMixT model. Thus, for a parameter $\theta$, we use the confidence interval $[\theta^-, \theta^+]$ to measure its identifiability[24].

In the DeMixT model, if we select genes with small confidence intervals of $\mu_{Tg}$ based on profile likelihood, which indicate high identifiability, the corresponding gene *g* will be more stable and reliable, so will the inferred tumor-specific mRNA proportions ($\boldsymbol{\pi}$). As a result, the length of confidence interval of $\mu_{Tg}$ serves as an estimable quantity with which we can evaluate the gene *g*'s identifiability and prioritize genes to increase the estimation quality of $\pi_i$, $\mu_{Tg}$, $\sigma_{Tg}$.

The profile likelihood is preferred to compute confidence intervals of parameters that often have better small-sample properties than those based on asymptotic standard errors calculated from the full likelihood[26]. Assume the *k*th gene's mean parameter $\mu_{Tk}$ is the parameter of interest. The definition of the profile likelihood function of $\mu_{Tk}$ is:

$$l_{\mu_{Tk}}(\mu_{Tk}=x|\boldsymbol{\pi},\boldsymbol{\mu_T},\boldsymbol{\sigma_T}) = max\{\sum_{i=1}^{M}[\sum_{g\neq k}^{G} log\left(f\left(\pi_i,\mu_{Tg},\sigma_{Tg}\right)\right) +log\left(f\left(\pi_i,\mu_{Tk}=x,\sigma_{Tk}\right)\right)]\}$$

The confidence interval of a profile likelihood function can be constructed through inverting a likelihood-ratio test [27]. Assume the null hypothesis as $H_0$: $\mu_{Tk} = x$, and the maximum likelihood estimator of ($\pi_i$, $\mu_{Tg}$,

$\sigma_{Tg}$) are ($\widehat{\pi}_i$, $\widehat{\mu}_{Tg}$, $\widehat{\sigma}_{Tg}$). The null hypothesis will not be rejected at the $\alpha$ level of significance if and only if $2\left[l(\widehat{\boldsymbol{\pi}}, \widehat{\boldsymbol{\mu}}_{\boldsymbol{T}}, \widehat{\boldsymbol{\sigma}}_{\boldsymbol{T}}) - l_{\mu_{Tk}}(\mu_{Tk} = x \mid \widehat{\boldsymbol{\pi}}, \widehat{\boldsymbol{\mu}}_{\boldsymbol{T}}, \widehat{\boldsymbol{\sigma}}_{\boldsymbol{T}})\right] \leq \chi^2_{1-\alpha}(1)$, where $\chi^2_{1-\alpha}(1)$ stands for $1$-$\alpha$ percentile of the $\chi^2$ distribution with degrees of freedom equal to $1$. Since the maximized likelihood $l(\widehat{\boldsymbol{\pi}}, \widehat{\boldsymbol{\mu}}_{\boldsymbol{T}}, \widehat{\boldsymbol{\sigma}}_{\boldsymbol{T}})$ and model parameters $\widehat{\boldsymbol{\pi}}, \widehat{\boldsymbol{\mu}}_{\boldsymbol{T}}, \widehat{\boldsymbol{\sigma}}_{\boldsymbol{T}}$ can be estimated by running the DeMixT model on all available gene sets, for any given $x$, we are able to investigate the profile log-likelihood function $l_{\mu_{Tk}}(\mu_{Tk} = x \mid \widehat{\boldsymbol{\pi}},\widehat{\boldsymbol{\mu}}_{\boldsymbol{T}},\widehat{\boldsymbol{\sigma}}_{\boldsymbol{T}})$. Consequently, we can estimate the lower and upper bounds of the confidence interval [$\mu^-_{Tk}$, $\mu^+_{Tk}$] as

$$\mu^-_{Tk} = \min_x \{x \mid 2\left[l(\widehat{\boldsymbol{\pi}},\widehat{\boldsymbol{\mu}}_{\boldsymbol{T}},\widehat{\boldsymbol{\sigma}}_{\boldsymbol{T}}) - l_{\mu_{Tk}}(\mu_{Tk} = x \mid \widehat{\boldsymbol{\pi}},\widehat{\boldsymbol{\mu}}_{\boldsymbol{T}},\widehat{\boldsymbol{\sigma}}_{\boldsymbol{T}})\right] \leq \chi^2_{1-\alpha}(1)\}$$
$$\mu^+_{Tk} = \max_x \{x \mid 2\left[l(\widehat{\boldsymbol{\pi}},\widehat{\boldsymbol{\mu}}_{\boldsymbol{T}},\widehat{\boldsymbol{\sigma}}_{\boldsymbol{T}}) - l_{\mu_{Tk}}(\mu_{Tk} = x \mid \widehat{\boldsymbol{\pi}},\widehat{\boldsymbol{\mu}}_{\boldsymbol{T}},\widehat{\boldsymbol{\sigma}}_{\boldsymbol{T}})\right] \leq \chi^2_{1-\alpha}(1)\}$$

Following the same procedure, we can derive the confidence interval of $\mu_{Tk}$ for all available genes.

In real data analysis, calculating the actual profile likelihood function of $\mu_{Tk}$ across all 20,000 genes is generally infeasible due to computational limits. An asymptotic approximation is necessary in order to quickly evaluate the profile likelihood function. If the measurement noise is small and the sample size is large enough, asymptotic confidence intervals are good approximations of the actual confidence intervals[24]. The asymptotic profile likelihood function can be derived from the observed Fisher information of the log likelihood, denoted as $H(\widehat{\boldsymbol{\pi}},\widehat{\boldsymbol{\mu}}_{\boldsymbol{T}},\widehat{\boldsymbol{\sigma}}_{\boldsymbol{T}})$. Then the asymptotic $\alpha$ level confidence interval of $\mu_{Tk}$ can be written as follows[24]

$$\mu_{Tk}{}^{\pm} = \widehat{\mu_{Tk}} \pm \sqrt{2\chi^2_{1-\alpha}(1)\, H(\widehat{\boldsymbol{\pi}},\widehat{\boldsymbol{\mu}}_{\boldsymbol{T}},\widehat{\boldsymbol{\sigma}}_{\boldsymbol{T}})^{-1}_{k,k}}\,. \qquad \text{Eq.S7}$$

*Validation.* We compared the actual profile likelihood function with the asymptotic profile likelihood function for a random set of 20 genes in real data (the TCGA prostate adenocarcinoma dataset) and observed good performance of the approximate profile likelihoods (**Supplementary Note Figure 4**). With 20 randomly selected genes, we calculated the root mean squared error (RMSE) between the confidence intervals from the true and asymptotic profile likelihoods is 0.05.

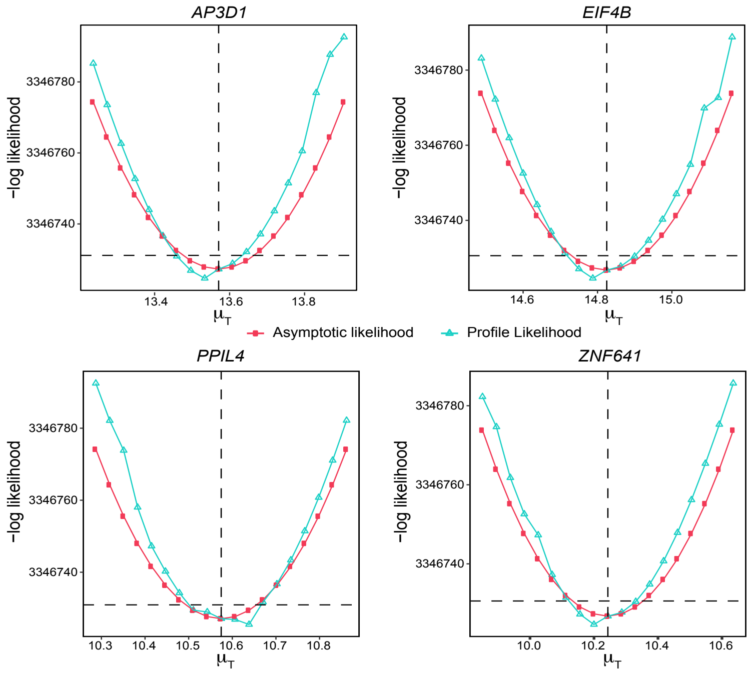

**Supplementary Note Figure 4. Asymptotic profile likelihoods for 4 genes using 259 samples from the TCGA prostate cancer dataset.** Comparison of asymptotic and actual profile likelihoods of $\mu_T$ for 4 randomly selected genes in the TCGA prostate adenocarcinoma data. The red curve shows the true profile likelihood of the corresponding parameter. The blue curve shows an asymptotic approximation of the profile likelihood of the corresponding parameter.

*Gene selection score*. We now introduce a metric, the gene selection score, which for gene $k$ is the width of the asymptotic 95% profile likelihood-based confidence interval of $\mu_{Tk}$ for gene $k$

$$gene\ selection\ score_k = 2\sqrt{2\chi^2_{0.05}(1)\ H(\hat{\pi},\hat{\mu}_T,\hat{\sigma}_T)^{-1}_{k,k}}. \qquad \text{Eq.S8}$$

Genes with a lower score have a smaller confidence interval, hence higher identifiability in their corresponding parameters. Genes are ranked based on the gene selection score from the smallest to the largest. A subset of genes that are ranked on top will be used for parameter estimation. In the DeMixT R package (freely available from Bioconductor), our proposed profile-likelihood based gene selection approach is included as function “DeMixT_GS”.

##### 2.2.3. A simulation study for profile-likelihood based gene selection

The DeMixT model assumes every gene $g$ has a shared mean ($\mu_{Tg}$) and variance ($\sigma_{Tg}$) parameters across all tumor samples. However, in real data, this assumption will be violated in selected genes, due to the fact that some genes are significantly differentially expressed in different subtypes of the cancer. For example, the PAM50 genes are known to be differentially expressed in different molecular subtypes in breast cancer, e.g., Basal, Her2, LumA, and LumB subtypes. Therefore, our simulation aimed to assess the performance of the proposed gene selection method in finding genes that best follow the DeMixT model. The detailed simulation design is described below.

We simulated gene expression of 269 mixed samples and 100 normal references with 10,000 genes, mimicking the real data scenario presented in the TCGA prostate adenocarcinoma dataset. The true $\pi$ were set as the tumor cell proportions derived from ASCAT. We generated the expression of 10,000 genes under four scenarios: 1) genes that are consistently differentially expressed between the tumor and normal components; 2) genes that are differentially expressed across subtypes of tumor samples; 3) genes that are consistently expressed similarly between the tumor and normal components; 4) genes that are expressed with large variances. Specifically, for scenario 1), we generated 6,000 genes for the pure tumor $T_{ig}$ and normal references $N_{ig}$ with distributions $log_2(T_{ig}) \sim N(\mu_{Tg}, \sigma_{Tg}^2)$ and $log_2(N_{ig}) \sim N(\mu_{Ng}, \sigma_{Ng}^2)$, where $i$ denotes sample, $i=1, \cdots, M$. We simulated $\mu_{Ng}, \mu_{Tg} \sim N(7,1.5^2)$, and $\sigma_{Ng}$, $\sigma_{Tg}$ were sampled with replacement from the observed standard deviations from the normal samples of TCGA prostate adenocarcinoma. For scenario 2), we generated additional 2,000 subtype genes with samples spit into equal-sized subgroups $M_1$, $M_2$, and $M_3$ with corresponding $\mu_{T_1g} \sim N(5,1.5^2)$, $\mu_{T_2g} \sim N(7,1.5^2)$, $\mu_{T_3g} \sim N(9,1.5^2)$ and $M= M_1+M_2+ M_3$. Then we generate the expression with $log_2(N_{ig}) \sim N(\mu_{Ng}, \sigma_{Ng}^2)$, $log_2(T_{ig}) \sim N(\mu_{T_kg}, \sigma_{Tg}^2)$, $k=1, 2, 3, i \in M_k$. For scenario 3), we generated 1,000 genes with strictly equal mean expression for pure tumor and normal reference, where $\mu_{Ng}=\mu_{Tg} \sim N(7,1.5^2)$, $log_2(T_{ig}) \sim N(\mu_{Tg}, \sigma_{Tg}^2)$ and $log_2(N_{ig}) \sim N(\mu_{Ng}, \sigma_{Ng}^2)$. For scenario 4), we generated the remaining 1,000 genes with large variances, where $\mu_{Ng}, \mu_{Tg} \sim N(7,1.5^2)$, $\sigma_{Ng} = \sigma_{Tg} = 1.5$, and the expression profile follow $log_2(T_{ig}) \sim N(\mu_{Tg}, \sigma_{Tg}^2)$, $log_2(N_{ig}) \sim N(\mu_{Ng}, \sigma_{Ng}^2)$. Then we mixed the $T_{ig}$ and $N_{ig}$ component expression linearly at the generated $\pi$ according to the DeMixT model: $Y_{ig}= \pi_i T_{ig}+(1-\pi_i)N_{ig}$, where $G=10,000$, $M=269$. Our proposed gene selection method ("DeMixT_GS") successfully ranked genes from scenario 1) much higher than others, whereas a routine DE analysis using the two-sided t-test statistic between mixed tumor and normal samples failed to identify these genes (**Supplementary Fig. 4b**). Across simulations where we selected 100, 250, 1500, 2500, and 8000 genes, "DeMixT_GS" always outperformed "DeMixT_DE" in estimating proportions (**Supplementary Note Figure 5a**). The dip test[28] was used to measure the unimodality of the distribution of gene expression. This test statistic was designed to test multimodality of a random variable based on the maximum difference between the empirical distribution and the unimodal distribution of all observed data points (**Supplementary Note Figure 5b**). It suggests that the proposed gene selection method successfully ranked subtype-specific genes lower than the DE method.

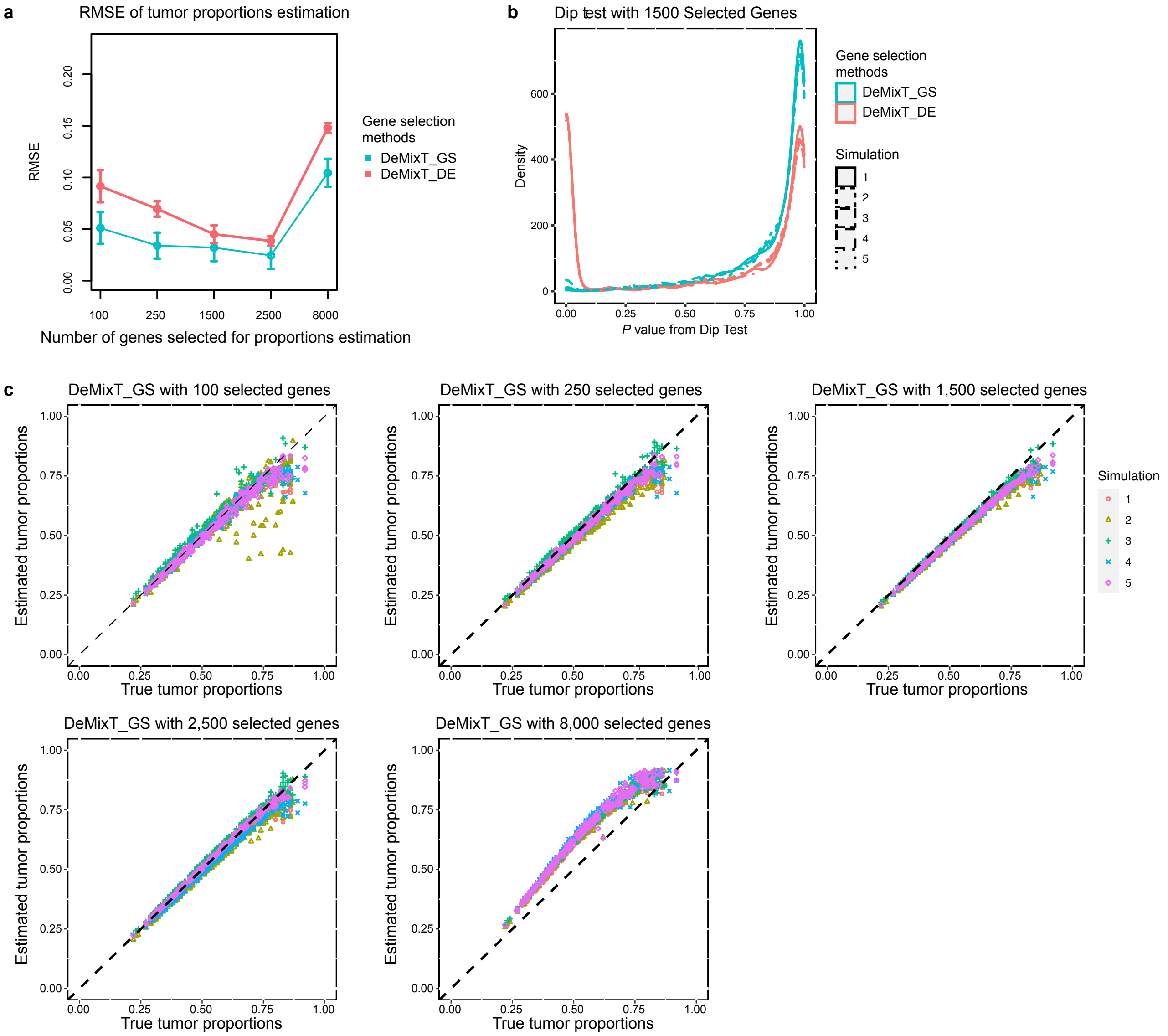

**Supplementary Note Figure 5. Profile-likelihood based gene selection (DeMixT_GS) improves tumor-specific total mRNA expression proportions estimation. a**, Root Mean Square Error (RMSE) was calculated for each simulation scenario. **b**, Density of *P* values based on a dip test for selected genes by "DeMixT_GS" and "DeMixT_DE" methods. The dip test was applied to indicate the distribution of gene expression for selected genes based on the "DeMixT_GS" and "DeMixT_DE" methods, respectively. A small *P* value of the dip test suggests the corresponding gene is not unimodally distributed, which violates the model assumption of $\log_2$-normal distribution across samples. **c**, Scatter plot of true versus estimated tumor-specific total mRNA expression proportions using "DeMixT_GS" method with different numbers of top-ranking genes with the smallest gene selection score.

*Optimal selection of genes*. We also observed that the number of genes selected by "DeMixT_GS" influences the performance of DeMixT. The accuracies of $\pi$ estimation based on 100, 250, 1,500, 2,500 and 8,000 genes selected by the proposed "DeMixT_GS" were compared. **(Supplementary Note Figure**

**5a,c)**. Accurate $\pi$ estimation, as measured by the RMSE, was achieved with 1,500 or 2,500 genes. In real data, we used either the top 1,500 or top 2,500 genes to estimate $\pi$.

##### 2.2.4. Virtual spike-ins to improve identifiability and a simulation study

When the true proportions are skewed towards the high end (*i.e.*, median above 0.5), which is expected to occur frequently in real data (tumor samples with a low percentage of tumor cells are already discarded), the DeMixT estimation procedure, after careful gene selection, tends to slightly underestimate the high proportions. We hypothesize that by shifting the mode of the $\pi$ distribution close to 0.5, the issue of underestimation will be alleviated. To achieve this, we simulate additional "mixed tumor" samples, *i.e.* spike-ins, with close to 0% of $\pi$, so that there are roughly the same number of samples with tumor proportions below and above 50%, *i.e.*, $S_P + \left|\{i \mid \rho_i<0.5\}\right| \cong |\{i \mid \rho_i \geq 0.5\}|$, where $S_P$ represents the number of spike-ins, $\rho_i$ represent tumor purity of sample $i$, and the $|\cdot|$ represent cardinality of a set. For the cancer type whose median tumor purity is below 0.5, we set $S_P$ at 5. The spike-ins are generated based on gene expression profiles observed from the input data of normal reference samples.

We simulated 100 mixed samples and 100 normal reference samples with 8,000 genes based on simulation settings described in **Section 2.2.3** with five replicates. Tumor-specific mRNA proportions were simulated from a normal distribution (mean = 0.55, SD = 0.2) and truncated at endpoints of 0.05 and 0.95. $\mu_{Ng}, \mu_{Tg} \sim N(7, 1.5^2)$ and $\sigma_{Ng}, \sigma_{Tg} \sim U(0.1, 0.8)$. The expression level of spike-ins is denoted as $P_{jg}$. We simulate $P_{jg} \sim LN\left(\hat{\mu}_{Ng}, \hat{\sigma}^2_{Ng}\right)$, for gene $g = 1,2,\ldots,G$ and sample $j = 1,2,\ldots,S_P$. The spike-ins were then combined with mixed tumor samples. We ran DeMixT on the combined samples while fixing $\pi$ for the spike-ins at 0.01. We found adding spike-ins can reduce biases in the estimation of tumor-specific

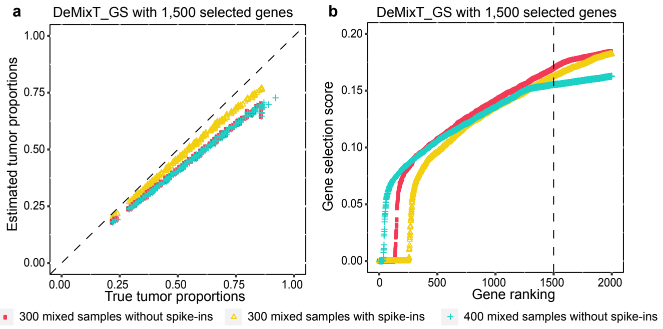

**Supplementary Note Figure 6. Adding spike-ins reduces systematic bias of estimated tumor-specific mRNA proportions at a high end. a**, Scatter plot of true versus estimated tumor-specific mRNA proportions using "DeMixT_GS" method under different strategies of adding spike-ins. **b**, Distribution of gene selection score to top 2,000 genes under different strategies of adding spike-ins, where the x-axis represents the rank of genes sorted by gene selection score from low to high and the y-axis represents the estimated gene selection score for the corresponding genes.

mRNA proportions (**Supplementary Note Figure 6a**), as demonstrated by the improved gene selection scores (the smaller the better) for the top-ranking genes (**Supplementary Note Figure 6b**).

##### 2.3. Tumor-specific total mRNA expression in patient samples

###### 2.3.1. Datasets

*The Cancer Genome Atlas data*

Publicly available transcriptome profiling HT-seq raw read counts from 7,054 tumor samples from 15 cancer types in TCGA (breast adenocarcinoma, bladder urothelial carcinoma, colorectal cancer (colon adenocarcinoma + rectum adenocarcinoma), head-and-neck squamous cell carcinoma, kidney chromophobe, kidney renal clear cell carcinoma, kidney renal papillary cell carcinoma, liver hepatocellular carcinoma, lung adenocarcinoma, lung squamous cell carcinoma, pancreatic adenocarcinoma, prostate adenocarcinoma, stomach adenocarcinoma, thyroid carcinoma, uterine corpus endometrial carcinoma) were downloaded from the GDC data portal (v14.0)[29] (https://portal.gdc.cancer.gov/). They were generated through the standard RNAseq analysis pipeline (https://docs.gdc.cancer.gov/Data/Bioinformatics_Pipelines/Expression_mRNA_Pipeline/) by aligning reads to the GRCh38 reference genome and then by quantifying the mapped reads. Cinical annotation data including overall survival (OS), progression-free interval (PFI), pathologic stage, age, and sex of patients across 15 cancer types was downloaded from the GDC data portal (https://gdc.cancer.gov/about-data/publications/pancanatlas). Somatic mutation data of the 15 cancer types were downloaded from the re-annotated mutation annotation file (MAF) format at the GDC (https://gdc.cancer.gov/about-data/publications/mc3-2017). ABSOLUTE tumor purity and ploidy data were downloaded from Aran *et al*[30]. ASCAT tumor purity and ploidy data were downloaded from Alexandrov *et al*[31]. Driver mutation and indels annotation were downloaded from the TCGA pan-cancer driver mutation database: http://intogen.org/download version 2016.5[32]. NarrowPeak format ATAC-seq data for TCGA samples was obtained from Corces *et al*[33]. NarrowPeak files were annotated using the R package chipseeker[34]. Peaks outside of promoter regions (-2kb to 1kb of transcription start sites) were excluded. For breast adenocarcinoma, molecular subtype, triple negative status, status of hormone receptor, were obtained from Koboldt *et al*[35]. Copy number alternation status of *MYC* and *PVT1* were called by GISTIC[36] using the SNP6 DNA microarray data from breast carcinoma in TCGA, were obtained from cBioPortal (https://www.cbioportal.org/)[37]. For prostate adenocarcinoma, Gleason scores were obtained from Abeshouse *et al*[38]. For head and neck squamous cell carcinoma, HPV status was obtained from Lawrence *et al*[39]. In renal papillary carcinoma, molecular subtypes were obtained from Linehan *et al*[40].

*International Cancer Genome Consortium – Early-Onset Prostate Cancer data*[41]

Matched RNAseq and whole genome sequencing (WGS) data from 121 tumor samples and 9 adjacent normal samples from 96 patients, the corresponding clinical data including biochemical recurrence (BCR), and Gleason scores were downloaded from an early-onset (treatment age < 55) prostate cancer patient cohort (ICGC-EOPC)[41]. Among these 96 patients, there were 13 with Gleason score = 3+3, 58 with Gleason score = 3+4, 11 with Gleason score = 4+3, 1 with Gleason score = 4+4, 6 with Gleason Score = 4+5, 6 with Gleason score = 5+4, and 1 with Gleason Score = 5+5.

For the ICGC-EOPC dataset, the gene expression read counts from the ultra-deep total RNAseq and relevant clinical data of 121 tumors samples from 96 patients were obtained from Gerhauser *et al*[41]. RNA reads were aligned to the human GRCh37 reference genome using BWA and SAMtools. Uniquely mapped reads were annotated using Ensembl v62. DNA library preparation and WGS was performed on Illumina sequencers[42] with a median insert size of 310 bp (SD = 57 bp) and a median WGS coverage of 61-fold for tumor and 38-fold for germline control samples. WGS data was aligned to the GRCh37 reference genome using BWA-MEM[43] according to Pan Cancer Analysis of Whole Genomes (PCAWG) protocol (https://doi.org/10.1101/161638). DNAseq-based purity and ploidy estimates for 113 samples from 89 patients were determined by Sequenza[44].

*Renal Medullary Carcinoma data*

Matched RNAseq and whole exome sequencing (WES) data of 11 fresh frozen tumor samples from 11 patients were downloaded from Msaouel, P. *et al*[45]. RNAseq data of 7 adjacent normal samples from 7 of the 11 patients were also downloaded from this study. Raw RNAseq reads generated by the HiSeq 2000 sequencer in fastq format were qualified using FastQC (0.10.1) and aligned to the human hg19 reference genome using STAR[46] (version 2.5.2b). Gene expression read counts were generated using featureCounts[47]. The raw WES reads generated by the HiSeq 4000 sequencer in fastq format were also qualified by FastQC and aligned to the human hg19 reference genome using BWA[43] and SAMtools. The Gleason scores of these patients were downloaded from[45].

*Clinical trial prostate cancer data (NCT01946165)*

A total of 27 radical prostatectomy samples were analyzed from 25 patients enrolled in a pre-operative study that assessed the effect of a neo-adjuvant 6-month treatment with Abiraterone Acetate (Abi) plus LHRH agonist versus Abi combined with Enzalutamide (Enza) and LHRH agonist (LHRHa) in patients with newly diagnosed high risk prostate cancer. The study was conducted according to the principles of the Declaration of Helsinki and the International Conference on Harmonization Good Clinical Practice Guideline, and all patients had consented for the processing of their tumor tissue samples in future research projects. The median age at treatment initiation was 60 years (49-73). Gleason scores in diagnostic biopsy samples performed prior to treatment initiation were: Gleason 6 (3+3) in one patient, Gleason score 8 (4+4) in eight patients and Gleason score 8 (5+3) in one patient, Gleason score 9 (4+5) in thirteen patients and Gleason score 9 (5+4) in 1 patient, Gleason score 10 (5+5) in one patient. The median overall pre-treatment PSA was 16.1 ng/mL (0.5-141). Seven patients were treated with 6 cycles of Abi plus LHRHa and 18 patients received 6 cycles of Abi, Enza and LHRHa. All patients except for 3 achieved an undetectable PSA prior to surgery. Two surgical specimens were staged as pT2, three as pT3a, and twenty as pT3b, while 13 patients had nodal disease involvement. The surgical margins were positive in 6 patients and equivocal in 1 patient. Median overall tumor cellularity (E), percentage of tumor cells in the largest tumor surface area, in the prostatectomy specimens was 45% (5-90). Median overall tumor volume(V), estimated by the three-dimensional estimation method, was 2.28 cc (0.48-8.64). Median overall tumor epithelial volume, estimated as E x V, was 0.734 (0.144-5.875)[48,49]. Twenty-two patients demonstrated a PSA relapse in a median of 16 months post-surgery. Bam files of the matched RNAseq and WES data from 27 radical prostatectomy samples were obtained. DNAseq-based purity and ploidy estimates for 27 samples were determined by Sequenza[44] using WES data. For RNAseq data, STAR[46] (version 2.5.2b) and HTseq[50] (version 0.11.1) were used to map reads to the human hg19 reference genome and generate the final raw counts matrix.

##### 2.3.2. TCGA

###### 2.3.2.1. Data processing

To estimate the tumor-specific mRNA proportions ($\pi$) for each sample, we used the two-component mode of DeMixT for 15 TCGA cancer types where sufficient normal reference samples were available (the

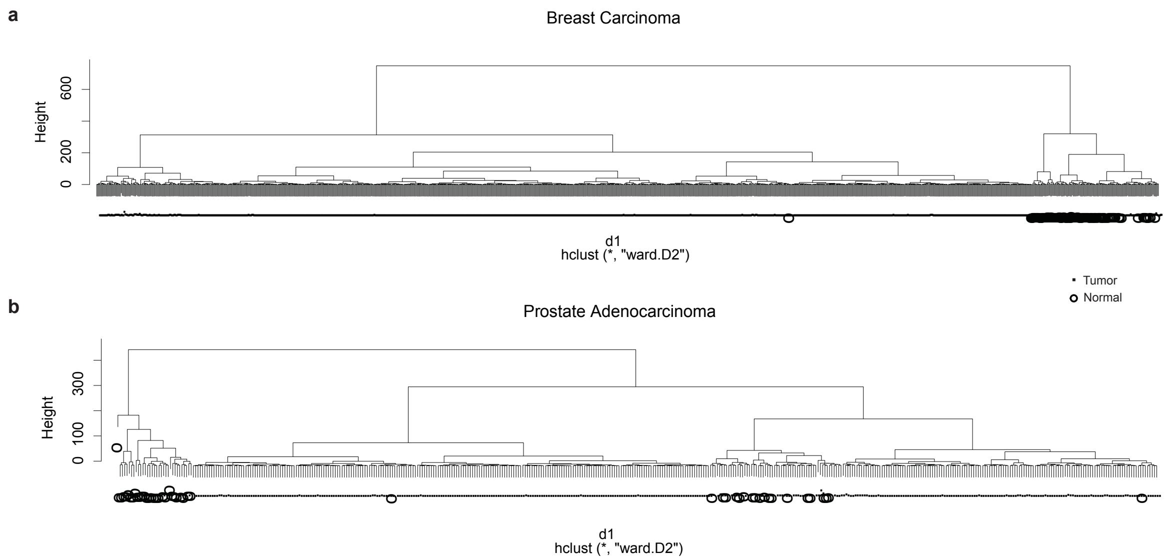

**Supplementary Note Figure 7. Hierarchical clustering of read count data from adjacent normal and mixed tumor samples for breast and prostate adenocarcinoma in TCGA as examples.** Dendrograms of the hierarchical clustering results based on the top 1,000 differentially expressed feature genes for breast carcinoma (**a**) and prostate adenocarcinoma (**b**), respectively. Circles represent adjacent normal samples and dots represent mixed tumor samples.

minimum number of normal samples is seven). For each cancer type, the following quality control was performed on both the tumor and normal samples to remove any suspicious samples. For each gene, we first used the Wilcoxon rank-sum test to test for differential expression between normal and tumor samples. The top 1,000 genes with the smallest *P* values were selected as the feature genes. The first two principal component scores of the feature genes were extracted for hierarchical clustering using Euclidean distance and the Ward method. We separated samples into two groups using the “cutree” function. In general, one cluster contained tumor samples and the other contained normal samples. Any samples that were clustered outside of its general group label, e.g., tumor samples clustered within the normal sample cluster or normal samples in the tumor cluster, were filtered out (**Supplementary Note Figure 7, Supplementary Note Table 6**).

**Supplementary Note Table 6. Summary of sample sizes for 15 TCGA cancer types.**

| Cancer type | Original number of normal samples | Original number of tumor samples | Number of normal samples after quality control | Number of tumor samples after quality control |
|---|---|---|---|---|
| Bladder urothelial carcinoma | 19 | 401 | 17 | 385 |
| Breast carcinoma | 113 | 1074 | 98 | 1032 |
| Colorectal carcinoma | 51 | 633 | 43 | 598 |
| Head & neck squamous cell carcinoma | 44 | 495 | 31 | 494 |
| Renal chromophobe | 24 | 64 | 23 | 64 |
| Renal clear cell carcinoma | 72 | 513 | 66 | 495 |
| Renal papillary carcinoma | 32 | 277 | 26 | 276 |
| Hepatocellular carcinoma | 50 | 362 | 50 | 362 |
| Lung adenocarcinoma | 59 | 455 | 57 | 446 |
| Lung squamous cell carcinoma | 49 | 488 | 48 | 486 |
| Pancreatic adenocarcinoma | 4 | 150 | 7* | 142 |
| Prostate adenocarcinoma | 52 | 406 | 47 | 295 |
| Stomach adenocarcinoma | 32 | 352 | 32 | 299 |
| Thyroid papillary carcinoma | 57 | 498 | 55 | 418 |
| Endometrial carcinoma | 35 | 526 | 26 | 524 |

*Pancreatic adenocarcinoma is the only cancer type with increased normal samples. This is due to the addition of pseudo-normal samples, which are tumor samples of stromal tissue with scant tumor presence.

Scale normalization at the seventy-fifth percentile based on the DSS package[51] was then applied to the post quality-control tumor and normal samples. Next, we applied two criteria to filter out spurious genes. First, we filtered out genes with a zero count in either the mixed tumor or normal samples. Second, we filtered out genes with a large variance ($\widehat{\sigma}_{Ng}^2 > 0.6$) in the normal reference samples. Here, the standard deviation of a gene is calculated as $\widehat{\sigma}_{Ng}^2 = sd(log_2(R_{.g}))$, where $R_{.g}$ is the normalized expression of gene *g* for normal reference samples.

For each cancer type, we applied the "DeMixT_DE" to the quality-controlled expression data together with simulated spike-ins as input data to generate initial tumor-specific mRNA proportions $\pi_0$. We used ASCAT estimated tumor purities as an informed prior to calculate a reasonable number for $S_P$. With other datasets in general, we set $S_P = max(50, 0.3*Sample\ size)$, as the default option of the "DeMixT_GS" function. Results from the TCGA datasets across 15 cancer types were largely consistent with small to moderate changes from the addition of spike-ins (**Supplementary Note Figure 8).**

We then used these $\pi_0's$ as initial values in the profile likelihood calculation on all genes to calculate gene selection scores. We ranked all genes based on their gene selection scores from smallest to largest. Based on a simulation study and observed distributions of gene selection scores in real data, we chose the top 1,500 or 2,500 genes to ensure accuracy in proportion estimation (**Supplementary Note Figure 5a)**. Within each cancer type, we used the spike-ins as benchmarking samples and evaluated the RMSE of the estimated proportions of the spike-ins with either the top 1,500 or top 2,500 genes ($\hat{\pi}_{1500}(Sp)$ and $\hat{\pi}_{2500}(Sp)$). If RMSE($\hat{\pi}_{1500}(Sp)$-0)<RMSE($\hat{\pi}_{2500}(Sp)$-0), we used the results of the top 1,500 genes, *i.e.*, the tumor proportions $\pi = \pi_{1500}$; otherwise, $\pi = \pi_{2500}$. In general, the RMSEs were small (median = 0.02 across 15 cancer types), and the two sets of tumor proportions, $\hat{\pi}_{1500}$ and $\hat{\pi}_{2500}$ , were consistent within each cancer type. We additionally removed samples with extreme estimates of $\pi$, >85% or ranked at the top 2.5 percentile of all samples within each cancer type, to mitigate the remaining underestimation when $\pi$ is close to 1 and control the estimation bias in high values of TmS.

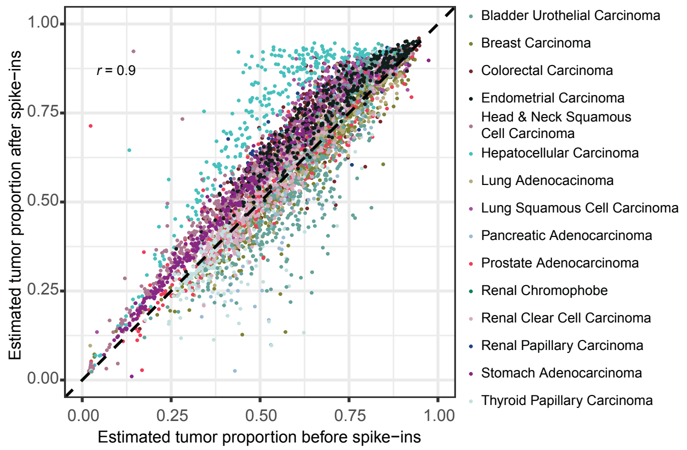

**Supplementary Note Figure 8. Comparison of tumor-specific mRNA proportions with and without spike-ins across 15 TCGA cancer types.** A scatter plot of estimated tumor-specific mRNA proportions using the "DeMixT_GS" method for 4,897 TCGA samples across 15 cancer types with and without spike-ins. The x axis represents the estimated tumor-specific mRNA proportions without spike-ins and the y-axis represents the estimated tumor-specific mRNA proportions with spike-ins.

##### 2.3.2.2. Consensus TmS estimation

We first calculated TmS values for 5,295 TCGA samples with matched tumor-specific mRNA proportions and ABSOLUTE or ASCAT derived tumor purity and ploidy estimates. We then fitted a linear regression model on $log_2$-transformed TmS calculated by ASCAT using $log_2$-transformed TmS calculated by ABSOLUTE as a predictor variable. We removed samples with a Cook's distance ≥ 4/*n* (*n*=5,295) (**Supplementary Fig. 3f-h**), and for the remaining samples, which were the majority, we calculated the final TmS as: $TmS = \sqrt{TmS_{ASCAT} \times TmS_{ABSOLUTE}}$. These TmS estimates were used throughout the paper (**Supplementary Note Table 7**). A CONSORT diagram (**Supplementary Note Figure 9)** demonstrates the sample exclusion for TmS in TCGA**.**

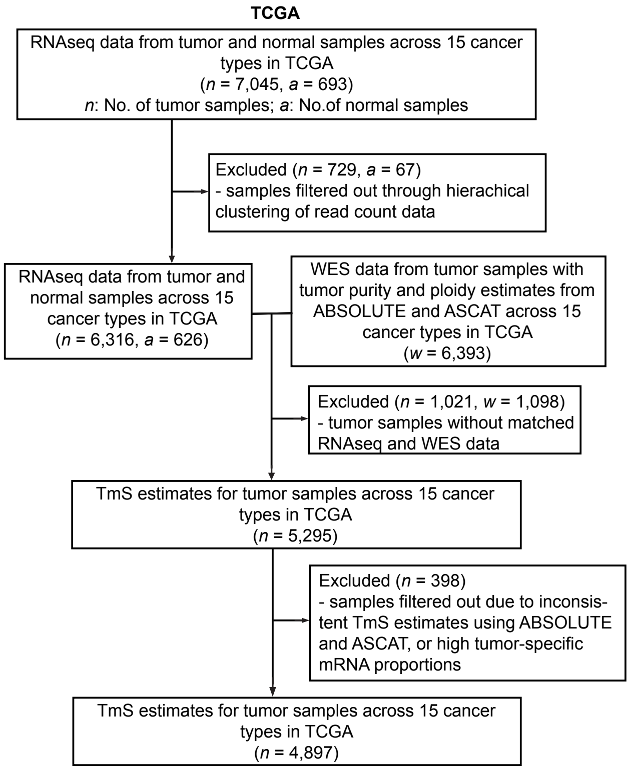

**Supplementary Note Figure 9. CONSORT diagrams for data exclusions in TCGA.**

**Supplementary Note Table 7. Summary of sample sizes for 15 TCGA cancer types before and after consensus TmS estimation.**

| Cancer type | Number of samples before consensus analysis | Number of samples after consensus analysis | Number of samples removed |
|---|---|---|---|
| Bladder urothelial carcinoma | 350 | 328 | 22 |
| Breast carcinoma | 932 | 916 | 16 |
| Colorectal carcinoma | 499 | 490 | 9 |
| Head & neck squamous cell carcinoma | 449 | 443 | 6 |
| Renal chromophobe | 59 | 59 | 0 |
| Renal clear cell carcinoma | 299 | 295 | 4 |
| Renal papillary carcinoma | 192 | 169 | 23 |
| Hepatocellular carcinoma | 333 | 317 | 16 |
| Lung adenocarcinoma | 399 | 395 | 4 |
| Lung squamous cell carcinoma | 440 | 431 | 9 |
| Pancreatic adenocarcinoma | 105 | 101 | 4 |
| Prostate adenocarcinoma | 266 | 259 | 7 |
| Stomach adenocarcinoma | 272 | 265 | 7 |
| Thyroid papillary carcinoma | 297 | 202 | 95 |
| Endometrial carcinoma | 403 | 361 | 42 |

###### 2.3.2.3. Intrinsic tumor signature genes

For each cancer type, we selected the top 1,500 or 2,500 genes based on a gene selection score (Eq.S8), ranked from smallest to largest, as the intrinsic tumor signature gene (**Supplementary Note Figure 10a**). We found the genomic locations of the selected genes covered 22 autosomes and the X chromosome (**Supplementary Note Figure 10b**) across 15 cancer types, which is expected for an unbiased gene set to measure global gene expression. For each cancer type, as well as consistently across 15 cancer types, we find that 54-68% (mean = 62%) of intrinsic tumor signature genes are housekeeping genes[52] or essential genes[53], and 3.5-7.2% (mean = 5.4%) are *MYC* targets genes (**Supplementary Note Figure 10c**). The common pan-cancer essential genes are derived from a total of 147 cancer cell lines and 16,733 genes that were screened independently by both the Sanger and Broad institutes[53].

We conducted gene set enrichment analyses on Hallmark pathways and KEGG pathways[54] using GSEA[55] and g:Profiler[56]. For each cancer type, the genes were ranked according to their gene selection scores from the smallest to the largest. For GSEA, we adopted permutation tests (1,000 times) to generate a normalized enrichment score (NES), the null distribution and an adjusted *P* value for each candidate pathway[55]. g:Profiler detects statistically significantly enriched pathways for the given gene list by implementing hypergeometric tests. For each candidate pathway a nominal *P* value is calculated by the hypergeometric test and adjusted for multiple testing by the BH method. As a result, the minimum NES of significantly enriched Hallmark pathways and KEGG pathways are above 1.74 and 1.70, respectively.

We further evaluated the chromatin accessibility of signature genes using ATAC-seq data TCGA samples[33]. For each sample, peak scores (-log10(p-value)) were scaled by dividing each individual peak score by the sum of all of the peak scores in the given sample divided by 1 million. These scaling values ranged from 1.4 to 67.4 across cancer types. The 75th percentile of normalized peak scores across all peaks within the promoter region were selected for each gene as representative peak scores, and genes with normalized peak scores less than 1 were excluded. A total of 7.1% to 20.4% of genes were excluded across cancer types (**Supplementary Note Figure 10d**). For each sample, we calculated the mean of the peak scores of all signature genes. A null distribution of mean peak scores was generated by calculating means from 1,000 random subsets of genes with the matching number of the signature genes from all genes. *P* values for signature genes were calculated as the percentile of the permuted means being greater than or equal to the observed mean. *P* values were adjusted for multiple testing by the BH method.

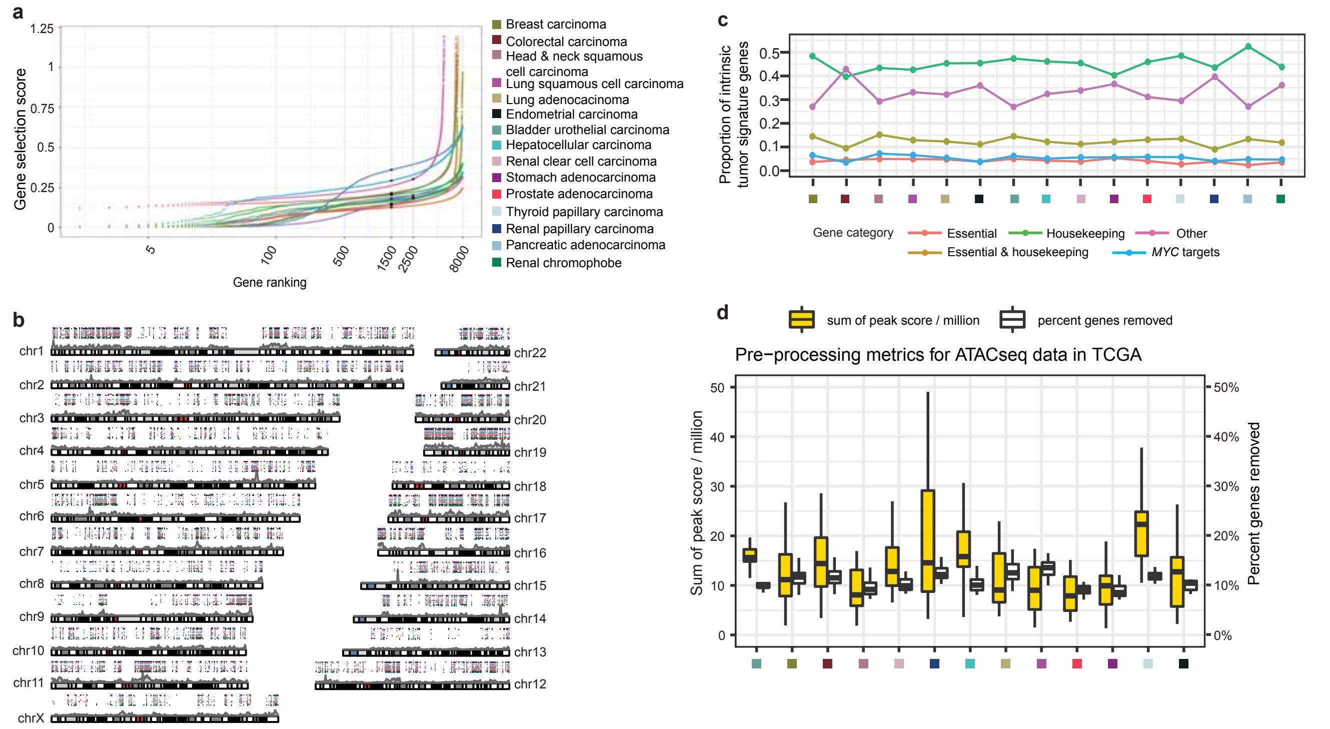

**Supplementary Note Figure 10. Validation of gene selection to represent intrinsic tumor signature across cancer types. a**, Ordered gene selection scores of all genes from low to high across 15 cancer types. The black solid dots represent the cutoffs for gene selection in the corresponding cancer type. **b**, Karyotype plots showing the genomic locations of signature genes for each cancer type. Signature genes are presented as dots colored by cancer type. An overall gene density track is shown in gray shades underneath the dots. The density of signature genes is consistent with the overall gene density. **c**, Proportions of the intrinsic tumor signature gene categories across 15 cancer types. **d**, Boxplots showing sums of peak scores across samples / 1 million (yellow) and percentages of genes removed per sample (white). The sums of peak scores were used as scaling factors to normalize the ATAC-seq data. The genes with normalized scores <1 were removed from downstream analysis. For a,b,c,e, all 15 TCGA cancer types are annotated using colored squares as shown in the legend of a.

#### 3. STATISTICAL ANALYSIS

##### *Association with survival outcomes*

For the TCGA datasets, we used clinical data that passed at least one of the three quality control steps introduced from the TCGA pan-cancer clinical paper[57]. We used two survival outcomes, the overall survival (OS) and progression-free interval (PFI). To ensure sufficient sample size in each category, we combined the pathologic stages into two categories: early stage and advanced stage. The early stage includes Stage I, Stage IA, Stage IB, Stage IC, Stage II, Stage IIA, Stage IIB, and Stage IIC, while the advanced stage consists of Stage III, Stage IIIA, Stage IIIB, Stage IIIC, Stage IV, Stage IVA, Stage IVB, and Stage IVC. With prostate cancer, we used Gleason score (Gleason Score = 7 versus Gleason Score ≥ 8) instead of early and advanced stage. The CONSORT diagram that demonstrates the sample exclusion for survival analysis in TCGA is shown in **Supplementary Note Figure 11.** The CONSORT diagram that demonstrates the sample exclusion in ICGC-EOPC is shown in **Supplementary Note Figure 12.**

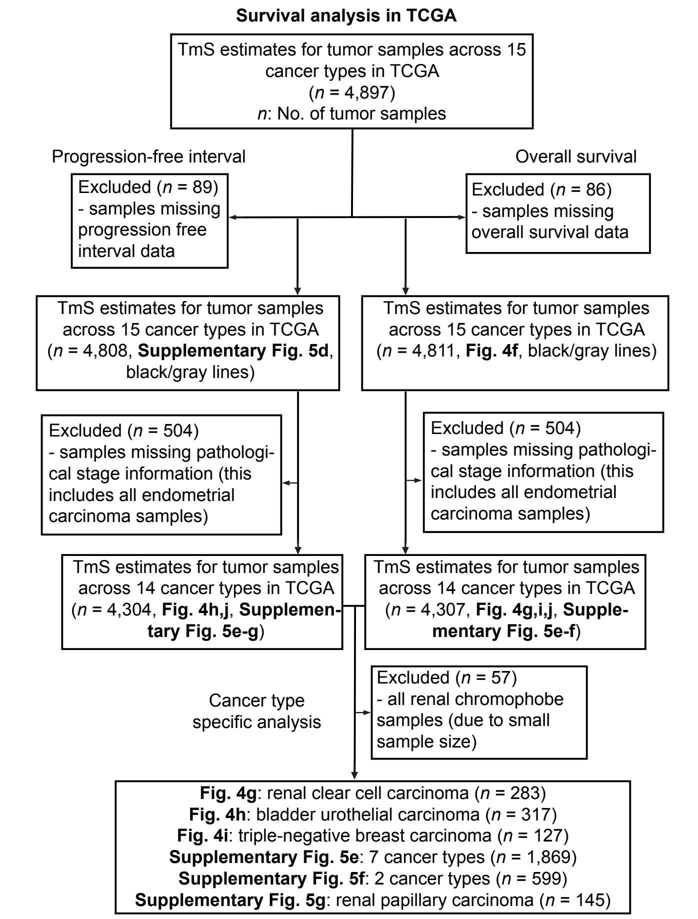

**Supplementary Note Figure 11. CONSORT diagram for data exclusions in TCGA survival analysis.**

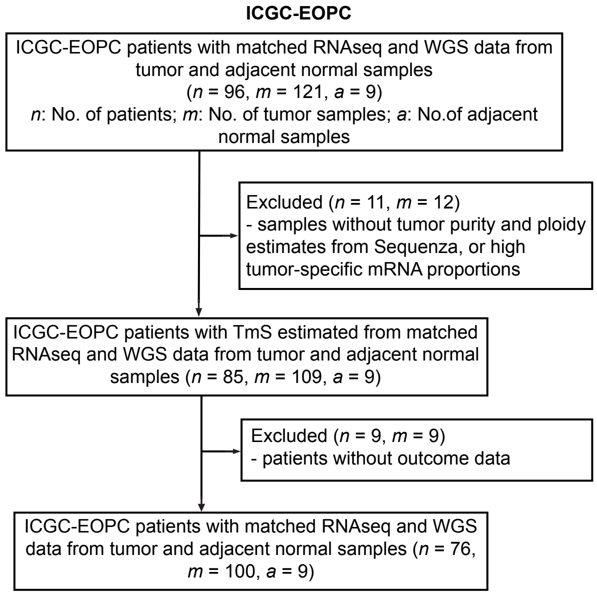

**Supplementary Note Figure 12. CONSORT diagram for data exclusions in ICGC-EOPC.**

Due to the potential nonlinear relationship between TmS and survival outcomes, we used a recursive partitioning survival tree model, rpart[58], to find an optimized TmS cutoff that best differentiates survival outcomes within each of the two stages as defined above in each cancer type. The splitting criteria were Gini index, and the maximum tree depth was set to 2. The TmS cutoffs of early/advanced stage across cancers are shown in **Supplementary Note Table** 8.

**Supplementary Note Table 8. Summary of TmS cutoffs for early/advanced stage across cancers.**

| Cancer type | Overall survival | | Progression-free interval | |
|---|---|---|---|---|
| | Early stage | Advanced stage | Early stage (Gleason score = 7 for prostate cancers) | Advanced stage (Gleason score ≥ 8 for prostate cancers) |
| Pan-Cancer(14 cancer types) | 1.10 | 1.72 | 1.65 | 1.72 |
| Bladder urothelial carcinoma | 0.15 | NA | 0.15 | 0.60 |
| Triple-negative breast carcinoma | 4.11 | 1.80 | 3.02 | 1.80 |
| Colorectal carcinoma | 1.94 | 4.52 | NA | 4.14 |
| Head & neck squamous cell carcinoma (HPV-) | 1.00 | 0.26 | 0.14 | 0.26 |
| Renal chromophobe | 2.21 | 3.95 | 1.79 | 0.74 |
| Renal clear cell carcinoma | 0.54 | 1.78 | 0.33 | 1.67 |
| Renal papillary carcinoma | 0.61 | 0.71 | 0.87 | 0.64 |
| Hepatocellular carcinoma | 0.16 | 1.81 | NA | 0.64 |
| Lung adenocarcinoma | 0.81 | 0.97 | 0.51 | 8.66 |
| Lung squamous cell carcinoma | 6.67 | 2.08 | 5.67 | 6.37 |
| Pancreatic adenocarcinoma | 1.83 | NA | 1.83 | NA |
| Stomach adenocarcinoma | 0.40 | 0.15 | 0.28 | 0.31 |
| Thyroid papillary carcinoma | NA | NA | 0.57 | 1.25 |
| Prostate adenocarcinoma | NA | NA | 0.50 | 0.48 |
| Early-onset prostate cancer (ICGC-EOPC) | NA | NA | 1.25 | 0.84 |

#### 4. HYPOXIA SCORE

Hypoxia scores were generated as described previously[59,60] using the Buffa Signature[61], which has been previously shown to be well-correlated with direct and transcriptional measures of hypoxia[62]. Signature scoring was done on Level 3 mRNA abundance data (2016-01-28 data release) as a single cohort of 7,791 patients to ensure comparability of scores across cancer types. For each gene in the signature, patients were median dichotomized. Patients with RNA abundance above the median were assigned a gene-score of +1, while those with RNA abundance below the median were assigned a gene-score of -1. The gene-scores for all signature genes were summed to generate a per-patient hypoxia-score. High values of this score suggest more hypoxia (lower levels of oxygen), while low values suggest less hypoxia (higher levels of oxygen). We also calculated the Spearman correlation coefficients between TmS, and pentose-phosphate-13/hypoxia/TMB/CIN scores for each cancer type (**Supplementary Note Table 9**).

**Supplementary Note Table 9. Spearman correlation coefficients between TmS and pentose-phosphate-13/hypoxia/TMB/CIN scores across 15 cancer types.**

| Cancer type | Pentose-phosphate-13 scores | Hypoxia score | TMB | CIN |
|---|---|---|---|---|
| Breast carcinoma | 0.33 | 0.65 | 0.35 | 0.46 |
| Lung adenocarcinoma | 0.33 | 0.61 | 0.3 | 0.23 |
| Thyroid papillary carcinoma | 0.22 | 0.44 | 0.066 | 0.1 |
| Pancreatic adenocarcinoma | 0.41 | 0.43 | 0.082 | 0.1 |
| Renal clear cell carcinoma | 0.46 | 0.41 | 0.17 | 0.35 |
| Lung squamous cell carcinoma | 0.25 | 0.38 | 0.04 | 0.051 |
| Bladder urothelial carcinoma | 0.18 | 0.35 | 0.21 | 0.24 |
| Renal papillary carcinoma | 0.38 | 0.35 | -0.21 | 0.18 |
| Colorectal carcinoma | 0.15 | 0.3 | -0.021 | 0.056 |
| Prostate adenocarcinoma | 0.043 | 0.26 | 0.072 | 0.3 |
| Endometrial carcinoma | 0.24 | 0.25 | -0.062 | 0.16 |
| Hepatocellular carcinoma | -0.039 | 0.2 | -0.066 | -0.095 |
| Head & neck squamous cell carcinoma (HPV+) | 0.22 | 0.17 | 0.32 | 0.27 |
| Head & neck squamous cell carcinoma (HPV-) | 0.17 | -0.08 | -0.017 | -0.081 |
| Stomach adenocarcinoma | 0.24 | NA | -0.073 | -0.15 |
| Renal Chromophobe | 0.43 | NA | 0.13 | 0.23 |
